# Genetically encoded reporters of actin filament organization in living cells and tissues

**DOI:** 10.1101/2024.04.26.591305

**Authors:** Carla Silva Martins, François Iv, Shashi Kumar Suman, Thomas C. Panagiotou, Clara Sidor, María Ruso-López, Camille N. Plancke, Shizue Omi, Rebecca Pagès, Maxime Gomes, Alexander Llewellyn, Sourish Reddy Bandi, Laurie Ramond, Federica Arbizzani, Caio Vaz Rimoli, Frank Schnorrer, François Robin, Andrew Wilde, Loïc LeGoff, Jean-Denis Pedelacq, Antoine Jégou, Stéphanie Cabantous, Sergio A. Rincon, Cristel Chandre, Sophie Brasselet, Manos Mavrakis

## Abstract

The cytoskeletal protein actin is crucial for cell shape and integrity throughout eukaryotes. Actin filaments perform essential biological functions, including muscle contraction, cell division and tissue morphogenesis. These diverse activities are achieved through the ability of actin filaments to be arranged into precise architectures. Much progress has been made in defining the proteome of the actin cytoskeleton, but a detailed appreciation of the dynamic organizational state of the actin filaments themselves has been hindered by available tools. Fluorescence polarization microscopy is uniquely placed for measuring actin filament organization by exploiting the sensitivity of polarized light excitation to the orientation of fluorophores attached to actin filaments. By engineering fusions of five widely used actin localization reporters to fluorescent proteins with constrained mobility, we have succeeded in developing genetically-encoded, green- and red-fluorescent-protein-based reporters for non-invasive, quantitative measurements of actin filament organization in living cells and tissues by fluorescence polarization microscopy.

## INTRODUCTION

Actin is one of the most abundant and conserved proteins throughout eukaryotes, including structural homologs in bacteria. Actin monomers polymerize through non-covalent association to generate filaments. These filaments perform a wide range of essential biological functions, such as muscle contraction, cell division, cell adhesion, cell motility, tissue morphogenesis and intracellular pathogen movement ^1^. These various activities are achieved through dozens of actin binding proteins which configure actin into a diverse set of organization states ^2^. The precise geometrical organization of actin filament assemblies, i.e. how actin filaments are physically oriented in space and to what extent they are aligned to each other (collectively referred to as filament *organization*, hereafter), is crucial to biological function ^3–7^. Typically, multiple actin-binding proteins populate actin filament (F-actin) assemblies in cells, and predicting actin filament organization and its effect on function remains elusive.

Cryo-electron tomography emerges as a very powerful method to visualize the nanoscale organization of actin filaments in cells ^8^, but it is not applicable to living samples. The most widely used approach for imaging actin filaments in living cells uses fluorescence microscopy and either one of the following three actin filament-binding probes: (1) small (∼1 kDa) organic fluorescent dye conjugates of the F-actin-binding drugs phalloidin ^9,10^ and jasplakinolide ^11^; such examples are AlexaFluor488 (AF488)-phalloidin and silicon rhodamine (SiR)-jasplakinolide, the latter known with the misleading name “SiR-actin”, (2) GFP fusions to G-actin ^12–16^, and (3) GFP fusions to actin-binding peptides or protein domains (for example, the actin-binding domain of moesin ^17^). The intensity and distribution of the fluorescent pixels in the images inform us on the relative localization and levels of filamentous actin within the cell. However, how actin filaments are organized at a given image pixel, within the optical resolution of the microscope (typically ∼200 nm), cannot be deduced from the fluorescence intensity alone, nor from the pattern of fluorescent pixel distribution.

Fluorescence polarization microscopy (hereafter, polarimetry) is ideally placed for measuring actin filament organization in living cells by exploiting the sensitivity of polarized light excitation to the orientation of fluorophores attached to actin filaments. Fluorescence is maximized when polarized light is aligned with the excitation dipoles of the fluorophores ^18^. Thus, an ensemble of fluorophores will be excited more efficiently in preferred directions depending on the organization of the fluorophores. By measuring the modulation of fluorescence induced by the rotation of light polarization in the sample plane, we can extract independently two angles per image pixel: the mean orientation of the fluorophores within the focal volume (Figure 1A, angle rho, ρ) and the in-plane projection of the angle explored by these fluorophores, (Figure 1A, angle psi, ψ) ^19^. If the fluorophores are linked to actin filaments in a constrained manner, the measured parameters reflect directly the molecular-scale organization of the labeled filaments in living cells at each pixel location, with ρ their mean orientation, and ψ their degree of alignment; the higher the filament alignment, the lower the ψ.

**Figure 1.**
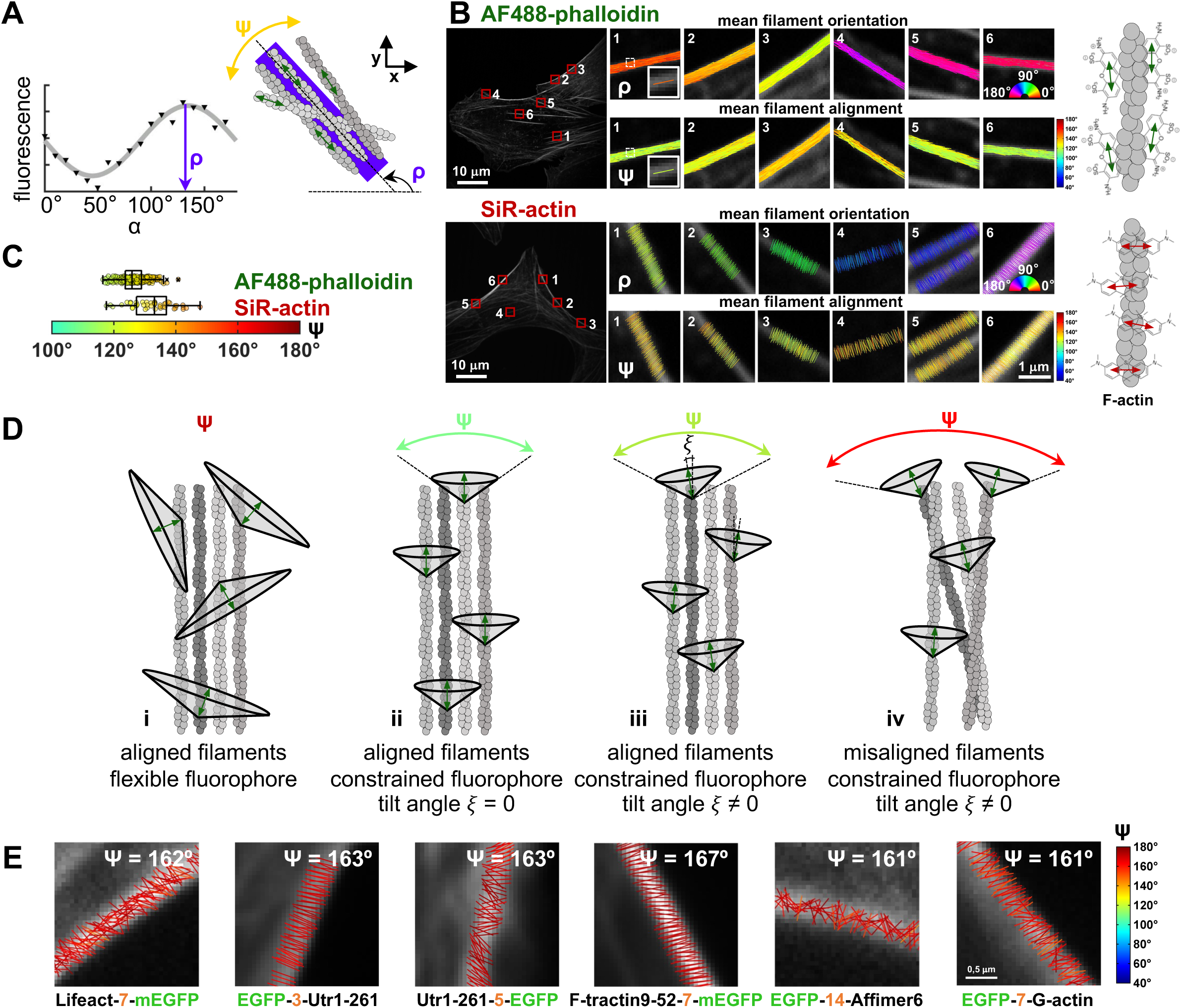
**Measuring actin filament organization in cells with polarimetry** (A) Left, example of the polarization response of a sample at a given pixel of the image as obtained from a recorded polarimetry stack. The polarimetry stack is made of 18 polarized fluorescence images acquired using an incident linear polarization angle, α, varying from 0° to 170° with steps of 10°. Raw datapoints are shown as triangles and the theoretical fitting curve as a solid line. Right, schematic of a hypothetical organization of four fluorescently-labeled actin filaments in the confocal volume of the measured pixel, with the different orientations of the fluorophore dipoles shown by green double-headed arrows. The fluorophore dipoles are parallel to the actin filament axis in this example. The angle ρ corresponds to the mean orientation of all dipoles and thus the average orientation of actin filaments in the confocal volume. The ρ value is represented with a purple stick whose orientation and color depict the mean filament orientation in the pixel (see colorbar in (B)). The angle ψ corresponds to the angular aperture explored by all dipoles and is thus a readout of the average filament alignment in the confocal volume. (B) Representative examples of polarimetry measurements of actin filament organization in fixed U2OS cells labeled with AF488-phalloidin (top) or SiR-actin (bottom). The zoomed-out images on the left are summed intensity images of the respective polarimetry stacks. Insets on the right show zoom-ins of selected regions of interest (red outlined boxes) containing actin stress fibers (SFs) in different orientations, with the measured ρ and ψ angles per pixel. The angles ρ (top insets) are represented as ρ stick maps (”orientation maps”), with a stick per pixel whose orientation and color depict the mean filament orientation in the pixel. The values of ρ, from 0° to 180°, are color-coded according to the colorbar. The angles ψ (bottom insets) are represented as ψ stick maps (”organization maps”), with a stick per pixel whose orientation depicts the mean filament orientation (ρ) and whose color corresponds to the mean filament alignment (ψ) in the pixel. The values of ψ, from 40° to 180°, are color-coded according to the colorbar. (C) Box plots depicting the distribution of ψ angle measurements on SFs as shown in (B). The data points, color-coded according to the ψ colorbar, are plotted on top of the respective box plots. On each box, the central mark indicates the median, and the left and right edges of the box indicate the 25th and 75th percentiles, respectively. The whiskers extend to the most extreme data points not considered outliers, and the outliers are plotted individually using the ’x’ symbol. The number of measurements in each box plot is n = 258 and 45 for AF488-phalloidin and SiR-actin, respectively. The respective median values are 126° and 133°. (D) Schematics showing the dependence of measured ψ angles on the underlying actin filament organization, the mobility of the fluorophore and the tilt angle, ξ, of the fluorophore with respect to the axis of the actin filament. ψ is color-coded as in (B). The mean filament orientation, ρ, is the same in all cases. Flexible fluorophores will lead to very high (>160°) ψ values and thus an overestimation of disorder even for highly aligned actin filaments (i). Constrained fluorophores allow us to detect changes in actin filament organization (ii and iii *vs.* iv). (E) Representative ψ stick maps on SFs from measurements in live cells expressing widely used GFP fusions of actin-binding peptides or domains, or G-actin itself. The number shown in orange corresponds to the number of amino acid residues of the linker between the GFP and the actin-binding moiety. Mean ψ values are shown.

Fluorophore conjugates to phalloidin and jasplakinolide have been used successfully for polarimetry in fixed cells ^20–24^. However, their use in living cells is far from ideal given that these drugs stabilize actin filaments ^25–28^. Introducing these drugs into tissues is also experimentally challenging, while controlling their intracellular concentration spatiotemporally is practically impossible, rendering their use in living tissues very limited. The goal of our study is to extend the potential of polarimetry to living cells and tissues by generating genetically-encoded, fluorescent protein-based reporters for live-cell measurements of actin filament organization. To this end, we tailor GFP fusions to widely used F-actin localization reporters by constraining the mobility of the GFP in order to render them usable for organization measurements by polarimetry. We use stress fibers in cultured mammalian cells as a model system of known F-actin organization to identify constrained GFP fusions that report faithfully the orientation and alignment of actin filaments in live cells. We further validate the functionality and use of the reporters in three genetically tractable *in vivo* model systems, the fission yeast *Schizosaccharomyces pombe*, the nematode *Caenorhabditis elegans* and the fruit fly *Drosophila melanogaster*.

## RESULTS

### Polarimetry allows measurements of actin filament organization in cells

For the sake of illustrating how we display and interpret polarimetry measurements in cells throughout our study, Figure 1B shows an example of polarimetry in fixed mammalian cells stained with AF488-phalloidin or SiR-jasplakinolide. We focus on the analysis of pixels containing stress fibers (SFs), which electron microscopy (EM) has shown to form bundles of actin filaments highly parallel to each other ^29,30^. The angles ρ and ψ are represented as orientation and organization maps, respectively. The distribution of ψ angles for multiple SFs in tens of cells is shown in Figure 1C. The analysis reveals three key findings. First, AF488-phalloidin and SiR-jasplakinolide have fluorophore dipoles that are, on the average, parallel and perpendicular to actin filaments, respectively, in line with previous reports ^20–22,24,31–33^. Second, actin filaments are oriented along the axis of the SFs: SFs oriented differently in the cell show distinct ρ stick colors according to their precise orientation, allowing for an easy visual tracking of changes in mean filament orientation (Figure 1B). Third, actin filaments within SFs are highly aligned. Single AF488 fluorophores, in their conjugates with phalloidin, wobble by ∼90-100° with a tilt angle of ∼20° off the actin filament axis ^31,32^ thus measured ψ of ∼120-130° (Figure 1C and 1D,iii) reflect that the contained filaments at the image pixels of SFs are highly parallel to each other.

The visual inspection of the color-coded ψ maps reveals that all SFs share very similar filament organization despite their different orientations (Figure 1B). Given the precision of only a few degrees for ρ and ψ angle measurements ^34^, the very narrow distribution of ρ and ψ angles within a region of interest in a given SF (for example, standard deviation=3-5° for ROI5 of AF488-phalloidin, Figure 1B) further reveals a very homogeneous population of orientations and aperture angles explored at an image pixel, in full line with what one expects from EM. This said, the reason why such measurements are possible in these examples is because AF488 and SiR are sufficiently constrained in their conjugates with phalloidin and jasplakinolide, respectively.

### Widely used genetically encoded actin localization reporters are not suitable for organization measurements with polarimetry

To assess the usability of available genetically-encoded F-actin localization reporters for polarimetry, we measured actin filament orientation and alignment on stress fibers (SFs) of live U2OS cells expressing either of five widely used F-actin-binding EGFP fusions: the F-actin binding peptides Lifeact ^35^ and F-tractin9-52 ^36^, the actin-binding domain of human Utrophin Utr1-261 ^37^, the synthetic actin-binding Affimer, Affimer6 ^38^, and human non-muscle beta G-actin ^14^. The measured ψ angles on SFs were very high (>160°) in all cases (Figure 1E). Given that actin filaments in stress fibers are highly aligned to each other, the high ψ values cannot result from disordered actin filaments, but they rather reflect the high rotational mobility of EGFP in the respective fusions (Figure 1D,i), rendering the latter not suitable for organization measurements.

### Design of GFP-based reporters with constrained GFP mobility

To tailor the available F-actin localization reporters for organization measurements, we engineered fusions with constrained GFP mobility. We reasoned that there are three main sources that contribute to the flexibility of the GFP. The first source is the presence of an amino acid linker between the actin-binding moiety (actin-binding peptide or protein domain, or G-actin) and the fused GFP. The second source is the flexibility of the terminus of the actin-binding moiety to which GFP is fused, and the third source of flexibility are the N- and C-termini of GFP themselves (Figure S1A). To constrain GFP mobility, we embarked on a screen of GFP fusions to widely used actin-binding peptides and protein domains (ABDs), namely Lifeact, Utrophin Utr1-261, F-tractin9-52, and Affimer6, and generated terminal and circularly permuted GFP fusions (Figure 2A) without linker sequences, with shortened GFP termini and shortened termini of the ABD (Figures S2-S4). We considered the use of monomeric superfolder GFP (msfGFP) as the best choice for GFP fusion engineering (Figure S1B-D; see Data S1 for a comprehensive discussion of the screen design and choice of GFP).

**Figure 2.**
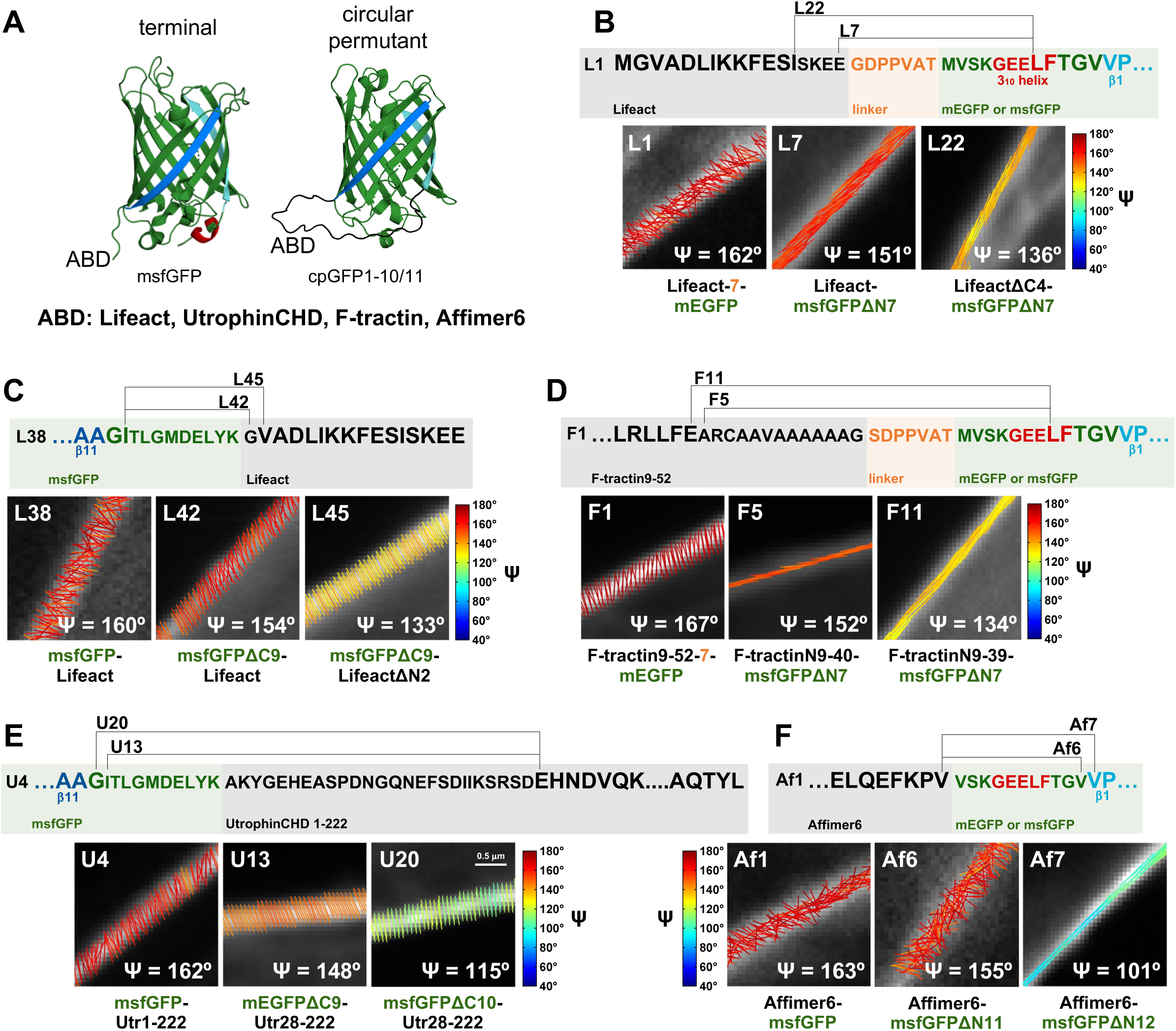
**Engineering of Lifeact-, F-tractin, Utrophin- and Affimer6-based actin filament organization reporters for live-cell polarimetry** (A) Designs used in this study to immobilize genetically-encoded fluorophore fusions to actin-binding peptides or protein domains (ABDs). ABDs were fused to the N- or C-terminus of msfGFP (left) or placed in-between the N- and C-terminus of GFP using a circularly permuted GFP (cpGFP) (right). (B-F) Engineering of Lifeact- (B,C), F-tractin- (D), Utrophin- (E) and Affimer6-based (F) actin filament organization reporters. Representative designs for constraining GFP mobility are illustrated for selected fusions, including for the best performing reporters (L22, L45, F11, U20, Af7, see Table 1). The full screens are shown in Figures S2, S3 and S4. The top panels show the primary sequence of an unconstrained fusion using its designated number (for example, L1 for panel B). Secondary structure elements of GFP are color-coded as in Fig.S1A. Only the region of the primary sequence that is modified is shown. The primary sequences of constrained GFP fusions (for example, L7 and L22 for panel B) are indicated using brackets connecting the fused residues. Constraining GFP mobility involves removing linker sequences and additionally shortening the terminus of GFP and/or the terminus of the ABD. Bottom panels show representative ψ stick maps on SFs from measurements in live U2OS cells expressing the respective fusions. Mean ψ values are indicated: the selected images correspond to median ψ values of the respective distributions shown in Figures S2, S3 and S4. The full names of the fusions are mentioned below the ψ stick maps: the number shown in orange, if any, corresponds to the number of amino acid residues of the linker between the GFP and the ABD.

**Table 1.** Overview of the best performing actin filament organization reporters for live-cell polarimetry. The table summarizes key features of the best performing reporters, including their binding affinity to actin filaments (for the reporters characterized *in vivo*), their dipole orientation with respect to the actin filament axis, reference ψ values from measurements on different F-actin structures, the different functional readouts used in this study, as well as their potentially perturbative character.

| actin filament organization reporter | binding affinity to actin filaments (This paper) | dipole orientation with respect to actin filaments | reference $\psi$ values | functional readouts (This paper) | perturbative character # (This paper) |
| --- | --- | --- | --- | --- | --- |
| **actin-binding domains, green FP fusions** |  |  |  |  |  |
| Affimer6: Af7 | Kd = $0.6 \pm 0.2$ μM | parallel | 100° (SF)<br>90° (CR)<br>75° (FM) | human cell culture<sup>1</sup>;<br>*in vivo*: fission yeast<sup>2</sup>,<br>*C.elegans*<sup>3</sup>, *Drosophila*<sup>4</sup> | fission yeast<sup>2</sup>: slower maturation and constriction of cytokinetic ring; promotes growth in the presence of LatA<br>*Drosophila*<sup>4</sup>: impaired flight muscle development and flightless when expressed throughout muscle development |
| Lifeact: L22 | Kd = $11.3 \pm 3.6$ μM | parallel | 135° (SF)<br>120° (CR)<br>115° (FM) | human cell culture<sup>1</sup>;<br>*in vivo*: fission yeast<sup>2</sup>,<br>*C.elegans*<sup>3</sup>, *Drosophila*<sup>4</sup> | fission yeast<sup>2</sup>: slower assembly of cytokinetic ring; impairs growth in the presence of CK666; promotes growth in the presence of LatA<br>*Drosophila*<sup>4</sup>: weak flier when expressed throughout muscle development |
| Lifeact: L45 | Kd = $4.5 \pm 3.5$ μM | perpendicular | 135° (SF)<br>135° (FM) | human cell culture<sup>1</sup>;<br>*in vivo*: fission yeast<sup>2</sup>,<br>*C.elegans*<sup>3</sup>, *Drosophila*<sup>4</sup> | |
| UtrophinCHD: U20 | Kd = $13.2 \pm 7.4$ μM | perpendicular | 115° (SF) | human cell culture<sup>1</sup>;<br>*in vivo*: fission yeast<sup>2</sup>,<br>*Drosophila*<sup>4</sup> | |
| F-tractin: F11 | nd* | parallel | 135° (SF) | human cell culture<sup>1</sup> |  |
| **G-actin, green FP fusions** |  |  |  |  |  |
| G-actin: A4 |  | perpendicular | 145° (SF) | human cell culture<sup>1</sup> |  |
| G-actin: A18 |  | parallel | 150° (SF) | human cell culture<sup>5</sup> |  |
| **actin-binding domains, red FP fusions** |  |  |  |  |  |
| Affimer6: Af30 | nd* | parallel | 110° (SF) | human cell culture<sup>1</sup> |  |
| Lifeact: L81 | nd* | perpendicular | 110° (SF) | human cell culture<sup>1</sup> |  |

<sup>1</sup> localization to different types of SFs; <sup>2</sup> timing of stages of cytokinesis, cell growth under sensitized conditions (CK666, LatA), genetic interactions with profilin mutant; <sup>3</sup> embryonic lethality, embryonic elongation; <sup>4</sup> adult wing growth, flight tests ; <sup>5</sup> actin isoform-specific localization, actin isoform-specific rescue of cell division defects, and compensatory actin isoform expression for the respective isoform-specific intramolecular fusion A8. \* not determined. SF, stress fibers; CR, cytokinetic ring in fission yeast; FM, *Drosophila* flight muscle

Similarly to the widely used localization reporters (Figure 1E), we expressed fusions transiently in U2OS cells and made polarimetry measurements on SFs. Given that SFs comprise highly aligned actin filaments, and if GFP mobility is sufficiently constrained, we expect to measure mean fluorophore dipole orientations (ρ angle values) that are either parallel or perpendicular to the SF axis. The smaller the measured ψ angles are on SFs, the more constrained will be the GFP mobility of the respective fusion and thus the larger the range of changes in F-actin organization that we will be able to detect. Data S1 and Figures S2-S4 show and discuss the results of the screening of all constructs.

Table 1 and Figure 2B-F summarize the best performing fusions in terms of constrained GFP mobility, namely L22 (LifeactΔC4-msfGFPΔN7), L45 (msfGFPΔC9-LifeactΔN2), F11 (F-tractinN9-39-msfGFPΔN7), U20 (msfGFPΔC10-Utr28-222) and Af7 (Affimer6-msfGFPΔN12). Comparison of their localization to different F-actin populations, notably different types of SFs, in U2OS cells did not show any significant differences. At the low-expression levels we used and during their transient expression in cells, we did not detect any evident signs of perturbation in terms of SF biogenesis, maintenance or organization. However, given that different actin-binding probes have different affinities for F-actin, and that cells and tissues express tens of actin-binding proteins to regulate the dynamics, localization and specific geometries of actin assemblies related to specific functions, we expected that these same probes behave differently in different cellular contexts. Thus, to gain deeper insights into the functionality of our organization reporters, we chose to test them additionally in a physiological context in the fission yeast *S. pombe*, the nematode *C. elegans* and the fruit fly *Drosophila melanogaster*.

### Actin filament organization reporters in fission yeast

We generated transgenic fission yeast expressing L22, L45, U20 and Af7 (Table 1), and included the respective constructs with unconstrained GFP for comparison, i.e. L2, L38, U7 and Af1, respectively. Cells expressing these reporters under the control of the promoter of the actin cytoskeleton regulator Cdc42, looked healthy, with no signs of vacuolation, and were rod-shaped, suggesting the absence of major polarity defects. We started by comparing the localization of the reporters in populations of actively dividing yeast cells (Figures 3A and S5A-C). Actin in fission yeast is found in three distinct structures: actin patches, actin cables and the cytokinetic actomyosin ring (Figure S5A). Although all reporters labeled all three structures, there were dramatic differences in their respective enrichments, as assessed by the differences in labeling intensities of the different structures (Figures 3A and S5A), as well as in the numbers of patches and cables labeled by each reporter (quantification in Figures 3A and S5A-C). Patches and, to a lesser extent, rings, were labeled well with Lifeact constructs, whereas cables and rings, and, to a lesser extent, patches, were most prominent with Affimer6 constructs (Figures 3A and S5A-C). Utrophin labeling seemed altogether very inefficient. We note that the total amount of patches detected were lower than those reported using fimbrin fused to mEGFP as a marker (Berro and Pollard, 2014). We attribute these striking differences in localization to differences in the affinities of the reporters for the different actin filament assembly geometries and turnover, or due to possible competition with distinct actin-binding proteins in the respective structures.

**Figure 3.**
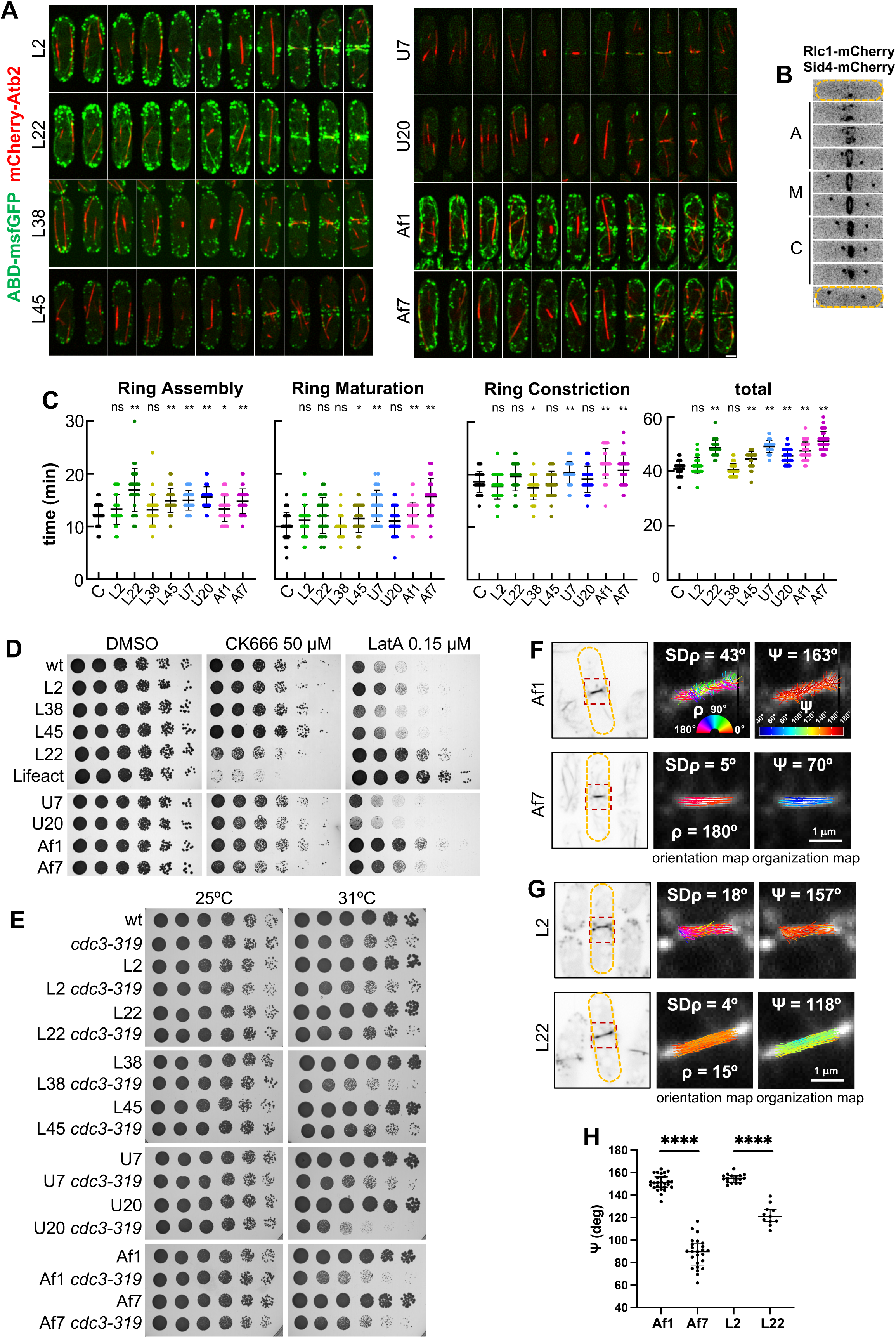
**Polarimetry measurements of actin filament organization in live dividing fission yeast expressing selected reporters** (A) Time-lapse maximum intensity projection images of fission yeast cells co-expressing the tubulin marker mCherry-Atb2 and selected actin organization reporters. Scale bar, 4 µm. (B) Time-lapse maximum intensity projection images of a fission yeast cell (orange dash outline) expressing an acto-myosin ring marker (Rlc1-mCherry) and a spindle pole body marker (Sid4-mCherry) to monitor major cytokinetic events. “A” represents the cytokinetic ring assembly stage, “M”, the cytokinetic ring maturation stage and “C”, the cytokinetic ring constriction stage. Scale bar, 2 µm. (C) Quantification of the time taken for ring assembly completion, ring maturation, ring constriction, and the total time for cytokinesis completion in fission yeast strains expressing each actin reporter and the cytokinetic markers. As a control (”C”), a strain expressing only the cytokinetic markers was used. Scatter plots show means ± SD. The number of cells for each strain is, from left to right: 41, 31, 29, 31, 31, 33, 29, 36, 33. The mean measured times for each strain are, from left to right: 12, 13, 17, 13, 15, 15, 16, 13, 15 min for ring assembly; 10, 11, 12, 10, 11, 14, 11, 12, 16 min for ring maturation; 19, 18, 20, 17, 18, 20, 19, 22, 21 min for ring constriction; and 41, 42, 49, 41, 45, 49, 46, 48, 51 min for total cytokinesis. A t-test was applied to evaluate statistical differences between each strain and the control; ns=not significant, P>0.05; * 0.05>P>0.03; ** P<0.03. (D) Serial dilution assay showing the sensitivity of the fission yeast strains expressing the corresponding actin reporter to CK666, LatA, and DMSO (vehicle control). As controls, a strain expressing Lifeact under the control of the actin promoter (’Lifeact’) and a wild-type strain (’wt’) were included in the assay. (E) Serial dilution assay showing the genetic interaction between the profilin mutant *cdc3-319* and the expression of the different actin reporters. (F-H) Polarimetry measurements of actin filament organization in the cytokinetic ring of living dividing fission yeast cells expressing Affimer6-based (F) and Lifeact-based (G) reporters. Representative measurements are shown for fusions with unconstrained GFPs (Af1, L2) and constrained GFPs (Af7, L22). Left panels in (F), (G) show inverted grayscale summed intensity images of polarimetry stacks for the respective dividing yeast cells (orange dash outlines). ρ and ψ stick maps of actin organization in the cytokinetic ring (red dash box in left panels) are shown in the middle and right panels, respectively. The standard deviation of ρ values (SDρ), mean ρ and ψ values are shown for each map. Scatter plots in (H) show the quantification of ψ angle distributions for each reporter. Scatter plots show medians with interquartile range. The number of cells for each strain is, from left to right: 26, 20, 17, 9. The respective median ψ values are 151, 90, 155, 121°. Statistical significance was obtained using a non-parametric Mann-Whitney test; **** P<0.0001. See also Figure S5A-F.

To assess the functionality of the reporters, we examined their effect in three different contexts: the timing of the different stages of cytokinesis (Figure 3B,C), the growth of cells in the presence of the Arp2/3 complex inhibitor CK666 and the G-actin sequestering drug latrunculin A (LatA) (Figure 3D), and finally genetic interactions with a profilin mutant (Figure 3E). Quantification of the timing of cytokinesis stages showed that cytokinesis proceeded overall similarly for all reporters with respect to a control strain expressing only cytokinetic markers, with cells completing cytokinesis within ∼40-50 minutes in all cases (Figure 3C). We observed the most important delays for the Affimer6 constructs, which showed increased maturation and constriction times, and for some of the Lifeact constructs, notably L22, which took longer for the assembly (Figure 3C).

The dilution assays to assess cell growth in the presence of CK666 and LatA led to three observations. First, there was no difference in cell growth upon expression of the reporters with DMSO as a vehicle control, reflecting their nonperturbative character. Second, cell growth in the presence of CK666 was slightly impaired upon expression of L22 and the effect was even more significant for the original Lifeact construct, which is expressed at a higher level, from the actin promoter ^39,40^. Third, the inability of cells to grow in the presence of LatA was significantly reversed for the Affimer6 constructs, as well as the original Lifeact construct and L22 (Figure 3D). The main effect of CK666 is on actin patches, for the formation of which Arp2/3 is essential. The CK666 results thus suggest that the original Lifeact and, to a lesser extent, L22 may interfere with branched actin formation, or, alternatively, that they may affect cofilin-mediated actin filament severing (Figure S5D), as previously shown in vitro ^41^. LatA, on the other hand, at the low concentration used, impacts primarily the formation of cables, which are not essential for growth, and of cytokinetic rings which are essential for cell division. The LatA results thus reveal a stabilization effect on the rings upon expression of the Affimer6 constructs, the original Lifeact construct and L22. The increased ring maturation and constriction times for Affimer6 and increased ring assembly time for L22 could well be explained by the same mechanisms revealed from the drug assays. Finally, dilution assays in the presence of a thermosensitive profilin mutant did not show any significant difference for the reporters, suggesting that they do not interfere with actin nucleation and polymerization *per se* (Figure 3E).

As an example of the applicability of the reporters *in vivo*, we chose to measure actin filament organization in the constricting cytokinetic ring of live dividing fission yeast cells. We focused on the Affimer6 constructs Af1 and Af7 and the Lifeact constructs L2 and L22 which were most efficient in labeling the cytokinetic ring (Figures 3A and S5A). The unconstrained GFP fusions Af1 and L2 displayed very high ψ values, whereas Af7 and L22 fusions bearing constrained GFP showed statistically significantly lower ψ values (Figure 3F-H). The measured values were comparable to the ones on SFs, suggesting highly aligned actin filaments, with filament alignment persisting at least during the initial stages of ring constriction (Figure S5E). Considering that Af7 and L22 dipoles are parallel to actin filaments (Figure 2B,F), the quantification of ρ angle distribution with respect to the division plane showed that actin filaments are parallel to the constricting ring axis (Figure S5F). The standard deviation of the distributions are of the order of a few degrees (Figure S5F) which is close to the measurement noise ^34^. This distribution is fully in line with the actin filament angle-to-membrane distribution from electron cryotomography of dividing fission yeast: nearly all filaments were shown to make small angles with the membrane, with the peak at 2°, an average of 7.8°, and the majority falling below 20°, revealing actin filaments running nearly parallel to each other and to the membrane ^42^.

### Actin filament organization reporters in *C. elegans*

Next, we sought to test selected reporters in the context of animal morphogenesis, notably *C. elegans* embryonic elongation (Figure 4A). To this end, we generated transgenic *C. elegans* expressing L22, L45 and Af7 under the control of an epidermal promoter (Figure 4B). To assess their functionality, we quantified embryonic lethality by scoring egg hatching in comparison with wild-type embryos and embryos expressing the original Lifeact-GFP reporter. The measured lethality was comparable to wild-type embryos and lower than the one of Lifeact-GFP expressing embryos (Figure 4C). In a second functionality test, we filmed elongating embryos and quantified embryonic length until hatching. Embryonic elongation proceeded similarly for all strains, with Lifeact-GFP, L22 and L45 showing 95-98% elongation and Af7 90% elongation compared to wild-type (Figure 4D), confirming their minimally perturbative character in this process.

**Figure 4.**
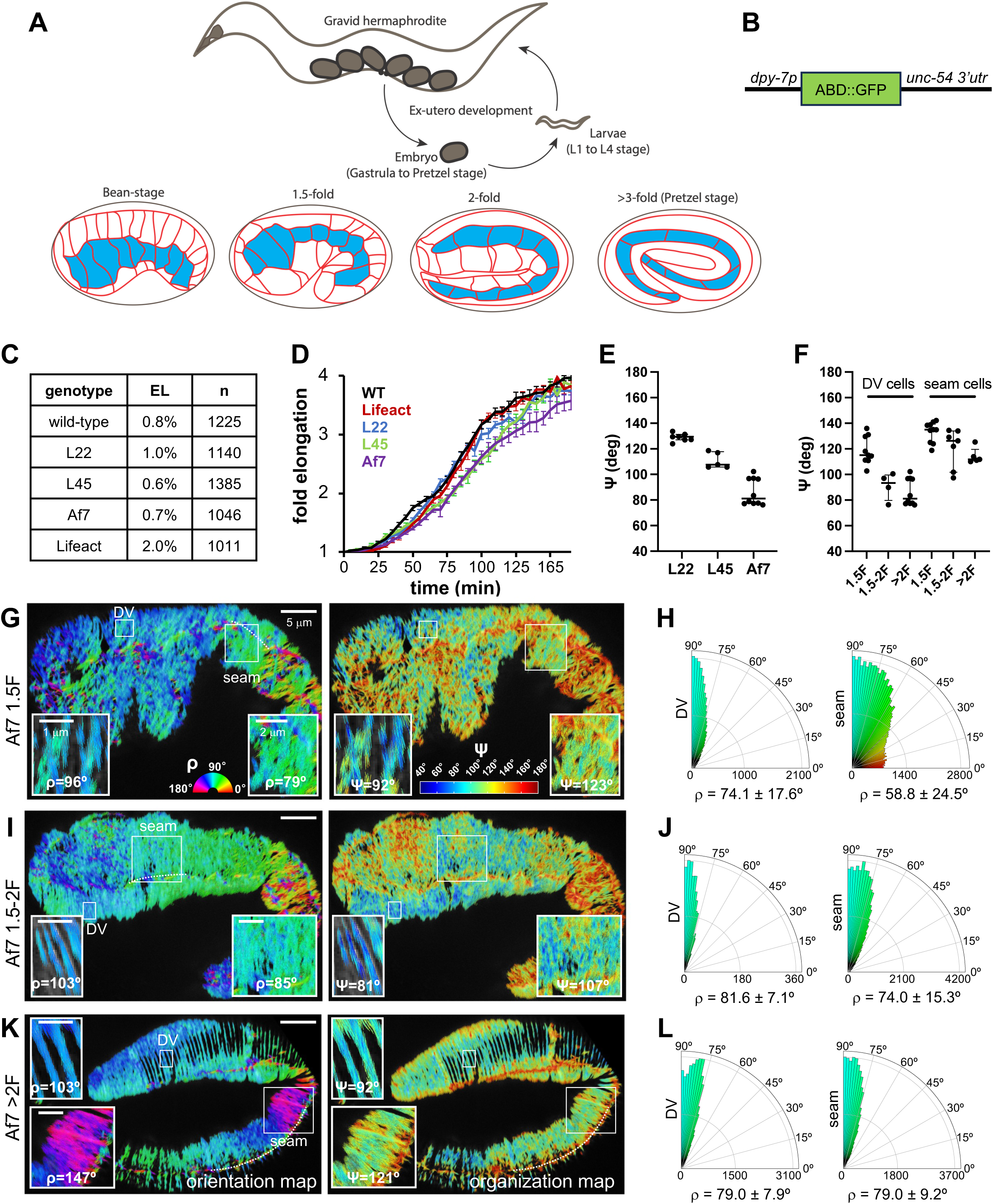
**Polarimetry measurements of actin filament organization in live elongating *C. elegans* embryos expressing selected reporters** (A) Schematic of a *C. elegans* gravid adult worm showing the ex-utero development of embryos (top) and an overview of embryonic elongation (bottom). The length of the embryo is used for staging: 2-fold (2F) stage means 2-fold increase in length from the beginning of elongation. Representative stages are shown; anterior is to the left and dorsal is up. (B) Schematic of the transgene design. The dpy-7 promoter drives expression of the actin organization reporters in epidermal cells. (C) Viability of *C. elegans* strains expressing different reporters assessed by the number of unhatched embryos 12-16 h after egg-laying. EL, embryonic lethality. n=number of scored embryos per genotype. See methods for details of the genotypes. (D) Embryonic growth curves, showing fold-change of embryonic length until hatching based on differential interference contrast (DIC) filming of the indicated *C. elegans* strains. Curves show means ± SEM. n=10 embryos were measured for each genotype. See also Figure S5G. (E) Polarimetry measurements of actin filament organization in the circumferential bundles of dorsal and ventral epidermal cells (DV cells) in >2-fold stage embryos expressing the indicated reporters. Scatter plots show the quantification of ψ angle distributions for each reporter. Scatter plots show medians with interquartile range. The number of embryos for each strain is, from left to right: 7, 5, 8. The respective median ψ values are 130, 108, 81°. See also Figure S5H,I. (F-L) Polarimetry measurements of actin filament reorganization in the epidermis during embryonic elongation. (F) Scatter plots show the quantification of ψ angle distributions in DV cells and in seam cells in 1.5-fold, 1.5-2-fold and >2-fold stage embryos expressing Af7 as shown in (G), (I), (K). Scatter plots show medians with interquartile range. The number of embryos for each stage is, from left to right: 9, 4, 8, 9, 7, 4. The respective median ψ values are 115, 93, 81, 135, 126, 112°. (G), (I), (K) Representative ρ (left) and ψ (right) stick maps in DV and seam cells in 1.5-fold (G), 1.5-2-fold (I) and >2-fold (K) stage embryos expressing Af7. Insets show zoom-ins of selected ROIs (white outlined boxes) in the respective cell types. Mean ρ and ψ values are shown for each ROI. For all panels, anterior is to the left and dorsal is up. (H), (J), (L) Polar histograms of ρ value distributions in DV cells (left) and seam cells (right) in 1.5-fold (H), 1.5-2-fold (J) and >2-fold (L) stage embryos expressing Af7. ρ values are represented with respect to the DV/seam boundary (dotted line in (G)): considering that Af7 dipoles are parallel to actin filaments, the more perpendicular mean actin filament orientations are to the boundary, the closer the angle values are to 90° and the narrower the respective distributions. Means ± SD are shown. The number of embryos for each stage and type of cells is as in panel F. See also Figure S5H-K.

Elongation of the *C. elegans* embryo proceeds along its anterior-posterior axis, increasing in length about fourfold and decreasing in circumference about threefold, with practically no cell divisions nor cell rearrangements ^43^. The epidermis comprises three major cell types, namely dorsal, ventral and lateral seam cells. Whereas actin filaments progressively form circumferential bundles in dorsal and ventral epidermal cells (DV cells), actin filaments in seam cells have been reported to stay rather disorganized throughout elongation ^44,45^. Recent work evidenced an interplay of stress anisotropy in seam cells and stiffness anisotropy in DV cells as critical for elongation through a circumferential squeezing-like mechanism ^45^, prompting us to quantify actin filament organization in DV and seam cells as a function of elongation.

The localization of L22, L45 and Af7 in the elongating epidermis was in line with phalloidin staining ^43^ and the localization of a GFP fusion to the actin-binding domain of the spectraplakin VAB-10 ^44,45^, showing the characteristic circumferential bundles in DV cells and a fuzzier mesh-like distribution in seam cells, the latter notably at earlier stages (Figure S5G-I). Considering that Af7 and L22 dipoles are parallel to actin filaments (Figure 2B,F) and that L45 dipoles are perpendicular to actin filaments (Figure 2C), quantification of actin filament organization in the circumferential bundles of DV cells in >2-fold elongated (>2F) embryos showed that actin filaments are parallel to the circumferential bundles (Figure 4K,L and Figure S5G-K), as expected, with ψ values comparable to the ones on SFs, suggesting highly aligned actin filaments (Figure 4E). Strikingly, quantification of actin filament orientation with respect to the DV/seam boundary showed that the polarization of actin filament orientation was already present in 1.5-fold elongated (1.5F) embryos before the formation of distinct circumferential bundles (Figure 4G,H and Figure S5G), with actin filament alignment increasing as bundles form (Figure 4F). What was, however, even more unexpected was that actin filament organization in seam cells was also polarized: seam cells in 1.5F embryos contained already actin filaments oriented perpendicular to the DV/seam boundary and with regions of actin filament alignment. As elongation proceeds, this polarization becomes progressively more pronounced, with actin filaments oriented essentially perpendicular to the DV/seam boundary, just like in DV cells, and with regions of high actin filament alignment (Figure 4F-L and Figure S5J,K).

### Actin filament organization reporters in *Drosophila*

We finally generated transgenic *Drosophila* expressing reporters with the inducible GAL4/ UAS expression system, allowing us to express them in the early embryo, the wing and the indirect flight muscle using respective tissue-specific promoters. Actin filament organization measurements in the actomyosin rings of living cellularizing embryos recapitulated earlier polarimetry measurements in fixed phalloidin-stained embryos ^21^ confirming the arrangement of highly aligned filaments following the contour of the associated membrane front (Figure 5A and Figure S6A). Apicolaterally to this front and at the basal-most part of the lateral membranes, a basal adherens junction has been reported to form ^46^. In this junction, actin filaments were found to be highly aligned to each other, following the junction contour, in line with a belt-like arrangement (Figure 5B).

**Figure 5.**
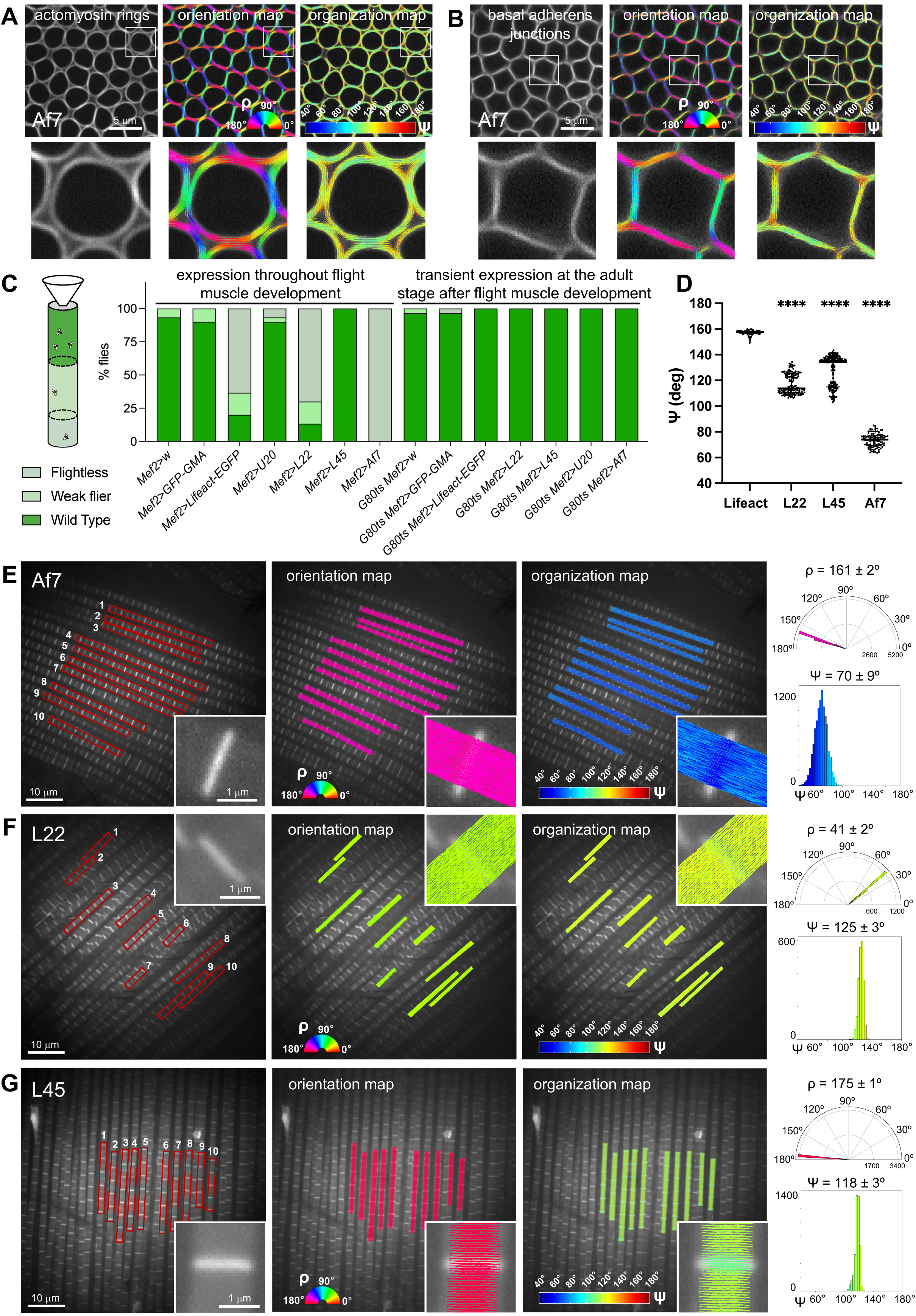
**Polarimetry measurements of actin filament organization in live cellularizing *Drosophila* embryos and live flight muscle expressing selected reporters** (A-B) Polarimetry measurements of actin filament organization in cellularizing *Drosophila* embryos expressing Af7, in the actomyosin rings associated with the invaginating membrane front (A) and at the basal adherens junctions right apicolaterally to the former (B). The left images are summed intensity images of the respective polarimetry stacks. The middle and right images show the measured ρ and ψ angles per pixel, respectively. Insets show zoom-ins of selected ROIs (white outlined boxes) (C) Flight tests were performed to compare the flight ability of strains expressing different actin-binding reporters. The left scheme depicts the flight test assay: depending on their flight ability, flies were scored as “wild-type”, “weak flier” or “flightless”. The bar graph on the right quantifies the respective percentages for each strain. Reporters were expressed under the control of a muscle-specific driver, Mef2-GAL4, either throughout muscle development (*Mef2>*), or transiently after muscle development (*G80ts Mef2>*). Thirty flies were scored in total for each strain in three independent experiments. The mean percentages of “wild-type”, “weak flier” or “flightless” flies are respectively, from left to right: 93,7,0; 90,10,0; 20,17,63; 90,3,7; 13,17,70; 100,0,0; 0,0,100; 97,3,0; 97,0,3; 100,0,0; 100,0,0; 100,0,0; 100,0,0; 100,0,0. (D-G) Polarimetry measurements of actin filament organization in live flight muscle expressing the indicated reporters. All reporters were expressed throughout flight muscle development except Af7 that was expressed transiently at the adult stage after flight muscle development. Scatter plots in (D) show the quantification of ψ angle distributions for each reporter measured as shown in (E)-(G). Scatter plots show medians with interquartile range. One datapoint represents one myofibril (see methods for details). The number of myofibrils for each strain is, from left to right: 70, 140, 190, 100. The respective median ψ values are 157, 114, 134, 74°. Statistical significance was obtained using a non-parametric Mann-Whitney test. The different constructs were compared to Lifeact; **** P<0.0001. (E)-(G) Left panels are summed intensity images of the respective polarimetry stacks. Ten myofibrils (red outlined boxes in left panels) were quantified in each field of view. Insets show zoom-ins of a selected sarcomere labeled by the respective reporter. Middle and right panels are ρ and ψ stick maps in the flight muscle of the shown hemithorax, respectively. The rightmost panels for each strain show examples of polar histograms of ρ value distributions and histograms of ψ value distributions for a single myofibril. Means ± SD are shown. See also Figure S6A-D.

We assessed the functionality of the reporters in two systems that depend on the integrity of the actin cytoskeleton, namely the adult wing and the adult flight muscle. To measure their effect on wing growth, we compared the wing area in wings expressing the reporters with the one in wings bearing only the wing-specific promoter (control), wings expressing the original Lifeact-GFP, and wings expressing a dominant-negative form of the insulin receptor expected to reduce wing growth. Wing growth was indeed reduced by ∼46% for the latter, but was largely comparable, within less than 8% of change, among the control and the other strains (Figure S6B). As a second functionality test, we performed flight tests to compare the flight ability of strains expressing different reporters with the one of strains bearing only the muscle-specific promoter (control) and strains expressing two widely used actin localization reporters: the actin-binding domain of moesin (GFP-GMA) and Lifeact-GFP (Figure 5C). Although all reporters localized largely as expected in the muscle sarcomeres (Figure S6C), flight tests showed significant differences among the strains. The flight ability of flies expressing L45 and U20 was comparable to the one of control and GFP-GMA flies. However, flies expressing either L22 or the original Lifeact-GFP, performed very poorly, with Af7 expressing flies being entirely flightless (Figure 5C); the development of the flight muscle in the latter strain was altogether impaired (Figure S6C). Interestingly, in line with the fission yeast data, we noted again that the N-terminally tagged L45 appeared much less perturbative that L22 and the original Lifeact, both of which are C-terminally tagged. The muscle-specific promoter being active in all embryonic, larval and adult muscle, we attributed this dramatic effect to the expression of the reporters throughout muscle development, and reasoned that their limited expression after muscle development could be less perturbative. In support of this scenario, and with the use of the temperature-sensitive GAL80ts system to express the reporters only in a narrow time window of a few days in adult muscle, the flight ability of all strains was improved, including the strains expressing the original Lifeact-GFP, L22, and Af7 (Figure 5C). Muscle morphology and sarcomere localization were comparable to the control in all cases (Figure S6D). Importantly, despite their limited temporal expression, the fluorescence levels of the reporters rendered them usable for polarimetry measurements in the flight muscle.

To this end, we dissected live flight muscle expressing L22, L45 and Af7, and the original Lifeact-GFP for comparison, and measured actin filament organization in the respective myofibrils. ψ values in the original Lifeact-GFP were too high to render this fusion usable for organization measurements, but L22, L45 and Af7 behaved as expected, with Af7 further displaying the lowest ψ values i.e., the highest filament alignment, from all systems (Figure 5D), in line with the crystal-like arrangement of actin filaments in sarcomeres reported by EM ^22^. Considering the GFP dipole orientations of L22, L45 and Af7 with respect to actin filaments, actin filament orientation and alignment maps revealed the expected high order within and across different myofibrils in the muscle (Figure 5E-G).

### Binding affinities of organization reporters to actin filaments

The affinity of any actin-binding probe to actin filaments is an important property that influences its labeling efficiency, the fluorescent background from nonfilamentous actin, as well as the extent of perturbation of actin dynamics. To measure the binding affinities for the four best performing reporters that we characterized *in vivo*, namely Af7, L22, L45 and U20 (Figure S6E-I), we employed fluorescence microscopy and co-sedimentation assays using recombinant purified reporters and reconstituted actin filaments *in vitro* (see methods for details). The binding affinities were in line with the values reported in the literature ^38,41,47,48^ suggesting that constraining GFP mobility in these constructs did not impact actin filament binding affinity.

### Engineering red FP-based actin filament organization reporters

Having succeeded in generating GFP-based reporters for the different ABDs, we naturally embarked on the making of the red FP fusion counterparts. We generated selectively sfCherry2-based terminal fusions for Lifeact and Affimer6. The best performing constructs were Af30 (Affimer6-sfCherry2ΔN12) and L81 (sfCherry2ΔC4-Lifeact) combining full-length Affimer6 and Lifeact with the most extensively truncated sfCherry2 termini (Figure S7A-F), very similarly to the respective best performing GFP-based ones, and are ideally placed for F-actin organization measurements in live cells and tissues (Table 1; see Data S1 for a discussion of the choice of the red FP).

### Engineering actin filament organization reporters using G-actin

Finally, with the ambition to generate constrained GFP fusions to G-actin directly, we generated terminal and intramolecular fusions of human non-muscle beta G-actin (Figure 6A-C) without linker sequences and with shortened GFP termini (Figure S7G-I). Besides the use of msfGFP, we employed two additional tags for G-actin fusions, the GFP strand β11 and a tetracysteine peptide (Figure 6A and Data S1). Data S1 provides a comprehensive account of the reasoning behind the choice of the G-actin terminus and insertion site for terminal and intramolecular fusions, respectively.

**Figure 6.**
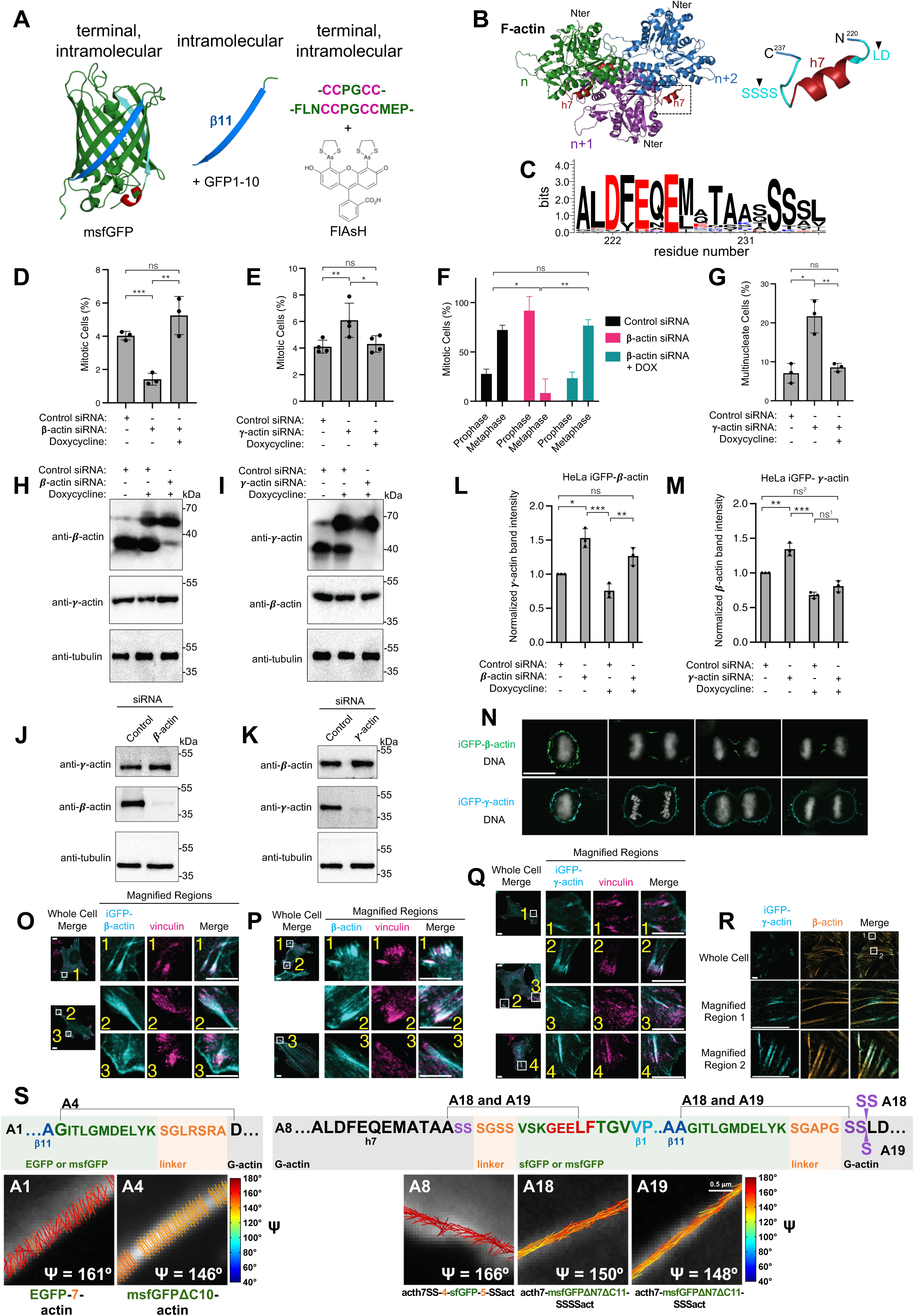
**Engineering of G-actin-based actin filament organization reporters for live-cell polarimetry** (A) Designs used in this study to immobilize genetically-encoded fluorophore fusions to G-actin. For G-actin terminal fusions, msfGFP or tetracysteine peptides were fused to the N-terminus of G-actin (left and right). For G-actin intramolecular fusions, msfGFP, the β11 strand alone, or tetracysteine peptides were placed intramolecularly within the G-actin structure (left, middle and right). (B-C) Ribbon representation of F-actin with three consecutive G-actin monomers colored in green, magenta and blue (PDB 5JLF) (B, left). Helix h7, used as an insertion site in intramolecular fusions, is shown in red. A close-up view of h7 (dashed box) shows residues in the loops (in cyan) flanking the helix, with arrowheads pointing to the insertion sites used in intramolecular fusions (B, right). (C) WebLogo3 representation of the conservation of residues in h7 and the flanking residues. Forty-five actin sequences were used for this representation, including organisms as diverse as *Drosophila*, fungi, Dictyostelium, Arabidopsis, and sea animals. Negatively- and positively-charged residues are shown in red and blue, respectively. (D-R) Functional characterization of intramolecular GFP (iGFP) fusions showing their usability for labeling specific G-actin isoforms. (D) Quantification of mitotic cells from an asynchronous population of HeLa FRT iGFP-beta-actin expressing cells treated as indicated. N = 3 for all conditions; n = 543 for ‘Control siRNA’; n = 715 for ‘Beta Actin siRNA – dox’; n = 321 for ‘Beta Actin siRNA + dox’. **p = 5.2×10^−3^, *** p = 5.0×10^−4^, ‘ns’ = 0.15 by two tailed t-test. (E) Quantification of mitotic cells from an asynchronous population of HeLa FRT iGFP-gamma-actin expressing cells treated as indicated. N = 4 for all conditions; n = 439 for ‘Control siRNA’; n = 451 for ‘Gamma Actin siRNA – dox’; n = 418 for ‘Gamma Actin siRNA + dox’. *p = 0.047; **p = 0.028, ‘ns’ = 0.62 by two tailed t-test. (F) Classification of mitotic cells described in panel E as either prophasic or metaphasic. As in E, N = 3 for all conditions. A total of 22 metaphase cells were scored for ‘Control siRNA’; 11 metaphase cells for ‘Beta siRNA -dox”; 17 metaphase cells for ‘Beta siRNA +dox”. *p = 0.010, **p = 0.0069, ‘ns’ = 0.15 by multiple unpaired t-tests with Welch correction. (G) Quantification of multinucleated cells from an asynchronous population of HeLa FRT iGFP-gamma-actin expressing cells treated as indicated. N = 3 for all conditions; n = 690 cells scored for ‘Control siRNA’; n = 640 cells for ‘Gamma siRNA -dox”; n = 621 cells for ‘Gamma siRNA +dox”. *p = 0.0069, **p = 0.0066, ‘ns’ = 0.40 by two-tailed t-test. (H) Western blot of cell lysates prepared from stable HeLa FRT iGFP-beta-actin cells treated as indicated. Lysates were probed with antibodies recognizing gamma-actin, beta-actin, and tubulin as a loading control. Presented blots are representative of three independent experiments. (I) Western blot of cell lysates prepared from stable HeLa FRT iGFP-gamma-actin cells treated as indicated. Lysates were probed with antibodies recognizing beta-actin, gamma-actin, and tubulin as a loading control. Presented blots are representative of three independent experiments. (J) Western blot of cell lysates prepared from stable HeLa FRT iGFP-beta actin cells treated as indicated. Lysates were probed with antibodies recognizing gamma-actin, beta-actin, and tubulin as a loading control. Presented blots are representative of three independent experiments. (K) Western blot of cell lysates prepared from stable HeLa FRT iGFP-gamma-actin cells treated as indicated. Lysates were probed with antibodies recognizing beta-actin, gamma-actin, and tubulin as a loading control. Presented blots are representative of three independent experiments. (L) Quantification of normalized gamma-actin band intensities from panels H and J. Band intensities were normalized to respective uninduced control siRNA-treated conditions. N = 3 independent experiments. *p = 0.022, ‘ns’ = 0.075 by one sample t test; **p = 6.3×10^−3^, ***p = 1.5×10^−3^ by two-tailed t-test. (M) Quantification of normalized beta-actin band intensities from panels I and K. Band intensities were normalized to respective uninduced control siRNA-treated conditions. N = 3 independent experiments. ‘ns^2^’ = 0.054, **p = 0.0064 by one sample t test; ‘ns^1^’ = 0.072, ***p = 3.8×10^− 4^ by two-tailed t-test. (N) Micrographs of mitotic and cytokinetic iGFP-beta-actin and iGFP-gamma-actin expressing HeLa FRT cells depleted of the corresponding endogenous actin isoform. Scale bar represents 10 μm. (O) Micrographs of iGFP-beta-actin expressing cells co-stained with vinculin, showing colocalization of iGFP-beta-actin with focal adhesions. iGFP-beta-actin is also visualized in focal adhesion-associated stress fibers and on membrane ruffles that are vinculin-negative. Scale bars represent 5 μm for both whole cell and magnified images. Cells were depleted of endogenous beta actin. (P) Micrographs of HeLa cells co-stained for beta-actin and vinculin, showing colocalization of beta-actin with focal adhesions. Beta-actin is also visualized in focal adhesion-associated stress fibers and on membrane ruffles that are vinculin-negative. Scale bars represent 5 μm for both whole cell and magnified images. Cells were depleted of endogenous beta actin. (Q) Micrographs of iGFP-gamma-actin expressing cells co-stained with vinculin, showing colocalization of iGFP-gamma-actin with a subset of focal adhesions. iGFP-gamma-actin is also visualized on membrane ruffles that are vinculin-negative. Scale bars represent 5 μm for both whole cell and magnified images. Cells were depleted of endogenous gamma actin. (R) Micrographs of iGFP-gamma-actin expressing cells co-stained with antibody recognizing beta-actin, showing their distinct localization patterns on stress fibers. Scale bars represent 5 μm for both whole cell and magnified images. Cells were depleted of endogenous gamma actin. (S) Engineering of G-actin-based actin filament organization reporters. Representative designs for constraining GFP mobility in N-terminal (left) and intramolecular (right) GFP fusions are illustrated for selected fusions, including for the best performing reporters (A4, A18, see Table 1). The full screen is shown in Figure S7G-I. The top and bottom panels show the primary sequences and respective ψ stick maps as described for Figure 2B.

To test if intramolecular human G-actin GFP fusions (“iGFP-actin” for short) are functional, we generated intramolecular full-length msfGFP fusions to the two vertebrate non-muscle actin isoforms, namely human beta- and gamma-actin isoforms. We used the same insertion site as for construct A8 (Figures 6S and S7G). The two isoforms have distinct localizations and functions ^49–52^, and intramolecular GFP fusions would provide new less-perturbative tools to study actin isoform function given that they differ only by four amino acids at their very N-terminus. To assess if the iGFP-actin fusions can functionally substitute for their respective endogenous actin isoform, we generated stable inducible HeLa cell lines expressing the iGFP-actin isoform fusions. First, we expressed the iGFP-actin fusions upon depletion of endogenous actins and analyzed dividing cells, a process where beta and gamma actin have well-defined functions and localizations ^49,53^. We observed that depletion of beta actin decreased the proportion of cells undergoing mitosis while depletion of gamma actin increased this proportion. Both phenotypes were rescued to control levels by expression of the respective iGFP-actin (Figure 6D-F). Furthermore, depletion of gamma actin increased the proportion of multinucleated cells arising from failed cytokinesis, which was also rescued by expression of iGFP-gamma actin (Figure 6G). Expression levels of iGFP-actins were similar to those of their endogenous counterparts; likewise, depletion of endogenous actins was observed to be efficient (Figure 6H, I). To investigate if the iGFP-actins demonstrated similar localization profiles to their endogenous counterparts, we analyzed fixed iGFP-actin expressing HeLa cells depleted of the respective endogenous actin. In dividing cells, iGFP-beta actin localized to the cell cortex in metaphase and to the cytokinetic furrow in anaphase and telophase, while iGFP-gamma actin localized to the cell cortex and later the polar cortex (Figure 6N); both in keeping with the localizations of their respective endogenous actin isoform ^53^. Beta and gamma actin filament production at these sites is mediated by the formins DIAPH3 and 1 respectively ^52,53^, who discriminate actin isoforms predominantly on the basis of the divergent actin N-terminal amino acid sequences and the unique FH2 linker region in each formin ^54^. These observations suggest that insertion of GFP into the actin sequence does not hinder the selectivity of formins for specific actin isoforms. In interphase cells, iGFP-beta actin localized to focal adhesions and stress fibers, similar to endogenous beta actin (Figure 6O,P), while iGFP-gamma actin was observed at focal adhesions (Figure 6Q). iGFP-gamma actin expressing cells co-stained with endogenous beta actin corroborated these differences, with iGFP-gamma actin found at the ends of stress fibers and beta actin throughout stress fibers (Figure 6R).

It has been reported that altering the levels of one cytoplasmic actin isoform leads to compensatory changes in the expression level of the other ^55^. In the absence of iGFP-actin expression, depletion of endogenous beta or gamma actin led to compensatory increases in the expression levels of the opposite endogenous isoform (Figure 6J-M). Expression of iGFP-beta or iGFP-gamma actin significantly reduced levels of the opposite endogenous isoform, and was sufficient to restore the opposite isoform to control expression levels when cells expressing iGFP-actins were also depleted of the same endogenous isoform (Figure 6J-M). We conclude that iGFP-actins are indeed capable of functionally substituting for their endogenous counterparts.

The screened G-actin fusions are shown in Figure S7G-I. All F-actin-binding fusions localized overall similarly to the ABD-GFP fusions (Figure S7I; see Data S1 for a discussion on observed differences). Terminal and intramolecular GFP fusions without linkers and with extensively shortened GFP termini were most efficient to constrain GFP mobility. Fusions A4 (msfGFPΔC10-actin) and A18 (actin-h7-msfGFPΔN7ΔC11-SSSSactin) are the best performing fusions for F-actin organization measurements in live cells (Figure 6S and Table 1).

### Selection of actin filament organization reporters

To guide the selection of an actin filament organization reporter for a given application, Table 1 summarizes key features of the best performing reporters. The two most important selection criteria are localization and functionality: the selected reporter should localize to the actin filament population of interest while minimally perturbing the relevant biological process. The latter depends on the cellular context under study ^56^. In line with the functionality tests in this work (”functional readouts” and “perturbative character” columns of Table 1), several studies have documented differential effects of actin localization probes depending on the exact process in question ^41,47,57–61^: a given reporter might have dramatic perturbative effects in one process but have no effect in a different context. Thus, functional readouts are essential to consider. To minimize perturbation, we recommend the use of inducible expression systems or weak promoters to obtain the lowest expression levels that generate reasonable signal-to-noise ratios for fluorescence detection. The extent to which a given reporter will label an actin filament population of interest is difficult to predict. A few reports have correlated the labeling of distinct actin pools with specific actin probes ^61^, but the precise link between actin filament assembly geometry and dynamics and actin probe labeling efficiency has not been established. Labeling efficiency will depend on the combined effect of binding affinity (Table 1), actin filament assembly geometry and turnover, and possible competition with other actin-binding proteins, and has to be determined empirically.

When a minimally perturbative organization reporter has been identified, the extent of GFP immobilization, i.e. the ψ values obtained on reference structures of highly aligned actin filaments (Table 1), is the next selection criterion. The lower the ψ values, the larger the range of ψ angles will be that can be explored. Af7 is the reporter with the lowest ψ values, reflecting a highly constrained GFP, thus making it the most sensitive one for detecting small changes in actin filament organization. A4 and A18 are, on the other hand, the ones with the highest ψ values, most likely due to the inherent flexibility of the actin N-terminus and of the insertion site loop in terminal and intramolecular fusions, respectively. Despite their comparatively high ψ values, these reporters are still useful for studies focusing on G-actin directly, notably actin isoforms.

Finally, dipole orientation with respect to the actin filament axis (Table 1) is a last selection criterium. Standard polarimetry techniques are typically limited to in-plane (2D) orientation measurements, with off-plane orientations leading to an overestimation of ψ values ^34^. Due to the helical nature of actin filaments, the precise actin filament assembly geometry will determine the off-plane contribution, and GFP dipoles of a given reporter can be found either parallel or perpendicular to the imaging plane. For example, parallel dipoles (Af7, L22) will stay parallel when associated with actin filaments mostly in-plane (<45°off the xy imaging plane), but will become perpendicular for actin filaments perpendicular to the xy imaging plane. Perpendicular dipoles (L45, U20) should be used in the latter case since those would now be parallel to the imaging plane.

## DISCUSSION

We succeeded in constraining GFP mobility in fusions to widely used actin localization reporters, thus providing the possibility to correlate any biological process of interest with live measurements of actin filament organization by polarimetry. We further provide reference angle values in different model structures (Table 1) for comparison and interpretation of measurements in any F-actin population of interest. Even though ensemble measurements cannot resolve individual filaments, their molecular-scale organization at a given image pixel is detectable and can be quantified by polarimetry. Moreover, the mesh size of the actin network is on the order of 100 nm or less ^62^, so far only attainable in a fixed cell context with EM and single molecule localization microscopy. Thus, the capacity to obtain quantitative measurements of actin filament organization per image pixel in a living cellular context is of significant added value.

Cells continuously and dynamically remodel actin filaments to accomplish specific biological functions. Bottom-up approaches with purified proteins are key for assigning specific functions, for example filament branching, to distinct actin-binding proteins ^63^. Whether and how these assigned functions account for F-actin organization in cells can now be formally tested by combining live cell organization measurements with mutants or treatments affecting one or several interactors.

The use of the reporters in the context of cell and tissue morphogenesis promises to uncover how specific geometries of actin filaments contribute to function and provide new insights into the developmental regulation of F-actin organization. The genetically-encoded character of the reporters provides them with the possibility to be used in the context of animal disease models, enabling potentially *in vivo* studies on the role of any gene of interest in F-actin organization in the context of pathophysiology. Being able to measure changes in filament organization in real-time will additionally help generate accurate biophysical models with experimentally testable predictions regarding how F-actin organization impacts biomechanics.

The F-actin organization reporters are compatible with all fluorescence microscopy techniques routinely used for live ensemble imaging that can be coupled with polarized fluorescence imaging, such as transmission polarized microscopy ^64,65^, wide-field ^66^, confocal ^34^ and spinning disk confocal microscopy ^67^, total internal reflection microscopy ^68^ and two-photon microscopy ^69–72^. The reporters are also compatible with super-resolution imaging techniques, either based on polarized structured illumination microscopy employing standard FPs ^73,74^, or based on single molecule orientation and localization microscopy using photoactivatable or photoconvertible FPs ^75^. We note that the introduction of a single point mutation to sfGFP or of a few point mutations to sfCherry2 are sufficient to generate the respective photoactivatable versions ^76,77^: given that these mutations are within the barrel structure, we do not expect them to alter the mobility of the FP. Our strategy can thus be simply adapted to the context of single-molecule organization measurements using, for example, single-particle tracking PALM coupled to polarization splitting ^31,32^ or Point Spread Function engineering ^75^. All measurements in this study have used a spinning disk confocal microscope with polarized excitation for second-scale measurements. If higher temporal resolution is needed, polarization splitting would allow for subsecond-scale measurements using the same reporters ^24,33,66,78–80^. Our open-source software, PyPOLAR, has been designed explicitly for use by biologists and biophysicists to further facilitate the use of the reporters.

This study reports constrained GFP fusions to five widely used F-actin localization reporters, including G-actin itself, increasing the chances that one is able to label and measure the organization of any F-actin pools of interest. Constrained Cherry fusions provide additional experimental flexibility, notably for two-color imaging with GFP fusions to any protein of interest while measuring F-actin organization. Our Cherry fusions are also compatible with the standard CFP/YFP-like donor/acceptor pairs used for FRET-based force measurements ^81^, enabling a direct correlation between F-actin organization and mechanical properties.

Finally, even though our results relate primarily to sfGFP and sfCherry2 fusions to ABDs, our designs can be rationally applied to other FPs and fusions to proteins other than actin-binding ones, to generate new tools for measuring any protein organization by polarimetry. Importantly, the highly constrained sfGFP and sfCherry2 fusions to Affimers (Af7 and Af30) have been designed in a manner that does not affect the Affimer protein scaffold nor the variable binding loops and thus provide a straightforward, universal method for constrained FP fusions to Affimers against any protein of interest ^38,82^. Structural and cell biology approaches employing FP-based sensors to probe protein proximity, protein-protein interactions and mechanical forces, notably the ones using FRET, are also likely to benefit from using constrained FP fusions.

## Limitations of the study

Our study did not explore to what extent the measured ψ values depend on the packing of actin filaments, namely interfilament spacing. It is conceivable that the mobility of GFP is constrained in the case of tightly packed actin filaments, for example, in the case of small and rigid actin cross-linkers ^83^. Such a scenario can be tested using cell-free reconstituted actin geometries with controlled packing properties. The second limitation is that polarized fluorescence cannot distinguish molecules pointing in one direction from molecules pointing in the opposite direction, i.e. it cannot make out, for example, 45° from 180°+45°=225°. As a consequence, it is not possible to detect processes sensitive to the pointing direction, such as actin filament polarity. The measured orientation of an actin filament bundle will be the same whether the contained filaments are parallel or antiparallel.

## RESOURCE AVAILABILITY

### Lead contact

Further information and requests for resources, reagents, and software should be directed to and will be fulfilled by the lead contact, Manos Mavrakis.

### Materials availability

All plasmids and strains generated in this study are available upon request. We have deposited the mammalian expression plasmids coding for the best performing reporters (Table 1) and for human iGFP-beta- and -gamma-actin with the nonprofit repository Addgene (see Table S1 for their Addgene#).

### Data and code availability

The datasets supporting the current study have not been deposited in a public repository but are available from the lead contact upon request. The codes and softwares developed and used in this study are open source and available on GitHub under a BSD license; the software identifiers are listed in the Key Resources Table and the links are provided in the respective method details sections.

## ACKNOWLEDGMENTS

We thank Louwrens van Dellen (I. Fresnel, Marseille, France) for the development of the image acquisition software. This research has received funding from the French National Research Agency (ANR) grants Equipex+ IDEC (France 2030 investment plan ANR-21-ESRE-0002), 3DPolariSR (ANR-20-CE42-0003), SEPTIMORF (ANR-17-CE13-0014), SEPTISS (ANR-22-CE13-0039), and from the France-BioImaging infrastructure (ANR-10-INBS-04). This research has further received funding from the Excellence Initiative of Aix-Marseille University – A*Midex (A*Midex grant NEUROPOL), a French “Investissements d’Avenir” program, the Fondation pour la Recherche Médicale (FRM grant ING20150531962) and SATT Sud-Est. A.J. and R.P. were supported by the Bettencourt-Schueller Fondation (Impulscience grant n° 1235).

## AUTHOR CONTRIBUTIONS

C.S.M. : Methodology, Investigation, Validation, Formal analysis, Visualization, Writing – original draft, Writing – review & editing; F.I. : Investigation; S.K.S. : Investigation, Validation, Formal analysis, Visualization, Writing – original draft, Writing – review & editing; T.C.P. : Investigation, Validation, Visualization, Formal analysis; C.S. : Investigation, Validation, Formal analysis, Visualization, Writing – original draft, Writing – review & editing; M.R.-L. : Investigation; C.N.P. : Investigation; S.O. : Investigation; R.P. : Investigation; M.G. : Investigation; A.L. : Investigation; S.R.B. : Investigation; L.R. : Investigation; F.A. : Investigation; C.V.R. : Investigation; F.S. : Writing – review & editing, Supervision, Resources; F.R. : Writing – review & editing, Supervision, Resources; A.W. : Writing – review & editing, Supervision, Resources; L.L. : Investigation, Validation, Formal analysis, Visualization, Writing – original draft, Writing – review & editing; J.-D.P. : Investigation, Visualization, Writing – original draft, Writing – review & editing; A.J. : Investigation, Validation, Formal analysis, Visualization, Writing – original draft, Writing – review & editing; S.C. : Investigation, Validation, Formal analysis, Visualization, Writing – original draft, Writing – review & editing, Resources; S.A.R. : Investigation, Validation, Formal analysis, Visualization, Writing – original draft, Writing – review & editing, Supervision, Resources ; C.C. : Software; S.B. : Conceptualization, Methodology, Investigation, Software, Writing – original draft, Writing – review & editing, Supervision, Resources, Project administration, Funding acquisition; M.M. : Conceptualization, Methodology, Investigation, Software, Validation, Formal analysis, Visualization, Writing – original draft, Writing – review & editing, Supervision, Resources, Project administration, Funding acquisition

---

**Figure S1.**
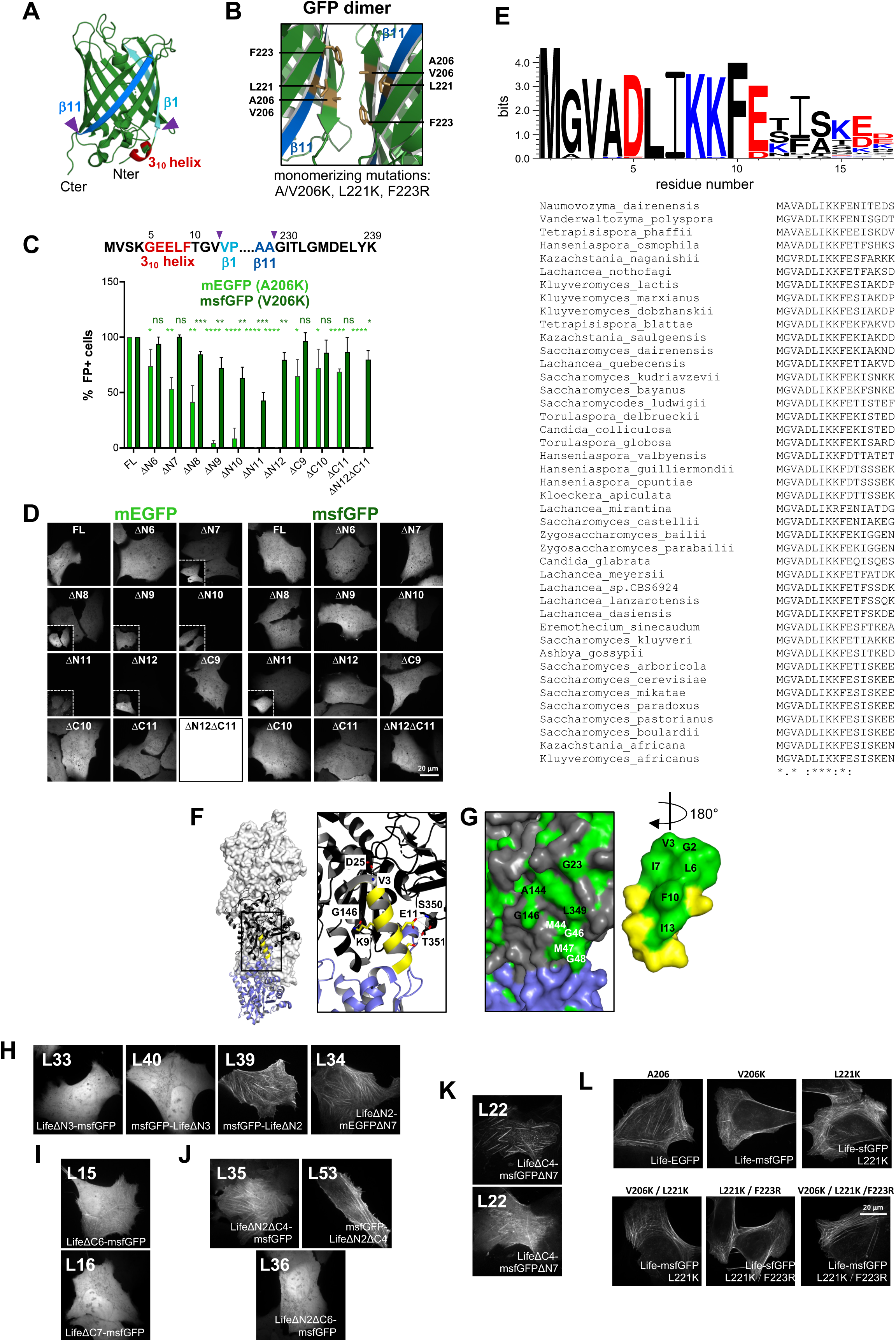
Engineering of GFP and of Lifeact-based actin filament organization reporters for live-cell polarimetry. Related to. **Figures 2 and S2.** (A) Ribbon representation of the crystal structure of superfolder GFP (sfGFP) (PDB 2B3P). The N-terminal 3_10_ helix and beta-strands β1 and β11 are depicted in red, cyan and blue, respectively. Magenta arrowheads point to the beginning of β1 and the end of β11. (B) Ribbon representation of the GFP dimerization interface from the crystal structure of GFP (PDB 1GFL). The hydrophobic residues A206 (V206 for sfGFP), L221 and F223 in the dimer interface are shown in brown. The respective monomerizing mutations used for impairing dimerization are shown. (C) Top, amino acid sequence of the N- and C-termini of monomeric EGFP (mEGFP) and monomeric sfGFP (msfGFP). The depicted secondary structure elements, color code and arrowheads are as in (F). Residue numbering is as shown. Bottom, screening of N- and C-terminal truncation mutants of mEGFP and msfGFP using fluorescence-activated cell sorting (FACS). Bars (mean + SD) depict the measured percentages of fluorescence-positive (FP+) cells for full-length (FL), N-terminally (ΔN) and C-terminally (ΔC) truncated proteins. The mean values are, from left to right: 100, 100, 74, 94, 53, 100, 42, 85, 4, 72, 9, 63, 0.4, 43, 0.1, 80, 65, 96, 72, 86, 69, 87, 0.2, and 80. Data are from three independent experiments. Statistical significance was obtained using an unpaired t-test. The different constructs were compared to the respective FL; ns=not significant (P>0.05); * P<0.05, ** P<0.01, *** P<0.001, **** P<0.0001. (D) Screening of the GFP constructs used in (C) with spinning disk fluorescence microscopy. Representative images of live cells expressing each construct are shown. For the sake of comparison, images are displayed with the same intensity range. In the case of weakly fluorescent cells, insets show contrast-enhanced images. No fluorescence was detectable for mEGFPΔN12ΔC11. (E) Top, WebLogo3 representation of the conservation of residues in the Lifeact sequence. Negatively- and positively-charged residues are shown in red and blue, respectively. Bottom, the Lifeact sequences from 43 different budding yeast strains used for the WebLogo representation are shown. The consensus symbols are from the ClustalO multiple sequence alignment: **\***, fully conserved residue; :, conservation between residues with strongly similar physicochemical properties; ., conservation between residues with weakly similar physicochemical properties. (F-G) 3D structure of the Lifeact-F-actin complex ^47,84^. (F) Surface representation (light grey) of three G-actin monomers within an actin filament (PDB 7AD9). Two neighboring actin subunits (n, n+2), colored in black and blue, are shown in ribbon representation, and Lifeact is shown in yellow. A close-up view (black outlined box on the left) illustrates polar interactions at the actin-Lifeact interface, with key residues and their side chains depicted in stick representation. (G) Surface representation of the actin-Lifeact interface shown in the close-up view of (F), highlighting hydrophobic residues in green. Residues from actin subunits n and n+2 are colored in black and white, respectively. Lifeact is rotated by 180° to visualize the hydrophobic residues facing the actin monomer. (H-J) Representative images of U2OS cells expressing the indicated reporters for assessing the contribution of specific Lifeact residues to actin binding. The localization of the reporters (diffuse cytosolic vs binding to SFs) shows that V3 is essential (H), that the six C-terminal residues are not essential (I), and that G2 is not essential but its absence can compromise actin binding when combined with other truncations (J). (K) Lifeact-based reporters localize both to SFs and to mitochondria: two different z-planes in the same cell show L22 on SFs (top plane) and mitochondria (bottom plane). (L) Fluorescence images of U2OS cells expressing Lifeact fusions to EGFP or to sfGFP bearing one, two or three monomerizing mutations. Lifeact localizes to arc nodes in all cases.

**Figure S2.**
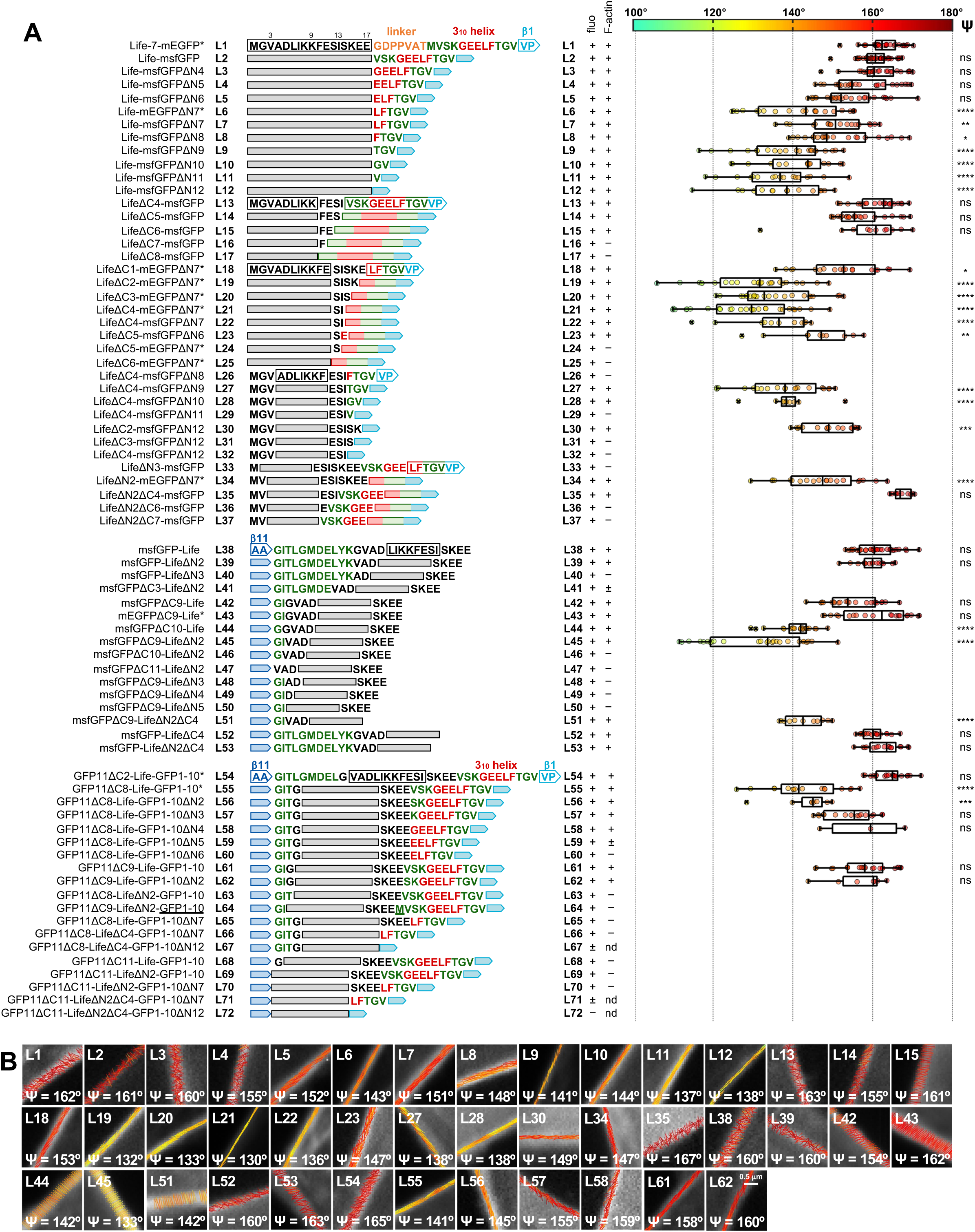
Engineering of Lifeact-based actin filament organization reporters for live-cell polarimetry. Related to. **Figure 2** (A) List of Lifeact (”Life” for short)-GFP fusions tested for their usability in live polarimetry measurements in U2OS cells expressing each fusion. All constructs were expressed with a CMV_trunc_ promoter except the ones with an asterisk (*) which used a CMV promoter. The left column lists the tested fusions as follows: a short name for each fusion (e.g., Life-msfGFPΔN6) followed by its designated number (”L5” in this case) and its primary sequence. Secondary structure elements of GFP are color-coded as in Fig.S1A. To facilitate the tracking of modifications, only the region of the primary sequence that is modified is shown. Sequences that stay unchanged from one construct to the next one are shown with an outlined box for the first one (e.g. L1) and then represented as gray-filled boxes in subsequent constructs (in this case, L2-L12). Additional columns report whether the construct is fluorescent (column “fluo”) and if it binds actin SFs (column “F-actin”). Constructs were classified as fluorescent (+), very weakly fluorescent (±), or nonfluorescent (-), and as binding to F-actin (+), very weakly binding to F-actin (±), or not binding to F-actin (-). For very weakly fluorescent or nonfluorescent constructs, F-actin binding was not determined (nd). Box plots on the far right depict the distribution of ψ angle measurements on SFs for the respective constructs. Box plots are depicted as in Fig.1C. Polarimetry measurements were not performed for constructs that were weakly fluorescent or weak F-actin binders and thus not usable for routine polarimetry. The number of measurements (see methods for details), from top to bottom, are n= 30, 40, 25, 25, 26, 21, 18, 22, 21, 15, 17, 15, 21, 22, 11, 19, 24, 23, 29, 11, 10, 20, 9, 17, 20, 10, 33, 15, 23, 20, 21, 26, 11, 18, 16, 21, 16, 8, 16, 3, 22, 7. The respective median ψ values are 162, 161, 160, 155, 152, 143, 151, 148, 141, 144, 137, 138, 163, 155, 161, 153, 132, 133, 130, 136, 147, 138, 138, 149, 147, 167, 160, 160, 154, 162, 142, 134, 142, 160, 163, 165, 141, 145, 155, 159, 158, 160°. Statistical significance (right-most column) was obtained using a non-parametric Kruskal-Wallis test followed by a Dunn’s multiple comparisons test. The different constructs were compared to L1; ns=not significant (P>0.05); * P<0.05, ** P<0.01, *** P<0.001, **** P<0.0001. (B) Representative ψ stick maps on SFs from measurements in live cells expressing the indicated fusions, with mean ψ values indicated. The selected images correspond to median ψ values of the respective distributions.

**Figure S3.**
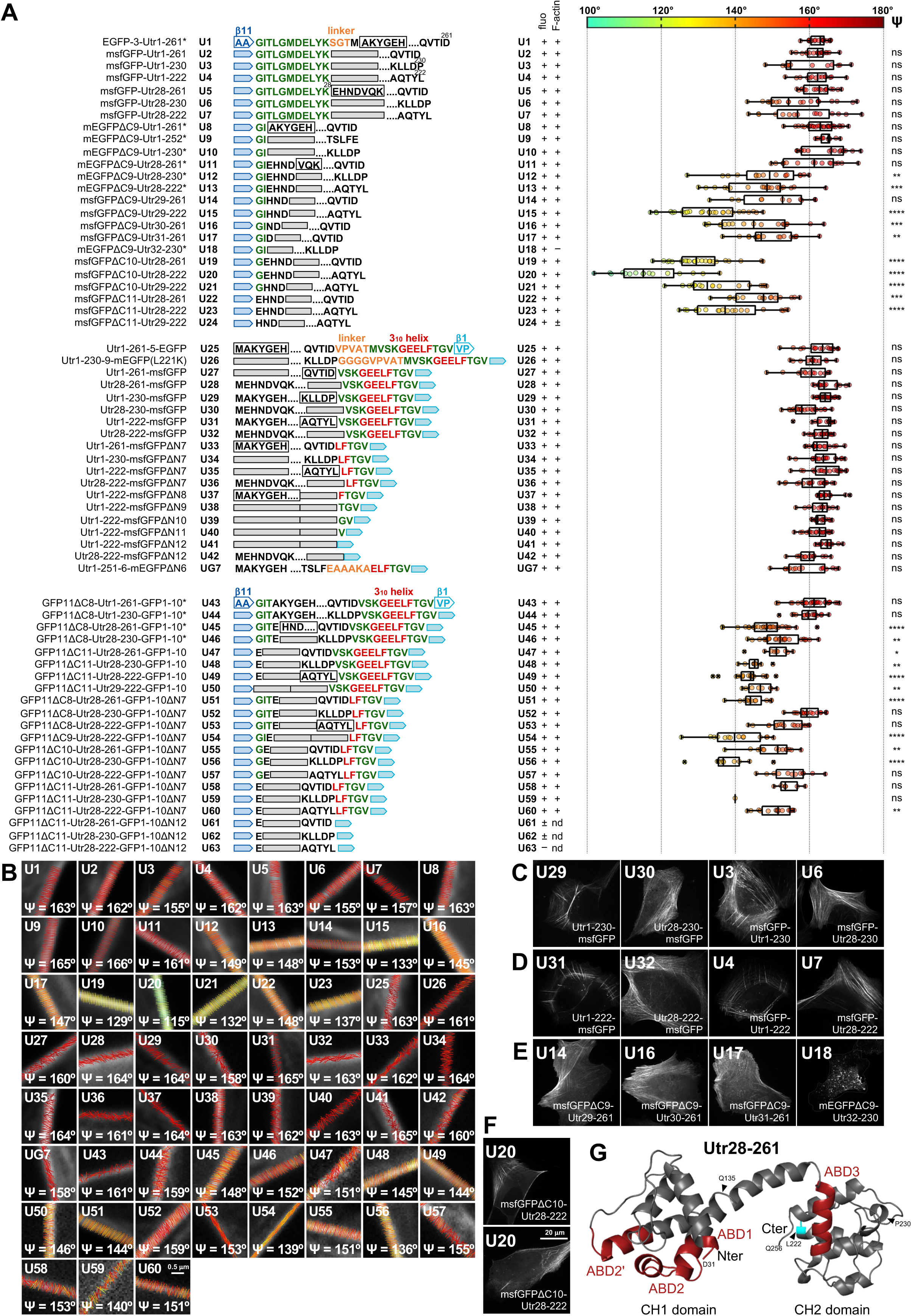
Engineering of Utrophin-based actin filament organization reporters for live-cell polarimetry, Related to. **Figure 2**. (A) List of Utrophin calponin homology domain (”Utr” for short)-GFP fusions tested for their usability in live polarimetry measurements in U2OS cells expressing each fusion. Fusion illustration and classification and box plots are as detailed in Figure S2A. The number of measurements (see methods for details), from top to bottom, are n= 18, 19, 13, 14, 17, 21, 15, 30, 11, 17, 16, 16, 20, 11, 26, 19, 18, 24, 16, 18, 16, 20, 17, 13, 14, 14, 25, 23, 13, 18, 16, 18, 19, 21, 15, 13, 17, 19, 14, 15, 16. 31, 17, 31, 27, 9, 8, 19, 8, 12, 17, 17, 14, 14, 10, 12, 6, 1, 15. The respective median ψ values are 163, 162, 155, 162, 163, 154, 157, 163, 165, 166, 161, 149, 149, 153, 133, 145, 147, 129, 115, 132, 148, 137, 163, 161, 160, 164, 164, 158, 165, 163, 162, 164, 164, 161, 164, 163, 162, 163, 165, 160, 158, 161, 159, 148, 152, 151, 145, 144, 146, 144, 159, 153, 139, 151, 136, 155, 153, 140, 151°. Statistical significance (right-most column) was obtained using a non-parametric Kruskal-Wallis test followed by a Dunn’s multiple comparisons test. The different constructs were compared to U1; ns=not significant (P>0.05); * P<0.05, ** P<0.01, *** P<0.001, **** P<0.0001. (B) Representative ψ stick maps on SFs from measurements in live cells expressing the indicated fusions, with mean ψ values indicated. The selected images correspond to median ψ values of the respective distributions. (C-E) Representative images of U2OS cells expressing the indicated reporters for assessing the contribution of specific Utrophin residues to actin binding. The localization of these reporters to SFs shows that removing the 27 N-terminal residues of Utrophin or/and shortening its C-terminus to Utr222 or Utr230 do not compromise actin binding (C,D). Proximity of C-terminally truncated GFP to Utr29-32 (E) can impair actin binding (U18). (F) Utrophin-based reporters localize both to SFs and to mitochondria: two different z-planes in the same cell show U20 on SFs (top plane) and mitochondria (bottom plane). (G) Ribbon representation of Utrophin structure showing the two calponin-homology (CH) domains and identified actin-binding sites (ABD) in red (PDB 1QAG) ^84,85^. Arrowheads point to specific residues. L222 and P230 relate to Utr222 and Utr230 fusions, respectively. Q135 points to the end of the CH1 domain, which is sufficient for actin binding ^84^.

**Figure S4.**
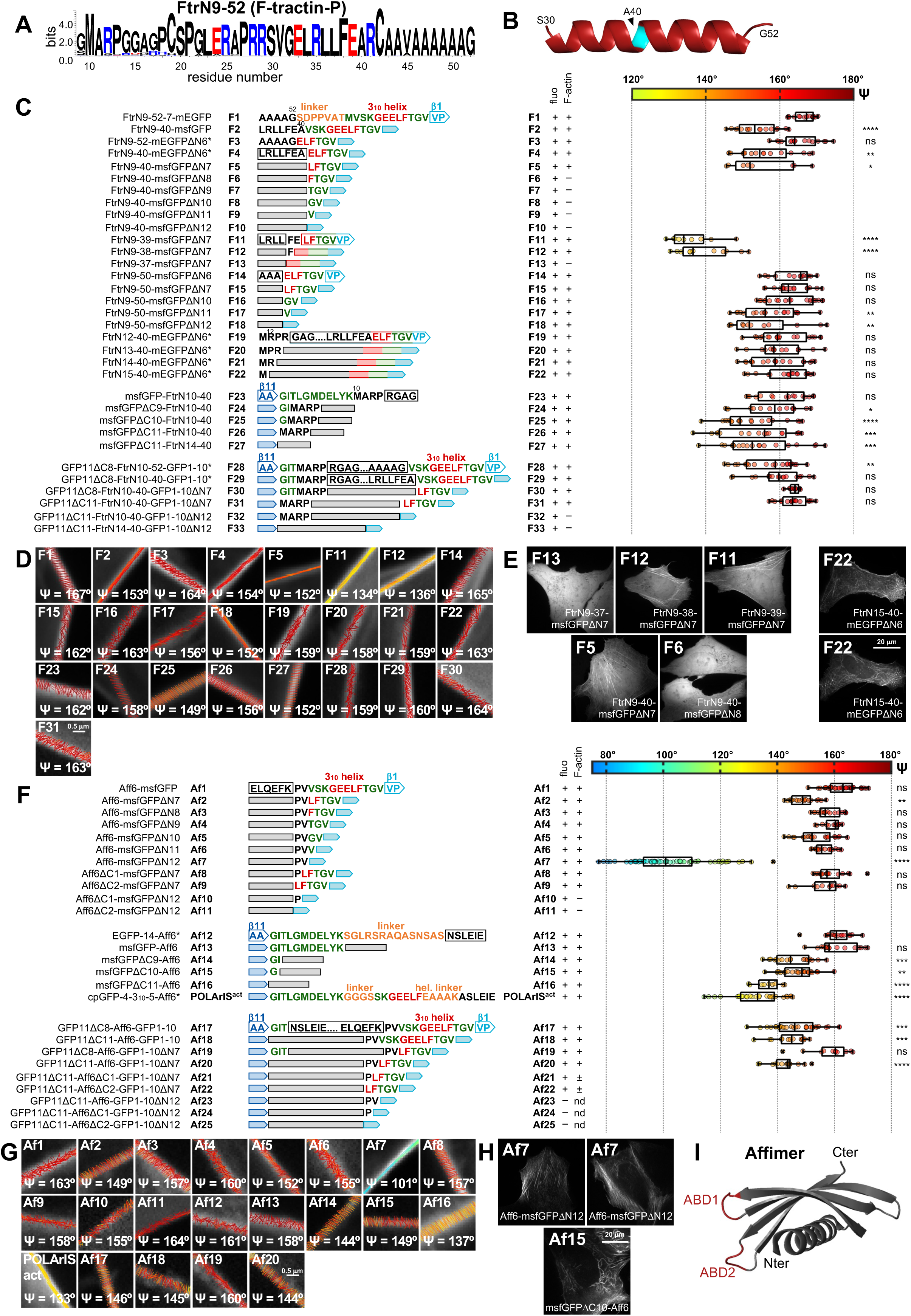
Engineering of F-tractin- and Affimer6-based actin filament organization reporters for live-cell polarimetry, Related to. **Figure 2**. (A-B) WebLogo3 representation illustrating the conservation of residues in the F-tractinN9-52 sequence (A). Sequences from 65 mammals were used for this representation. Negatively- and positively-charged residues are shown in red and blue, respectively. AlphaFold helix prediction for residues 30-52 is shown in (B), with A40 shown in cyan. A40 relates to F-tractinN9-40 fusions. (C) List of F-tractin (”Ftr” for short)-GFP fusions tested for their usability in live polarimetry measurements in U2OS cells expressing each fusion. Fusion illustration and classification and box plots are as detailed in Figure S2A. The number of measurements (see methods for details), from top to bottom, are n= 15, 16, 16, 16, 8, 9, 11, 14, 16, 19, 19, 12, 19, 14, 18, 18, 16, 14, 19, 17, 19, 23, 18, 11, 19. The respective median ψ values are 167, 153, 164, 154, 152, 134, 137, 165, 162, 163, 156, 152, 159, 158, 159, 163, 162, 159, 149, 156, 152, 159, 160, 164, 163°. Statistical significance (right-most column) was obtained using a non-parametric Kruskal-Wallis test followed by a Dunn’s multiple comparisons test. The different constructs were compared to F1; ns=not significant (P>0.05); * P<0.05, ** P<0.01, *** P<0.001, **** P<0.0001. (D) Representative ψ stick maps on SFs from measurements in live cells expressing the indicated F-tractin fusions, with mean ψ values indicated. The selected images correspond to median ψ values of the respective distributions. (E) Representative images of U2OS cells expressing the indicated reporters for assessing the contribution of specific F-tractin residues to actin binding. The localization of the reporters to SFs (diffuse cytosolic vs binding to SFs) shows that residues 37-40 are critical for actin binding. Removing the 14 N-terminal residues does not compromise actin binding (F22). F-tractin-based reporters localize both to SFs and to mitochondria (rightmost panels): two different z-planes in the same cell (F22) show F22 on SFs (top plane) and mitochondria (bottom plane). (F) List of Affimer6 (”Aff6” for short)-GFP fusions tested for their usability in live polarimetry measurements in U2OS cells expressing each fusion. Fusion illustration and classification and box plots are as detailed in Figure S2A. The number of measurements (see methods for details), from top to bottom, are n= 33, 19, 17, 14, 21, 17, 66, 17, 11, 7, 8, 16, 16, 19, 25, 9, 28, 30, 14, 11, 16. The respective median ψ values are 163, 149, 157, 160, 152, 155, 101, 157, 158, 155, 164, 161, 158, 145, 149, 137, 133, 146, 145, 160, 144°. Statistical significance (right-most column) was obtained using a non-parametric Kruskal-Wallis test followed by a Dunn’s multiple comparisons test. The different constructs were compared to Af12; ns=not significant (P>0.05); ** P<0.01, *** P<0.001, **** P<0.0001. (G) Representative ψ stick maps on SFs from measurements in live cells expressing the indicated Affimer6 fusions, with mean ψ values indicated. The selected images correspond to median ψ values of the respective distributions. (H) Affimer6-based reporters localize both to SFs (Af7, left cell) and to mitochondria (Af7, right cell and Af15). (I) Ribbon representation of the Affimer structure showing the two actin-binding sites (ABD) in red (PDB 4N6T) ^38,86^.

**Figure S5.**
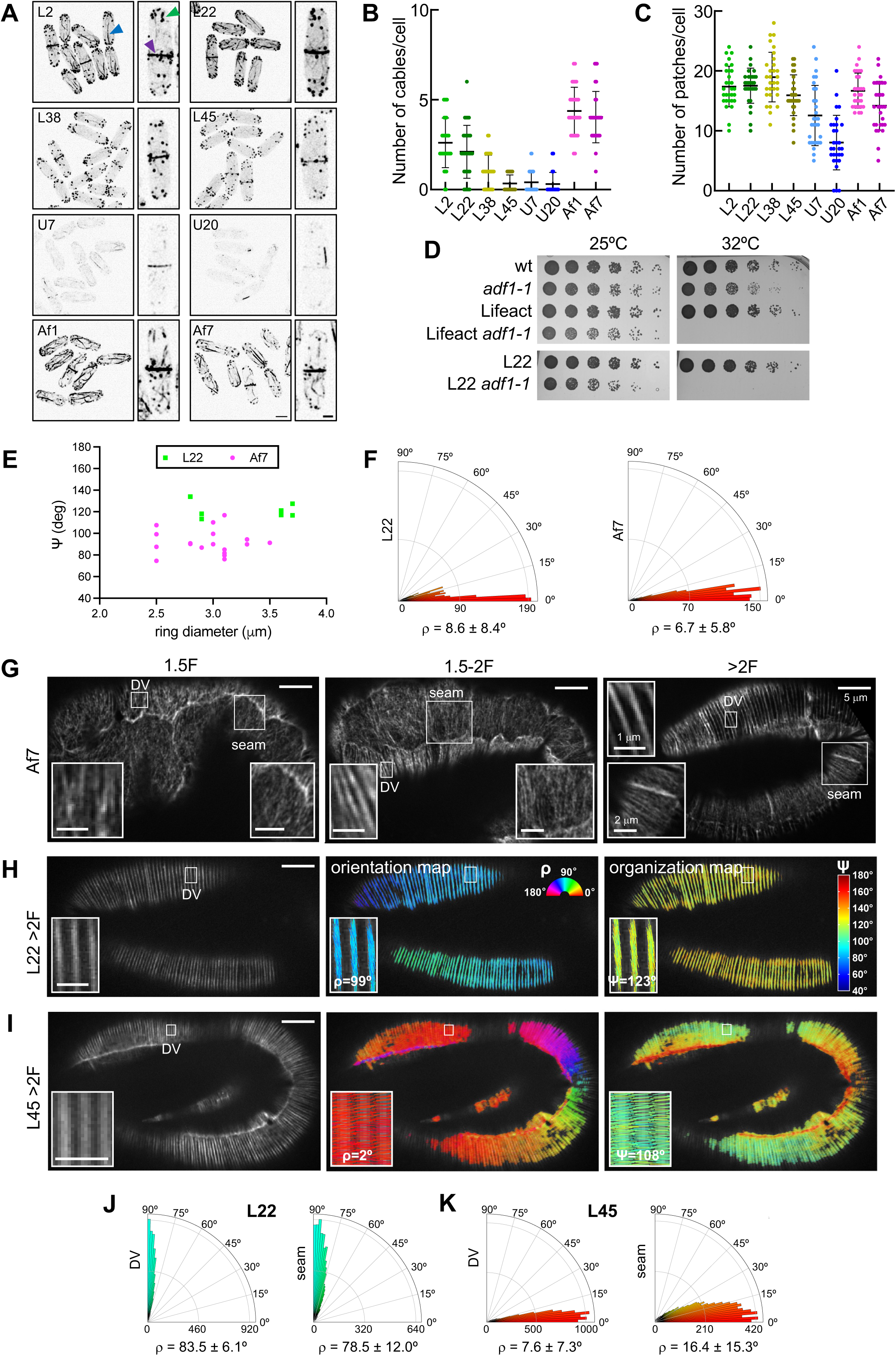
Polarimetry measurements of actin filament organization in live dividing fission yeast and live elongating *C. elegans* embryos expressing selected reporters, Related to. **Figures 3 and 4**. (A) Maximum intensity projection images of fission yeast cells expressing the corresponding actin reporter; scale bar, 4 µm. On the right hand of each panel, a magnified dividing cell is shown to observe details of the actin structures (patches, cables, ring) decorated by the actin reporter. Green, blue and magenta arrowheads point to a patch, cable and ring, respectively. Scale bar, 2 µm. (B) Quantification of actin cables per cell detected in the fission yeast strains expressing the corresponding actin reporter. Scatter plots show means ± SD. 30 cells were analyzed for each strain. The mean number of cables per cell for each strain are, from left to right: 2.6, 2.1, 1.0, 0.3, 0.4, 0.3, 4.4 and 4.0. (C) Quantification of actin patches per cell detected in the fission yeast strains expressing the corresponding actin reporter. Scatter plots show means ± SD. 30 cells were analyzed for each strain. The mean number of patches per cell for each strain are, from left to right: 17.4, 17.5, 19.0, 15.9, 12.6, 8.0, 16.7 and 14.2. (D) Serial dilution assay showing the genetic interaction between the cofilin mutant *adf1-1* and the expression of the shown reporters. (E) Mean actin filament alignment (ψ angle) in the cytokinetic ring of fission yeast expressing Af7 (magenta) or L22 (green) as a function of the constricting ring diameter. (F) Polar histograms of ρ value distributions in the cytokinetic ring of fission yeast expressing L22 (left) or Af7 (right). ρ values are represented with respect to the ring axis: considering that Af7 and L22 dipoles are parallel to actin filaments, the more parallel mean actin filament orientations are to the ring axis, the closer the angle values are to 0°. Means ± SD are shown. The number of cells is as in Fig.3H. (G) Summed intensity images of the respective polarimetry stacks shown in Fig.4 G, I, K. For all panels, anterior is to the left and dorsal is up. (H-K) Representative ρ (middle) and ψ (right) stick maps in >2-fold stage embryos expressing L22 (H) or L45 (I). The top images are summed intensity images of the respective polarimetry stacks. Insets show zoom-ins of selected ROIs (white outlined boxes). Mean ρ and ψ values are shown for each ROI. (J), (K), Polar histograms of ρ value distributions in DV and seam cells in >2-fold stage embryos expressing L22 (J) or L45 (K). ρ values are represented with respect to the DV/seam boundary: considering that L22 dipoles are parallel to actin filaments and that L45 dipoles are perpendicular to actin filaments, the more perpendicular mean actin filament orientations are to the boundary, the closer the angle values are to 90° (for L22) or to 0° (for L45) and the narrower the respective distributions. Means ± SD are shown. The number of embryos is as in Fig.4E. For all panels, anterior is to the left and dorsal is up.

**Figure S6.**
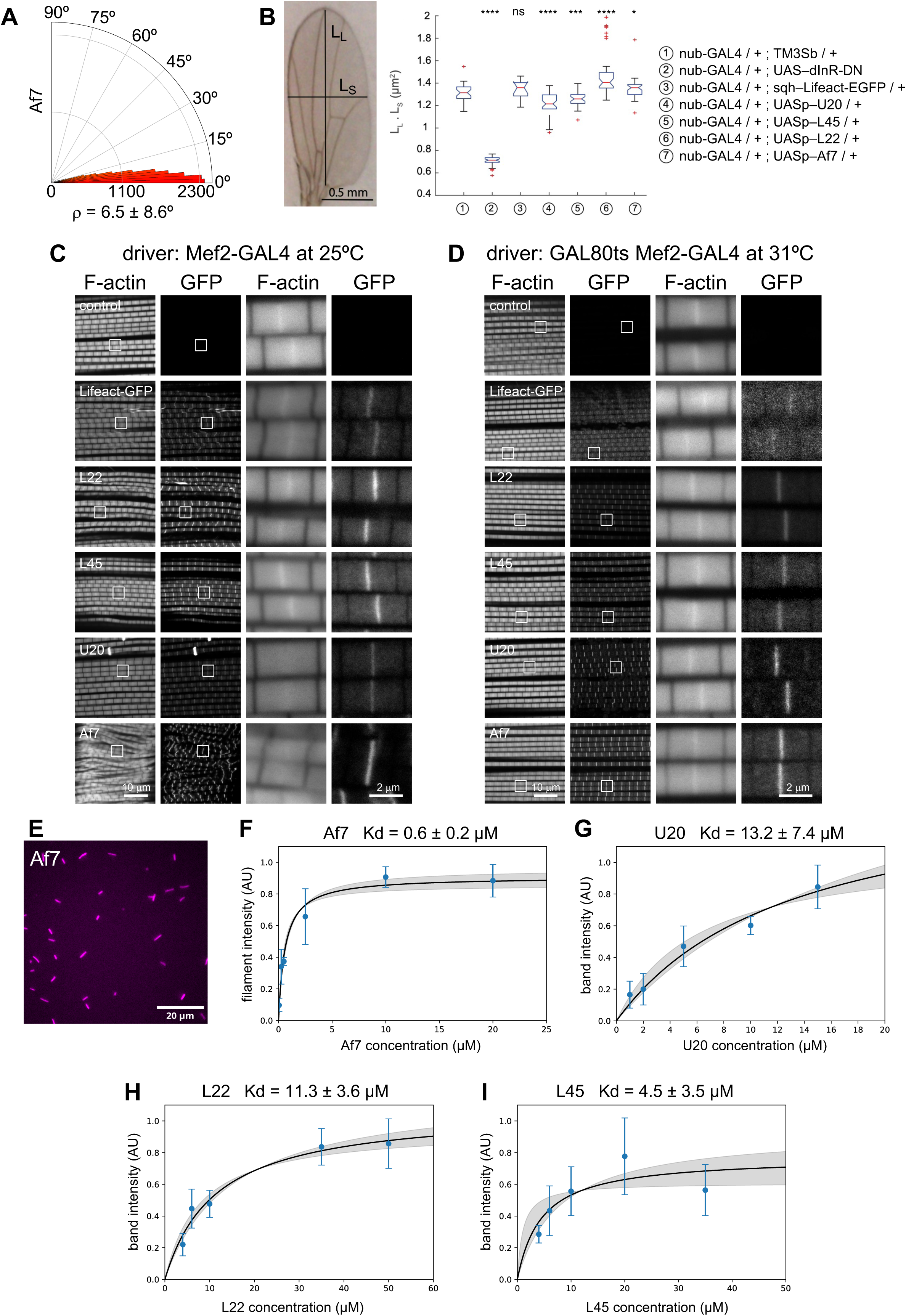
Functional characterization of actin organization reporters in *Drosophila* and binding affinites of best performing reporters to actin filaments. Related to. **Figure 5 and Table 1**. (A) Polar histogram of ρ value distributions in the actomyosin ring of a cellularizing *Drosophila* embryo expressing Af7. ρ values are represented with respect to the ring contour: considering that Af7 is parallel to actin filaments, the more parallel mean actin filament orientations are to the ring contour, the closer the angle values are to 0°. Mean ± SD is shown. (B) Effect of the expression of selected actin organization reporters on *Drosophila* wing growth. The image on the left displays an adult *Drosophila* wing and highlights the landmark points used for measuring the long axis (L_L_) and short axis (L_S_) of the wing (see methods for details). The product L_L_·L_S_ is utilized as a proxy for wing area. The accompanying graph shows box plots quantifying wing area for the shown genotypes. On each box, the central mark indicates the median, and the bottom and top edges of the box indicate the 25th and 75th percentiles, respectively. The whiskers extend to the most extreme data points not considered outliers, and the outliers are plotted individually using the ’+’ symbol. The table on the right shows the respective genotypes (see Key Resources Table for details): (1) serves as a positive control, (2) serves as a negative control for a perturbation resulting in reduced wing size (Insulin receptor dominant negative), (3) is a commonly used actin probe, and (4-7) represent the organization reporters described in this study. The number of wings for each genotype are, from left to right: 33, 40, 21, 46, 36, 37, 43. The respective median values are 1.31, 0.71, 1.36, 1.21, 1.26, 1.40, 1.36. A two-sample t-test was applied to evaluate statistical differences between each genotype and the positive control; ns=not significant, P>0.05; * P<0.05, *** P<0.001, **** P<0.0001. (C-D) Representative micrographs of phalloidin (”F-actin”) stainings of *Drosophila* flight muscles expressing selected actin organization reporters (”GFP”) as shown (see methods for details of genotypes). Expression was driven throughout flight muscle development (C), or transiently at the adult stage after muscle development (D). Insets show zoom-ins of selected sarcomeres (white outlined boxes). See also Figure 5. (E-I) Measurements of binding affinities of the best performing reporters characterized *in vivo* to actin filaments (see Table 1). The binding affinity of Af7 to filaments was performed by TIRF microscopy (E) in the presence of 0.6 µM actin and a range of Af7 concentrations. Fluorescence intensities of the GFP signal along actin filaments as a function of Af7 concentrations is shown in F. The graph depicts means ± SD from 3 independent experiments. Binding affinities of U20 (G), L22 (H) and L45 (I) to 2 µM actin filaments were performed by co-sedimentation assays. For all conditions, the graphs depict means ± SD from 3 independent experiments. The solid line is a fit of the data by a quadratic equation to derive a Kd value, with the shaded area representing the 95% confidence interval of Kd and plateau values.

**Figure S7.**
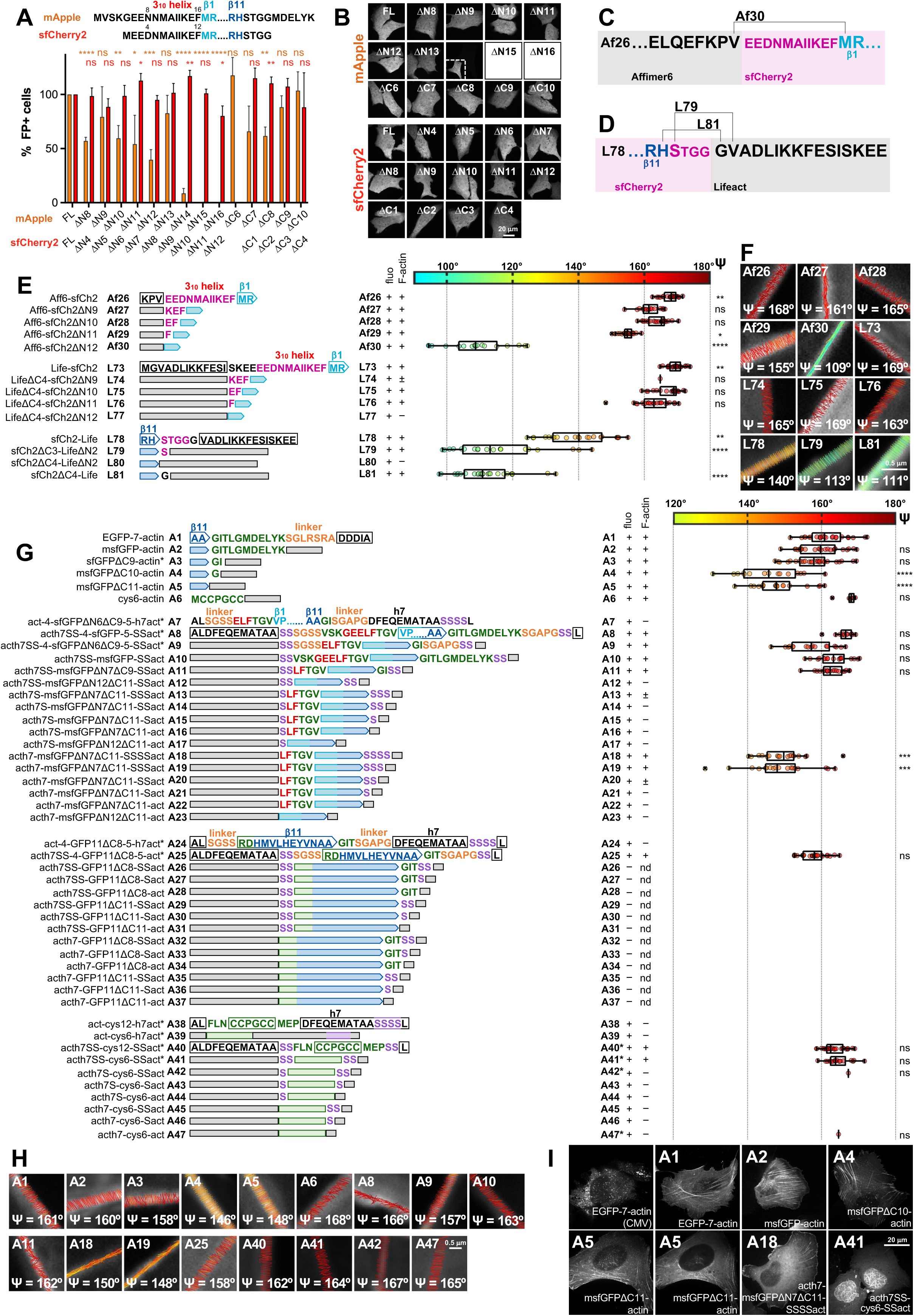
Engineering of G-actin- and red fluorescent protein-based actin filament organization reporters for live-cell polarimetry, Related to. **Figure 6**. (A) Top, amino acid sequence of the N- and C-termini of monomeric Apple (mApple) and superfolder Cherry2 (sfCherry2). The depicted secondary structure elements of mApple and sfCherry2 and color code are as in Fig.S1A. Residue numbering is as shown. Bottom, screening of N- and C-terminal truncation mutants of mApple and sfCherry2 using FACS. Bars (mean + SD) depict the measured percentages of fluorescence-positive (FP+) cells for full-length (FL), N-terminally (ΔN) and C-terminally (ΔC) truncated proteins. The mean values are, from left to right: 100, 100, 57, 99, 79, 89, 60, 99, 54, 113, 40, 95, 83, 102, 9, 117, 0.03, 101, 0.06, 80, 118, 66, 115, 62, 110, 88, 107, 104, 88. Data are from three independent experiments. Statistical significance was obtained using an unpaired t-test. The different constructs were compared to the respective FL; ns=not significant (P>0.05); * P<0.05, ** P<0.01, *** P<0.001, **** P<0.0001. (B) Screening of the same constructs used in (A) with spinning disk fluorescence microscopy. Representative images of live cells expressing each construct are shown. For the sake of comparison, images are displayed with the same intensity range. For weakly fluorescent cells, insets show contrast-enhanced images. No fluorescence was detectable for mAppleΔN15 and mAppleΔN16. (C-D) Engineering of red fluorescent protein-based actin filament organization reporters. Representative designs for constraining sfCherry2 mobility in Affimer6- (C) and Lifeact-based (D) fusions are illustrated for selected fusions, including for the best performing reporters (Af30, L81, see Table 1). The full screen is shown in panel E. (E) List of Affimer6- and Lifeact-based sfCherry2 (”sfCh2” for short) fusions tested for their usability in live polarimetry measurements in U2OS cells expressing each fusion. sfCherry2 fusions were either C-terminal (Af26-Af30, L73-L77), or N-terminal (L78-L81). Fusion illustration and classification and box plots are as detailed in Figure S2A. The number of measurements (see methods for details), from top to bottom, are n= 23, 16, 23, 25, 18, 23, 1, 22, 22, 22, 20, 25. The respective median ψ values are 168, 161, 165, 155, 109, 170, 165, 169, 163, 140, 113, 111°. Statistical significance (right-most column) was obtained using a non-parametric Kruskal-Wallis test followed by a Dunn’s multiple comparisons test. The different Affimer6 constructs were compared to Af12; the different Lifeact constructs were compared to L1; ns=not significant (P>0.05); * P<0.05, ** P<0.01, **** P<0.0001. (F) Representative ψ stick maps on SFs from measurements in live cells expressing the shown Affimer6-based (Af) and Lifeact-based (L) sfCherry2 fusions, with mean ψ values indicated. The selected images correspond to median ψ values of the respective distributions. (G) List of human β-actin (”act” or “actin” for short) fusions tested for their usability in live polarimetry measurements in U2OS cells expressing each fusion. The fluorophores used were GFP, the β11 strand alone (”GFP11”), or tetracysteine peptides (”cys6” or “cys12”) as shown in Fig.6A. For G-actin terminal fusions, GFP or tetracysteine peptides were fused to the N-terminus of G-actin (A1-A6). For G-actin intramolecular fusions, msfGFP (A7-A23), the β11 strand alone (A24-A37), or tetracysteine peptides (A38-A47) were placed intramolecularly within the G-actin structure (see Fig.6B). Fusion illustration and classification and box plots are as detailed in Figure S2A. Fusions A6, A40 and A41 localized both to SFs and to nuclear F-actin, and box plots include measurements from both F-actin pools. Measurements for fusions A42 and A47 are from nuclear F-actin. The number of measurements (see methods for details), from top to bottom, are n= 23, 24, 26, 15, 18, 6, 12, 17, 17, 19, 18, 18, 26, 26, 17, 1, 1. The respective median ψ values are 161, 160, 158, 146, 148, 168, 166, 157, 163, 162, 150, 148, 158, 162, 164, 167, 165°. Statistical significance (right-most column) was obtained using a non-parametric Kruskal-Wallis test followed by a Dunn’s multiple comparisons test. The different constructs were compared to A1; ns=not significant (P>0.05); *** P<0.001, **** P<0.0001. (H) Representative ψ stick maps on SFs from measurements in live cells expressing the indicated G-actin fusions, with mean ψ values indicated. ψ stick maps for A42 and A47 are from nuclear F-actin. The selected images correspond to median ψ values of the respective distributions. (I) Representative images of U2OS cells expressing the indicated reporters for assessing the contribution of linkers and shortened GFPs to their localization to specific actin populations. Expression is driven by a CMV_trunc_ promoter: the widely used full-strength CMV leads systematically to aggregation (leftmost panel). The absence of linkers and the proximity of GFP to the N-terminus of G-actin do not compromise binding to F-actin but result in lower enrichment of the reporters in myosin-II containing actin pools. G-actin-based reporters localize both to SFs and to mitochondria: two different z-planes in the same cell (A5) show A5 on SFs (left plane) and mitochondria (right plane). Fusions with tetracysteine peptides localized also to nuclear F-actin (A41).

## STAR Methods

### EXPERIMENTAL MODEL AND SUBJECT DETAILS

#### Cell lines and cell culture

U2OS osteosarcoma cells were used for the screening of actin organization reporters with respect to their localization and usability for polarimetry measurements. U2OS cells were from ATCC (HTB-96). Cells were maintained in McCoy’s medium (Thermo Fisher, Cat#16600082) supplemented with 10% fetal bovine serum (Dominique Dutscher, Cat# S181H), 100 U/mL penicillin and 100 μg/mL streptomycin antibiotics (Sigma-Aldrich, Cat#P4333) in a humidified atmosphere at 37°C containing 5% CO_2_. Transfections were performed 16 h prior to live imaging using FuGENE HD Transfection Reagent (Promega, Cat#E2311), following the manufacturer’s instructions. To obtain single cells for imaging, 25×10^3^ U2OS cells were typically seeded into a 24-well glass bottom plate (Cellvis, Cat#P24-1.5H-N) a day prior to the day of transfection, for allowing an optimal number of cells to attach and spread. A total of 0.2 μg of DNA and a 4:1 ratio of FuGENE HD (μL) : DNA (μg) were used per reaction. To minimize overexpression, the amount of DNA for pCMV plasmids was reduced to 50 ng, leading to a 16:1 ratio of FuGENE HD (μL) : DNA (μg). Cells were imaged 16h post-transfection.

HeLa cells were used for the characterization of intramolecular GFP (iGFP)-beta and - gamma actin fusions. Stable HeLa cell lines with regulated expression of either iGFP-beta actin or iGFP-gamma actin were generated with the Flp-In system (Life Technologies) using HeLa cells that contained a single FRT site according to the manufacturer’s instructions ^87^. The resulting cell lines were cultured in Dulbecco’s Modified Eagle Medium (Wisent Bio Products, Cat#319-030 CL) supplemented with 10% fetal bovine serum (Wisent Bio Products, Cat#080-650), 1% penicillin/streptomycin (Invitrogen Cat#15140122), 5 μg/mL blasticidin (BioShop Canada Inc, Cat#BLA477), and 2 μg/mL puromycin (BioShop Canada Inc, Cat#PUR333). Expression of GFP fusion proteins was induced by addition of 0.25 μg/mL doxycycline to the growth media for 24 h before either fixation or harvesting. Cells were maintained in a Forma Series II incubator (Thermo Scientific) at 37°C in a 5% CO_2_ atmosphere.

#### Fission yeast strains, maintenance and genetics

Standard *Schizosaccharomyces pombe* media and genetic manipulations were used ^88^. All strains used in the study were isogenic to wild-type 972 and their genotypes are described in Table S3. The generation of transgenic strains is described in the method details section. Strains from genetic crosses were selected by random spore germination and replica in plates with appropriate supplements or drugs. Transformations were performed using the lithium acetate-DMSO method as described in ^89^. Drop assays (Figures 3D,E and S5D) were performed by serial dilutions of 1/4 from a starting sample of optical density of 1.0 of the indicated strains, which were plated on YE5S medium supplemented with the corresponding drug and incubated for 3 days at 28°C unless stated differently.

#### *C. elegans* strains, maintenance and genetics

Bristol strain N2 was used as the wild-type strain. *C. elegans* strains used in this study and their genotypes are listed in the Key Resources Table and were reared using standard methods ^90^. The generation of transgenic strains is described in the method details section. The strains were grown at 20°C and fed Escherichia coli OP50. The EG6699 (unc-119(ed3) III) strain, used as the host strain of FBR193, FBR195 and FBR196 strains generated in this study, was grown at 15°C and fed E. coli HB101 before injection ^91^.

#### *Drosophila* strains, maintenance and genetics

*Drosophila melanogaster* strains used in this study and their genotypes are listed in the Key Resources Table. The generation of the transgenic strains UASp–L22, UASp–L45, UASp–U20, and UASp–Af7 is described in the method details section. Fly stocks were grown and maintained at 25°C on semi-defined medium (https://bdsc.indiana.edu/information/recipes/germanfood.html). The GAL4/UASp expression system was used to drive expression in the *Drosophila* embryo using GAL4 expressed under the control of the maternal alphaTub67C promoter (mat–α-tub–GAL4) (BDSC_80361). Crosses were maintained at 25°C. The GAL4/UAS expression system was used to drive the expression of actin organization reporters in *Drosophila* indirect flight muscles. Mef2-GAL4 (BDSC_27390) or tub-GAL80ts ; Mef2-GAL4 (BDSC_7108, BDSC_27390) females were crossed to males of the following genotypes: w[1118] (BDSC_3605), UAS–GFP-GMA (BDSC_31776), UAS–Lifeact-EGFP (BDSC_35544), UASp–L22, UASp–L45, UASp–U20, and UASp–Af7. Crosses with the Mef2-GAL4 driver were grown at 25°C. Crosses with the tub-GAL80ts ; Mef2-GAL4 driver were grown at 18°C (no GAL4 activity): a few days after eclosion, adults were transferred to a permissive temperature of 31°C for 5 days prior to flight tests or fixation/staining or to a permissive temperature of 25°C for 5 days prior to live polarimetry. The GAL4/UAS expression system was used to drive the expression of actin organization reporters in the *Drosophila* wing. nub-GAL4 (BDSC_86108) females were collected within 2 days and combined to create a uniform population. Eight of these females were crossed to three males of the following genotypes: UAS–dInR-DN (BDSC_8253), sqh*–*Lifeact-EGFP, UASp–L22, UASp–L45, UASp–U20, and UASp–Af7. The rearing temperature was maintained at 25°C, and the tubes were flipped daily.

### METHOD DETAILS

#### Generation of mammalian expression plasmids for screening actin organization reporters

All constructs were designed in silico with SnapGene (Dotmatics) and are listed in Table S1. To drive expression of the constructs in mammalian cells, we used the immediate early enhancer and promoter of human cytomegalovirus (CMV promoter, 508 base pairs), as well as a truncated version (CMV_trunc_, 54 base pairs) for low-level expression; the latter was originally generated for reduced expression of EGFP-beta-actin ^92^. Addgene plasmids #31502 and #54759 were used to obtain the CMV_trunc_ and CMV backbones, respectively. All constructs screened for actin organization reporters were driven by CMV_trunc_ apart from the ones labeled with an asterisk in Figures S2A, S3A, S4C,F, S7G and the iGFP-beta- and -gamma constructs (Figure 6D-R) which were driven by CMV. Fluorescent protein constructs screened for fluorescence in the truncation screens (Figures S1C,D and S7A,B) were driven by CMV. Lifeact-mEGFP and EGFP-beta-actin cDNAs were a gift from Yannick Hamon (CIML, France). FtrN9-52-mEGFP and EGFP-Affimer6 cDNAs were a gift from John Hammer (NIH/NHLBI, USA) and Michelle Peckham (University of Leeds, UK), respectively. Beta- and gamma-actin cDNAs were a gift from Boris Hinz (University of Toronto, Canada). Synthetic genes for sfGFP, msfGFP, human beta-actin and GFP11ΔC8-FtrN10-52-GFP1-10 were from Eurofins Genomics (Germany). Fluorescent protein fusions were generated using monomeric (A206K) EGFP (mEGFP) ^93,94^, monomeric (V206K) superfolder GFP (msfGFP) ^93,95–97^, monomeric Apple (mApple) ^96,98^ and superfolder Cherry2 (sfCherry2) ^76^. To optimize the intramolecular self-association of β11 with the GFP1-10 moiety in our circular permutants, we have used the sfGFP-evolved sequences, GFP1-10 OPT and GFP11 M3, which have been optimally engineered to work in bipartite split-GFP complementation assays ^99^. All constructs were generated with seamless cloning (In-Fusion HD Cloning Plus Kit from Takara Bio, Cat. # 638910) using NheI/BamHI (or AflII/BamHI for iGFP constructs) linearized plasmid backbones and the oligonucleotide primer sequences listed in Table S2. Primers were Cloning Oligo (<60 bp) or EXTREmer (>60 bp) synthesis and purification quality from Eurofins Genomics (Germany). Restriction enzymes were FastDigest enzymes from Thermo Scientific. All plasmids were verified by sequencing (Eurofins Genomics, Germany) after each cloning step.

We note the following with respect to residue numbering of EGFP/sfGFP in our study: although the valine following the initiating methionine is typically numbered 1a to maintain correspondence between EGFP/sfGFP and wild-type GFP numbering ^100^, we number this valine as 2 in this study to facilitate the naming of N-terminal truncations in the screen. As a result, the last residue of EGFP/sfGFP is 239, an N-terminally truncated msfGFP mutant missing the first six residues (ΔN6) starts with ELFTGV…, and a C-terminally truncated msfGFP mutant missing the last nine residues (ΔC9) ends with …AAGI.

#### Screening of actin organization reporters in live U2OS cells

##### Confocal fluorescence microscopy and image processing

For live cell imaging, right before microscopy and due to the absence of CO_2_ control on our microscope setup, the culture medium was exchanged by Leibovitz medium (Thermo Fisher, Cat#21083027) supplemented with 10% fetal bovine serum and antibiotics. Cells were kept at 37°C in a heating chamber (OkoLab, Cat#H301-TUNIT-BL). Fluorescence images were acquired using a custom spinning disk microscope (detailed in the Polarimetry methods section) with a Nikon Plan Apo ×100/1.45 NA oil immersion objective lens, 488-561- and 641-nm laser lines and an iXon Ultra 888 EMCCD camera. Z-stacks were acquired with a Δz interval of 0.5 μm. Exposure times were in the range of 0.5–2.0 s depending on the exact condition.

Images were processed with the open-source image processing software ImageJ/Fiji. The images displayed in Figure S1D and Figure S7B are maximum intensity projections of two consecutive z-planes displayed with the same intensity range to allow for intensity comparison between the FP truncation mutants. All the other shown images are maximum intensity projections of two consecutive z-planes contrasted manually in order to optimize the image display.

##### Polarimetry measurements in live U2OS cells

Polarimetry stacks using 36 polarization angles were recorded in focal planes containing peripheral stress fibers with a typical exposure time of 0.1-0.2 s per polarized image (see details for the optical setup and signal processing in the Polarimetry methods section). Typically, a minimum of five fields of views containing single cells was acquired per experimental condition. Polarimetry stacks were systematically registered using the StackReg plugin for ImageJ to correct for x and y axis drift during acquisition. To select peripheral SF-associated pixels for analysis, binary masks of SF segments were generated using the open source tool FilamentSensor 0.2.3 ^101^, freely available at http://www.filament-sensor.de/. A pre-processing tab in the FilamentSensor software requires adjustment in the contrast and removal of standalone pixels, followed by the optional application of filters. A standard and optimized preprocessing was using Laplace filter, 8 neighbors and factor 4; Gaussian filter, sigma 1; Cross correlation filter, size 10 and zero 30%; and a directed Gaussian filter, sigma 8. The binarization method chosen was by area, and filament detection parameters were typically chosen as follows: minimum mean value 25, sigma 2, minimum standard deviation 5, minimum filament length 20, minimum angle difference 20, tolerance 5%. The final selection was done manually, and only identified filaments that were colocalizing with peripheral SFs were used to generate the binary masks for selecting the pixels for polarimetry analysis.

Polarimetry data were analyzed according to the framework defined by ^34^ to obtain the ρ and ψ angle per image pixel. Analysis and data representation, including color-coded stick representations of the measured angles per pixel were done with the Polarimetry software which is a Matlab App Designer standalone application. The source code and desktop app are available at https://github.com/cchandre/Polarimetry.git. The Matlab-based Polarimetry software is the precursor of the Python-based app PyPOLAR used for the analysis of the yeast, *Drosophila* and *C. elegans* polarimetry data (see respective methods sections). The distributions of the ψ angles are represented in box plots with overlaid data points. Each data point represents a single actin fiber. If more than one fibers were identified in the same field of view, measurements for each fiber are shown as distinct datapoints, which results in more than one measurements per field of view. On each box, the central mark indicates the median, and the left and right edges of the box indicate the 25th and 75th percentiles, respectively. The whiskers extend to the most extreme data points not considered outliers. Boxplots were generated with custom-written Matlab code (see *Data post-processing* in https://github.com/cchandre/Polarimetry.git). The number of measurements for each construct, the respective median ψ values and the statistical test used in GraphPad Prism to evaluate differences are mentioned in the respective legend.

#### Polarimetry measurements in fixed U2OS cells

U2OS cells were fixed for 20 min with 4% paraformaldehyde (Electron Microscopy Sciences, Cat#15714) in 37°C-prewarmed cytoskeleton buffer (10 mM MES pH 6.1, 150 mM NaCl, 5 mM EGTA, 5 mM MgCl_2_, 5 mM glucose), followed by 2 × 5 min wash steps in phosphate-buffered saline (PBS) solution. Cells were subsequently incubated with 0.165 μM Alexa Fluor 488-phalloidin (Thermo Fisher Scientific, Cat# A12379) or 0.165 μM SiR-actin (Spirochrome, Cat#SC006) in PBS containing 0.1% saponin and 1% IgG-free/protease free bovine serum albumin (Jackson ImmunoResearch, Cat#001-000-161) for 1 h at RT. Coverslips were mounted with 15 μL Fluoromount (F4680; Sigma-Aldrich) for image acquisition.

Polarimetry stacks using 18 polarization angles were recorded in focal planes containing ventral and peripheral stress fibers with a typical exposure time of 0.1-0.2 s per polarized image. The pixels of the stress fibers for analysis were selected by a combination of intensity thresholding and manual selection of the region to analyze. Analysis and data representation, including color-coded stick representations of the measured angles per pixel were done with Polarimetry or PyPOLAR softwares. The distributions of the ψ angles are represented in box plots with overlaid data points as described above for measurements in live cells. The number of measurements for each dye and the respective median ψ values are mentioned in the respective legend.

#### Flow cytometry analysis of truncation mutants of fluorescent protein variants

U2OS cells were seeded at a density of 5×10^4^ cells per well in 24-well plates. After 24 h, cells were transfected with 0.5 μg of corresponding plasmid DNAs in 50 μL of Jet Prime Buffer mixed with 1 μL of jetPRIME reagent (Polyplus, Cat#101000046). Twenty-four hours after transfection, cells were trypsinized, resuspended in PBS supplemented with 2% fetal bovine serum, then transferred in 96-well conical bottom plates. Cells were fixed with 3.7% paraformaldehyde for 10 min and resuspended in 1% BSA, PBS buffer. Green (mEFP, msfGFP) and red (mApple, sfCherry2) fluorescence were collected on 10,000 cells using a MACSQuant VYB flow cytometer (Miltenyi Biotec). The fluorescence threshold was defined based on the background fluorescence of untransfected U2OS cells. The percentage of green and red positive fluorescence was analyzed with FlowJo® software (BD Biosciences). The fluorescence of full-length constructs was normalized to 100% for each independent experiment. Bar graphs of the measured fluorescence were prepared with GraphPad Prism. The mean values and the statistical test used to evaluate differences are indicated in the respective legend.

#### Characterization of intramolecular GFP (iGFP)-beta and -gamma actin fusions

##### siRNA treatment and rescue experiments

HeLa cells grown to 40% confluency in 6-well plates were transfected with 100 pmol double-stranded siRNA targeting *ACTB* (sequence #1:

AAAUAUGAGAUGCGUUGUUACAGGA; sequence #2: UCCUGUAACAACGCAUCUCAUAUUUGG) or *ACTG1* (sequence #1: GCAUGGGUUAAUUGAGAAUAGAAAT; sequence #2: AUUUCUAUUCUCAAUUAACCCAUGCAG) using Lipofectamine 2000 (Invitrogen, Cat#11668019) following manufacturer’s instructions. For rescue experiments, siRNA-resistant transgenes were expressed 24 h after siRNA transfection and either fixed or harvested 48 h post-siRNA transfection. All siRNAs were obtained from IDT (Integrated DNA Technologies).

##### Cell harvesting, SDS-PAGE, and western blotting

HeLa cells grown to near confluency in 6-well dishes were harvested by scraping in 100 μL RIPA lysis buffer (50 mM Tris pH 7.4, 150 mM NaCl, 1% NP-40, 0.5% sodium deoxycholate, 0.1% SDS, 1 mM PMSF, 2 μg/mL aprotinin, 2 μg/mL leupeptin) on ice. Lysates were spun at 20,800 *g* for 20 min, after which supernatants were collected, mixed with SDS-PAGE sample buffer, and boiled at 95°C for 5 min. Samples were run on a 10% polyacrylamide gel and subsequently transferred onto nitrocellulose membranes (Bio-Rad). Membranes were blocked for 1 h in 5% skim milk powder in TBST (TBS with 0.0025% Tween-20), before incubation with either mouse anti-gamma actin antibody (Bio-Rad, Cat#MCA5776GA; dilution 1:200), mouse anti-beta actin antibody (Bio-Rad, Cat#MCA5775GA; dilution 1:200), or mouse anti-acetylated alpha tubulin antibody (Santa Cruz, Cat#sc-23950; dilution 1:2000) for 1 h. Following secondary antibody incubation, membranes were developed with chemiluminescent solutions (Thermo) for 1-2 min at room temperature and visualized using a Bio-Rad MP Imager (Bio-Rad). Actin isoform band intensities were measured with ImageLab software (Bio-Rad, Canada). Intensities of actin isoform bands were first normalized to respective tubulin band intensities. They were then divided by the normalized intensity of respective uninduced control actin isoform bands to give the presented normalized actin isoform band intensities.

##### Multinucleation and mitotic staging assays

Stable iGFP-actin HeLa cells were seeded onto glass coverslips in 6-well dishes. At roughly 40% confluency, cells were transfected with 100 pmol of either control, beta, or gamma actin-targeting siRNA (Integrated DNA Technologies) with Lipofectamine 2000 (Invitrogen) following manufacturer’s instructions. Where indicated, 24 h post-transfection cells were induced to express either iGFP-beta or -gamma actin by the addition of 0.25 μg/mL doxycycline. Cells were fixed 48 h post-transfection as described below. For multinucleation assays, cells were classified as either mono- or multi-nucleate by manually scoring the number of nuclei, as reported by Hoechst staining, within each cell boundary, as reported by acetylated alpha-tubulin staining, omitting cells fixed mid-division. For mitotic staging experiments, cells were manually classified as mitotic if the following features were observed: i) condensed chromosomes by Hoechst staining and/or b) an intercellular bridge as reported by acetylated alpha-tubulin staining. Early mitotic cells were further classified into either ‘prophase’ or ‘metaphase’ populations based on the organization of their condensed chromosomes, with ‘metaphase’ cells exhibiting sharp alignment with the metaphase plate, and ‘prophase’ cells exhibiting chromosomal rosettes or otherwise unaligned chromosomes. Data was entered into GraphPad Prism to generate bar graphs and perform statistical tests; the number of cells scored for each condition and details of statistical tests performed are described in the respective figure legends.

##### Immunofluorescence

To visualize iGFP-tagged actins, HeLa cells were fixed with 3.7% paraformaldehyde in PHEM buffer (60 mM PIPES pH 7.0, 25 mM HEPES, 10 mM EGTA, 4 mM MgSO_4_.7 H_2_O) for 10 min at room temperature and permeabilized by 0.2% Triton-X-100 in PBS for 10 min. Coverslips were blocked for 1 h in 3% BSA in PBS. Where indicated, iGFP-expressing cells were probed with anti-vinculin antibody (Sigma-Aldrich, Cat#V9131, 1:100) for 16 h at 4°C. Coverslips were subsequently incubated with either Alexa 594 or Alexa 647 conjugated goat-anti mouse secondary antibody (Invitrogen, 1:400) for 1 h.

For experiments visualizing endogenous beta actin and either iGFP gamma actin or vinculin, cells were fixed with 3.0% paraformaldehyde in PHEM buffer for 30 min at 37°C, followed by a second fixation for 5 min in -20°C methanol. Coverslips were blocked with 3% BSA in PBS overnight at 4°C, before incubation for 1 h with mouse anti-beta actin antibody (Bio-Rad, Cat#MCA5775GA; dilution 1:600), either with or without anti-vinculin antibody (Sigma-Aldrich, Cat#V9131, 1:100), diluted in 1% BSA in PBS. Coverslips were subsequently incubated with either Alexa 594 or Alexa 647 conjugated goat-anti mouse secondary antibody (Invitrogen, 1:400) for 1 h.

For multinucleation and mitotic staging experiments, cells were fixed with -20°C methanol for 10 min, blocked with 3% BSA in PBS, and subsequently stained with mouse anti-acetylated alpha tubulin antibody (Santa Cruz, sc-23950, 1:400) for 1 h. Coverslips were then incubated with Alexa 594 conjugated goat-anti mouse secondary antibody (Invitrogen, 1:400) for 1 h.

Prior to mounting with Mowiol (Polyvinyl alcohol 4-88, Fluka), coverslips were incubated in 1 μg/mL Hoechst 33258 (Sigma) for 10 min and rinsed in ddH_2_O. Cells were visualized with either a PerkinElmer UltraView spinning disk confocal scanner mounted on a Nikon TE2000-E with a 60x/1.4 NA oil-immersion objective lens and 1.515 immersion oil at room temperature or a Leica SP8 scanning confocal microscope with a 63x/1.4 NA oil-immersion objective lens and Leica Type F immersion oil. Images were acquired using METAMORPH software (v.7.7.0.0; Molecular Devices) driving an electron multiplying charge-coupled device (CCD) camera (ImagEM, Hammamatsu) or LAS-X software (v.1.4.4; Leica) driving HyD detectors. Z sections (0.2 μm apart) were acquired to produce a stack that was then imported into AutoQuant X3 (Media Cybernetics) for 3D deconvolution (5 iterations). Single Z-slices were generated in ImageJ (v2.1.0). Images were overlaid in Adobe Photoshop (v23.0.2) involving adjustments to brightness and contrast.

#### Characterization of fission yeast strains expressing actin organization reporters

##### Generation of fission yeast strains

msfGFP-tagged actin organization reporters were expressed from the fission yeast *leu1*+ locus under the control of the *cdc42*+ promoter using the integrative vector pJK148 ^102^. Briefly, around 500 bp from the *cdc42*+ promoter were amplified by PCR and cloned into the pJK148 vector using the *Sac*I and *Xba*I sites, creating pSRP12. The *adh1*+ terminator was amplified by PCR and cloned into pSRP12 using the *BamH*I and *Sal*I sites, creating pSRP14. Finally, the fragments coding for each of the actin reporters fused to msfGFP were obtained by digestion with *Nhe*I and *BamH*I from the respective mammalian expression plasmids and cloned into pSRP14, between the *cdc42*+ promoter and the *adh1*+ terminator, creating the plasmids pSRP16 to pSRP23, respectively (see Table S1). All oligos used are listed in Table S2. Plasmids were linearized by *Nru*I digestion, before transformation of a wild-type strain. Genetic crosses were performed to combine the actin reporter-expressing strains to strains expressing the proper marker to check cytokinesis dynamics, microtubule organization or to the profilin or cofilin mutant thermosensitive strains, *cdc3-319* and *adf1-1*, respectively (see Table S3).

##### Microscopy and image analysis

For imaging, fission yeast cells were grown at 28°C (32°C for cells shown in Figure 3A) in YE5S medium to exponential growth. For time-lapse imaging, 300 µL of early log-phase cell cultures were placed in a well from a µ-Slide 8 well (Ibidi, Cat#80821) previously coated with 10 µL of 500 µg/mL soybean lectin (Sigma-Aldrich, Cat#L1395). Cells were left for 1 min to attach to the bottom of the well and culture media was removed carefully. Then, cells were washed three times with the same media and finally 300 µL of fresh media were added ^103^, before incubation in the microscope chamber at the same temperature at which cells had been cultured.

Time-lapse images shown in Figure 3B are maximum intensity projections obtained from z-stacks of 7 slices at 1 µm interval every 2 minutes, acquired using an Olympus IX81 spinning disk confocal microscope with Roper technology controlled by Metamorph 7.7 software (Molecular Devices), equipped with a 100X/1.40 Plan Apo oil lens, a Yokogawa confocal unit, an EVOLVE CCD camera (Photometrics) and a laser bench with 491-561 nm diode. Exposure time for green or red channels was 0.5 s.

Time-lapse images shown in Figure 3A are maximum intensity projections obtained from z-stacks of 7 slices at 0.3 µm interval every 6 minutes, acquired using a Dragonfly 200 Nikon Ti2-E spinning disk confocal microscope controlled by Fusion software (Andor), equipped with a 100X/1.45 Plan Apo oil lens, an Andor confocal unit, an sCMOS Sona 4.2B-11 camera (Andor) and a laser bench with 405-561 nm diode (Andor). Exposure time was 0.3 s for the green channel and 0.2 s for the red channel. Microscopy images shown in Figure S5A are maximum intensity projections obtained from z-stacks of 7 slices at 0.3 µm interval, acquired using the same microscope setup from Nikon. Exposure time was 0.35 s. For the sake of comparison, images in Figure 3A and S5A are displayed using the same intensity range with Metamorph 7.7.

Quantification of the time for acto-myosin ring assembly, maturation and constriction was performed by analyzing the time between the initial recruitment of myosin cortical nodes and their compaction into a tight ring, the time until the ring starts to constrict and the time until the myosin signal disappears after final constriction, respectively. Scatter dot plots of the measured times were prepared with GraphPad Prism. The number of cells used for each strain, the mean measured times and the statistical test used to evaluate differences, the latter performed with GraphPad Prism, are indicated in the respective legend. Actin patch and actin cable number per cell were quantified from maximum intensity projections obtained from z-stacks of 7 slices at 1 µm from cells in G2 phase (around 10 µm long). Scatter dot plots of the measured actin cables and patches were prepared with GraphPad Prism. The number of cells used for each strain and the mean numbers of cables and patches are indicated in the respective legend.

##### Polarimetry measurements in the cytokinetic ring of live fission yeast

For live polarimetry measurements, strains co-expressing Af1, Af7, L1 or L22 and an mCherry-tubulin marker (strains SR3.51, SR3.54, SR3.57 and SR3.58 in Table S3) were incubated at 25°C in YE5S medium. 1 mL of exponentially growing cells were harvested by centrifugation for 60 s at 800 g, most of the supernatant was discarded and 1 mL of the cells was deposited onto a 2% YE5S agar pad at the center of a polydimethylsiloxane slide chamber prepared as described in ^104^.

Three large field-of-view images (66 x 66 μm) typically containing 5-10 dividing cells per image, were collected for each strain. Before each polarimetry measurement, a two-color z stack was acquired (Δz = 1.0 μm) to image both GFP fusions and microtubules; the distribution of the latter was used in addition to the morphology of the actomyosin ring to confirm that cells were undergoing cytokinesis. A polarimetry stack using 18 polarization angles was then recorded for each position within a z stack (Δz = 1.0 μm) for the GFP channel, and thus allowed to obtain polarimetry images throughout the cytokinetic rings, containing both tangential-most views with the ring parallel to the xy plane, and more equatorial views showing cross-sections that appear as spots on either side of the ring. An exposure time of 0.5 s was used per polarized image. To minimize bias in the measured orientations due to the contribution of off-plane orientations we focused on the tangential-most views for the analysis (Figure 3F-H). One tangential view of the ring was analyzed per cell; in a few cells where both tangential views were present in the respective z planes, both were analyzed. Equatorial views were used for measuring the diameter of the constricting rings (Figure S5E).

Polarimetry stack images were first processed with the open-source image processing software ImageJ/Fiji. Images within each polarimetry stack were registered using the StackReg plugin to correct for drift during the acquisition. The z planes containing the tangential-most views of the ring were identified for each cell. The pixels of the cytokinetic ring for analysis were selected by a combination of intensity thresholding and manual selection of the region to analyze. Each region of interest contained typically 40-80 analyzed pixels i.e. 40–80 color-coded sticks in the tangential-most view of the cytokinetic ring per cell (Figure 3F-G). Analysis and data representation, including color-coded stick representations of the measured angles per pixel and polar histograms were done with PyPOLAR. The source code and desktop app are available at https://github.com/cchandre/Polarimetry.git. GraphPad Prism was used to generate scatter plots of the quantified ψ angle distributions per strain; the number of cells measured for each strain, the respective median values and the statistical test used to evaluate differences are mentioned in the respective legend. Considering that Af7 and L22 dipoles are parallel to actin filaments, in order to assess the extent to which the measured actin filament orientations were more parallel or more perpendicular with respect to the ring axis, the ring axis angle in each cell was used as the reference angle in the “reference angle” tool in PyPOLAR to normalize the angle distributions from 0°– 180° to 0°–90° and generate 0°–90° polar histograms, with 0° and 90° defining orientations parallel and perpendicular to the ring axis, respectively (Figure S5F).

#### Characterization of *C. elegans* strains expressing actin organization reporters

##### Plasmid construction for generation of transgenic animals

Sequences encoding actin organization reporters were codon optimised for optimal expression in the worms using the *C. elegans* codon adapter web tool ^105^ and synthetized by GENEWIZ. Plasmids pFBR101, pFBR102 and pFBR105 (see Table S1) were constructed in two steps from pML36 (kind gift from Michel Labouesse lab), which contained a pCFJ151 backbone (ttTi5605 insertion homology arms) with a dpy-7 promoter for epidermal cell expression and a unc-54 3’UTR (universal 3’UTR for optimal expression). Briefly, the plasmid pML36 was opened and amplified by PCR using custom made oligos (Sigma-Aldrich). The sequences encoding organization reporters were also amplified by PCR and joined using overlapping ends into opened pML36 plasmid using the NEBuilder HiFi DNA Assembly Cloning kit (New England Biolabs, Cat#E5520S). All PCR reactions were carried out by Phusion High-Fidelity DNA Polymerase (ThermoFisher Scientific, Cat#F531L). Primers were custom made by Sigma-Aldrich. The sequences of all oligos are listed in Table S2. The final plasmids were verified by DNA sequencing (Eurofins Genomics, Germany).

##### Transgenic worm construction by MosSCI method

Worm MosSCI transgenesis was performed by direct microinjection as described in ^106^. Briefly, the injection mix was injected in the arm of both gonads in the young hermaphrodite animal of EG6699 strain. The injection mix contained a cocktail of pJL43.1 (50 ng/mL), pCJF90 (2.5 ng/mL), pCFJ104 (5 ng/mL), and an expression clone (50 ng/mL) in DNase/RNase-free water. All plasmids used for injection were purified by HiSpeed Plasmid Midi kit (Qiagen, Cat#12643). After injection, transgene insertion screening was performed as described at http://www.wormbuilder.org. Transgenic animals were verified by PCR genotyping and DNA sequencing.

##### Worm embryonic growth and lethality tests

A few young hermaphrodite animals were picked and fed on freshly seeded Escherichia coli OP50 for 2h. After 2 h, all animals were transferred on fresh OP50 plates and laid eggs were counted. This process was redone until sufficient number (>1000) of embryos were achieved and counted. After 12-16 h, all previously scored embryo-containing plates were recounted for unhatched embryos (dead eggs) and hatched larvae. For this experiment, strains were grown at 20°C. The embryonic lethality for each strain, scored as the percentage of unhatched embryos, is shown in Figure 4C.

Embryonic growth rate was measured by imaging embryos by differential interference contrast (DIC) microscopy on a Leica DM6000 microscope until they hatch as larvae. Briefly, embryos were collected by dissecting gravid hermaphrodites in M9 medium and mounted on a 5% agarose pad for imaging. Z-stack images were acquired with 40-50 planes per embryo and a Δz interval of 1 μm, and with a 10 min interval for 12-14 h at 20°C. Embryonic length was measured manually with the segmented line tool in the ImageJ/Fiji software and growth curves plotted with Microsoft Excel software (Figure 4D). The number of embryos measured for each strain is indicated in the respective legend.

##### Polarimetry measurements in live C. elegans embryos

More than >30 young gravid hermaphrodite animals were picked and fed on freshly seeded OP50 E. coli overnight. Next day, mixed stage embryos were picked and mounted on a 5% agarose pad in M9 medium for imaging. Temporary hypoxic conditions were created by adding OP50 E. coli, preventing embryonic muscle activity that usually starts around 1.7-fold. Embryonic stages were evaluated by brightfield microscopy. Length measurements were subsequently performed using ImageJ/Fiji. Polarimetry stacks using 18 polarization angles were recorded in the epidermis of 1.5-fold, 1.5-2-fold and >2-fold stage embryos. The pixels containing dorsal and ventral epidermal cells (DV cells) and seam cells were selected by a combination of intensity thresholding and manual selection of the region to analyze. Analysis and data representation, including color-coded stick representations of the measured angles per pixel and histograms were done with PyPOLAR. GraphPad Prism was used to generate scatter plots of the quantified ψ angle distributions per strain and per developmental stage; the number of embryos for each strain and for each stage and the respective median ψ values are mentioned in the respective legend. To assess how the measured actin filament orientations in DV and seam cells distribute with respect to the DV/seam boundary for each developmental stage, the ρ angle distributions were normalized with respect to the DV/seam boundary from 0°–180° to 0°–90° to generate 0°–90° polar histograms. The DV/seam boundary for each embryo was drawn manually with the freehand line selection tool or the elliptical selection tool in Fiji and converted to a mask which was then used with the “edge detection” tool in PyPOLAR to define the boundary as the reference for the normalization of the angles. Considering that Af7 and L22 dipoles are parallel to actin filaments and that L45 dipoles are perpendicular to actin filaments, the more perpendicular mean actin filament orientations are to the boundary, the closer the angle values are to 90° (for Af7 and L22) or to 0° (for L45) (Figure 4H,J,L and Figure S5J,K).

#### Generation of *Drosophila* expressing selected actin organization reporters

##### *Drosophila* transgenics

Selected actin organization reporters were subcloned into pUASp plasmids for generating *Drosophila* transgenics. The respective mammalian expression plasmids were used as templates to subclone the reporters into *Kpn*I/*BamH*I linearized UASp vectors using seamless cloning (In-Fusion HD Cloning Plus Kit, Takara Bio, Cat#638910). All primers were Cloning Oligo synthesis and purification quality from Eurofins Genomics and are listed in Table S2. Restriction enzymes were FastDigest enzymes from Thermo Fisher Scientific. All plasmids were verified by sequencing (Eurofins Genomics) after each cloning step. Midipreps of each UASp construct DNA were sent to BestGene Inc. (California, USA) for injections into *D. melanogaster* w1118 embryos and generation of the transformants UASp–L22, UASp–L45, UASp–U20, and UASp–Af7 (see Key Resources Table and Table S1).

##### Preparation of live *Drosophila* embryos for polarimetry measurements

mat–α-tub–GAL4 females were crossed to UASp–Af7 males, and F2 embryos were collected and prepared for imaging following standard procedures ^107^. Briefly, F1 progeny was placed in embryo collection cages with fresh yeasted apple juice agar plates. For live imaging, cellularizing F2 embryos were dechorionated with 50% bleach, washed with water, transfered onto a heptane glue-coated round coverslip, covered with halocarbon oil 200 (Tebubio, Cat#25073) and mounted in an Attofluor cell chamber (ThermoFisher Scientific, Cat#A7816).

##### Polarimetry measurements in live cellularizing *Drosophila* embryos

Polarimetry stacks using 18 polarization angles were recorded in focal planes through actomyosin rings at the invaginating membrane front (Figure 5A) or in focal planes apicolaterally to the former through the basal adherens junctions (Figure 5B). An exposure time of 0.2 s was used per polarized image. The pixels of the actomyosin rings or basal adherens junctions for analysis were selected by a combination of intensity thresholding and manual selection of the region to analyze. Analysis and data representation, including color-coded stick representations of the measured angles per pixel and polar histograms were done with PyPOLAR. Considering that Af7 dipoles are parallel to actin filaments, in order to assess how the measured actin filament orientations distribute with respect to the actomyosin ring contour, the “edge detection” tool in PyPOLAR was used in combination with intensity thresholding to isolate the ring contour-associated pixels and normalize the angle distributions from 0°–180° to 0°–90° and generate 0°–90° polar histograms, with 0° and 90° defining orientations parallel and perpendicular to the ring contour, respectively (Figure S6A).

#### Characterization of actin organization reporters in the *Drosophila* flight muscle

##### Flight tests

Flight tests were performed as described in ^108^. 3 to 7 day-old males were dropped into a 1 m long / 15 cm in diameter plexiglass cylinder with marked sections. Landing in the different sections depends on their flight ability, which was thereby scored (top 40-cm section: wild type, middle 40-cm section: impaired flight (”weak flier” in Figure 5C), bottom 20-cm section: flightless) (see cartoon in Figure 5C). For each genotype, flight assays were performed three times with a minimum of ten males per assay. The total number of flies scored for each genotype is mentioned in the respective figure legend. GraphPad Prism was used to generate bar graphs of the quantified flight ability per genotype and the mean percentages are mentioned in the respective legend.

##### Preparation of fixed adult flight muscles

Head, wings and abdomen were cut off the thorax of anaesthetized adult flies with fine scissors, and the thoraxes were fixed for 30 min in 4% paraformaldehyde in PBST (PBS + 0.3% Triton X-100). After three 10 min washes in PBST, thoraxes were placed on double-sided tape and cut sagittally dorsal to ventral with a microtome blade (Feather C35). The thorax halves were placed in PBST with Alexa 568-phalloidin (Invitrogen, 1:500) and incubated overnight at 4°C on a shaker. Hemithoraxes were then washed 3 times 10 min in PBST at room temperature and mounted in Vectashield with 2 spacer coverslips on each side.

##### Preparation of live flight muscles

Flight muscles were dissected, mounted in Schneider medium (no fixation), and imaged within 30 min following dissection. After removal of the head, abdomen and wings, a first incision was performed through the cuticle with sharp forceps (Dumont #5 forceps, Fine Science Tools, Cat#11252-20) at the median plane. The thorax was then gently pulled open into two halves, which were then fully disconnected through cutting of the ventral connective tissues using fine dissection scissors (Fine Science Tools, Cat#15009-08). The dissection resulted in relatively intact flight muscles still attached to the tendon cells of the thorax. Samples were mounted in Schneider medium using two coverslip spacers and imaged immediately.

##### Polarimetry measurements in live Drosophila flight muscle

The polarimetry analysis shown was from flight muscle expressing the reporters throughout muscle development with the Mef2-GAL4 driver apart from flight muscle expressing Af7, for which Af7 was expressed only transiently after muscle development with the tub-GAL80ts ; Mef2-GAL4 driver. One hemithorax per animal was used for polarimetry measurements, with 4-7 hemithoraces measured for each strain. Polarimetry stacks using 18 polarization angles were recorded in 1-7 different fields of view for each hemithorax. Ten myofibrils were analyzed per field of view (red-outlined boxes in Figure 5E-G). The pixels containing individual myofibrils within each field of view were selected by a combination of intensity thresholding and manual selection of the region to analyze. Each myofibril contained typically 3,000-10,000 analyzed pixels. Analysis and data representation, including color-coded stick representations of the measured angles per pixel and histograms were done with PyPOLAR. The histograms shown in Figure 5E-G are from single myofibrils. GraphPad Prism was used to generate scatter plots of the quantified ψ angle distributions per strain; the number of myofibrils measured for each strain, the respective median values and the statistical test used to evaluate differences are mentioned in the respective legend.

#### Characterization of actin organization reporters in the *Drosophila* wing

For wing analysis, we anesthetized 50 young adult males from the progeny (using CO_2_) and removed one wing from each fly with fine tweezers. These wings were then directly mounted between a slide and a coverslip using UV-cured optical adhesive (Thorlabs, Cat#NOA63). Images of the wings were captured using a digital microscope (Dino-Lite). For wing size analysis, we utilized landmarks within the wing vein pattern to measure specific distances. For the long axis, we measured the distance between the proximal end of L5 and the distal end of L3, following the nomenclature from ^109^. For the short axis, we measured the distance between the distal end of L5 and the intersection of the opposite side of the wing with a line perpendicular to the long axis, passing through the distal end of L5. These two distances are represented in Figure S6B. MATLAB was used to generate box plots of the quantified wing area. The number of wings for each genotype, the respective median values of L_L_·L_S_ and the statistical test used to evaluate differences are mentioned in the respective legend.

#### *in vitro* measurements of binding affinities to actin filaments for selected actin organization reporters

##### Generation of bacterial expression plasmids

Selected actin organization reporters were subcloned into pnCS (pCDF-DuET backbone) ^110^ plasmids for bacterial expression (see Table S1). The respective mammalian expression plasmids were used as templates to subclone TEV-cleavable Strep-tag-II-tagged reporters into *Nde*I/*BamH*I linearized pnCS vectors using seamless cloning (In-Fusion HD Cloning Plus Kit, Takara Bio, Cat#638910). All primers were Cloning Oligo synthesis and purification quality from Eurofins Genomics and are listed in Table S2. Restriction enzymes were FastDigest enzymes from Thermo Fisher Scientific. All plasmids were verified by sequencing (Eurofins Genomics) after each cloning step.

##### Purification of recombinant actin organization reporters

Recombinant Af7, L22, L45, and U20 were expressed and purified from *Escherichia coli* BL21(DE3). For each construct, 1 L of LB media containing 80 μg/mL spectinomycin was grown, shaking at 200 rpm at 37°C until the absorbance at 600 nm was between 0.6 and 0.8. Expression was induced by the addition of 1 mM of IPTG and by growth overnight at 16°C. Cells were harvested by centrifugation (2,000 g for 10 min, at 4°C), washed once in ice cold 30 mL of PBS, and a second centrifugation run. Cells were resuspended into buffer A (50 mM Tris-HCl pH 8, 300 mM KCl, 5 mM MgCl_2_, 1 mM DTT) supplemented with a cOmplete^TM^ protease inhibitor cocktail (Roche), 0.1 mM PMSF, 10 mg/mL DNAse, and 0.25 mg/mL lysozyme. After 30 minutes incubation, cells were sonicated for a total duration of 5 minutes with 10 s on, and 20 s of rest. Lysate was pelleted at 20,000 g for 30 min at 4°C. Supernatant lysate was incubated for 2 hours at 4°C with 2.5 mL Strep-Tactin Sepharose High Performance resin (Cytiva, Cat#28-9355-99) previously washed with buffer A. Elution was performed using an Econo-column and a solution of buffer A with 2.5 mM desthiobiotin. The sample was centrifuged for 10 minutes at 7,000 g, concentrated using Amicon filter (cutoff 30 kDa) down to ∼ 2 mL and injected onto a HiLoad 16/60 Superdex 200 gel filtration column on an Akta pure system, equilibrated with the final storage buffer (20 mM Tris-HCl pH 8, 50 mM KCl, 1 mM MgCl_2_, 1 mM DTT). Relevant fractions were pooled, protein concentration measured using an extinction coefficient at 280 nm of 52,495 (Af7), 26,025 (L22), 24,535 (L45), and 60,515 (U20) M^−1^.cm^−1^, and flash frozen using liquid nitrogen for −70°C long term storage.

##### alpha-skeletal actin protein purification and fluorescence labeling

α-skeletal muscle actin was purified from rabbit muscle acetone powder following the protocol described in ^111^, based on the original protocol from ^112^. Actin was fluorescently labeled on accessible surface lysines of filamentous actin using Alexa Fluor 568 succinimidyl ester (Thermo Fisher Scientific, Cat#A20003), and used at 10% labeled fraction.

##### Co-sedimentation assay

The binding affinity of L22, L45 and U20 to actin filaments was measured by performing co-sedimentation assays. First, 20 µM actin was polymerized in FME buffer for 2 hours at room temperature. Polymerized actin was then diluted to 2 µM and incubated with a range of reporter concentrations for 5 minutes at room temperature. Solutions were centrifuged at 200,000xg for 30 minutes, at room temperature. Pellets from 3 independent experiments were analyzed by SDS–PAGE, with Coomassie blue stained band intensities from reporters measured using FiJi/ImageJ and normalized first to the actin intensity band of each well, then to the most intense reporter band of the gel. Binding affinities were determined by fitting data points with a quadratic equation Fraction_bound_ = [P]_free_⁄([P]_free_ + K_D_) using the least-square curve_fit function from the Scipy python package, giving the best value and its 95% confidence interval. We note that the concentration of purified recombinant U20 was not high enough to allow us to reach a plateau (Figure S6G).

##### Fluorescence microscopy assay

The binding affinity of Af7 to actin filaments was performed using TIRF microscopy. Assays were performed in between two PEG-silane passivated coverslips, using melted parafilm as a spacer. Surfaces were previously passivated with PEG-Silane (Laysan Bio, Cat#MPEG-SIL-5000-5): out-of-the-box coverslips (Menzel-Gläser 22×40 1.5#) were first exposed to plasma for 5 minutes. 200 µL PEG-Silane (1 mg/mL in 80/20% ethanol/water, pH 2.0) solution was deposited and dried at 70°C for 20 minutes. Coverslips were rinsed with ethanol and deionized water, and finally dried with filtered air. 0.6 µM of 10%-labeled actin was mixed with a range of concentrations of Af7 in FME buffer, supplemented with 0.2% methylcellulose and imaged within 5 minutes, with a Nikon TiE inverted microscope, equipped with a 100x 1.49 NA oil immersion objective and a Kinetix22 sCMOS camera (Teledyne Photometrics). Experiments were performed at 25°C (objective-collar heater from Oko-lab). Image acquisition and TIRF illumination (iLAS2 from Gataca Systems) were controlled using Micromanager software.

For each condition, Af7 fluorescence intensity along actin filaments was measured using FiJi/ImageJ, subtracting local background fluorescence. Binding affinity was determined by fitting data points with a quadratic equation Fraction_bound_ = [P]_free_⁄([P]_free_ + K_D_) using the least-square curve_fit function from the Scipy python package, giving the best value and its 95% confidence interval.

##### Buffers

Co-sedimentation and fluorescence microscopy experiments to determine the affinity of actin organization reporters to actin filaments were performed in FME buffer: 5 mM Tris HCl pH 7.4, 50 mM KCl, 1 mM MgCl_2_, 0.2 mM EGTA, 0.2 mM ATP, 10 mM DTT, and 1 mM DABCO. FME buffer was supplemented with 0.2% methylcellulose (4000 cP at 2%; Merck, Cat#M0512) for TIRF microscopy assays to keep filaments in the vicinity of the glass bottom and image them using TIRF laser penetration depth set to 70 nm.

#### Protein structures and protein sequence alignments

Cartoon representations of protein structures were generated with the open-source software PyMOL (Schrödinger). The structure shown for cpGFP1-10/11 in Figure 2A corresponds to the structure of circular permutated red fluorescent protein Kate (PDB 3RWT) and is used to illustrate the design principle of our circular permutants. The PDB IDs for the remaining structures are as follows: 2B3P (sfGFP, Figures 2A, 6A, S1A), 1GFL (wild-type GFP, Figure S1B), 7AD9 (Lifeact-F-actin complex, Figure S1F,G), 1QAG (Utrophin, Figure S3G), 4N6T (Adhiron/Affimer, Figure S4I) and 5JLF (F-actin-tropomyosin complex, Figure 6B). The structure of F-tractin (Figure S4B) was generated using the AlphaFold database at EMBL-EBI. The multiple sequence alignment of Lifeact sequences (Figure S1E) was generated with Clustal Omega (EMBL-EBI). Graphical representations illustrating the conservation of residues for Lifeact, F-tractin and G-actin (Figures 6C, S1E, S4A) were generated using the WebLogo application (University of California, Berkeley). Interface areas were analyzed using PISA calculations as implemented on the EMBL-EBI server and visually inspected using PyMOL.

#### Polarimetry

##### Optical setup

Fluorescence images were acquired with a confocal spinning disk unit (CSU-X1-M1, Yokogawa) connected to the side-port of an inverted microscope (Eclipse Ti2-E, Nikon) using a x2 magnifier (Yokogawa), a Nikon Plan Apo ×100/1.45 NA oil immersion objective lens and an EMCCD camera with 1024×1024 pixels, 13×13 µm pixel size (iXon Ultra 888, Andor) resulting in an image pixel size of 65 nm. Z-stacks were acquired using a piezo stage (P-736, PI). The lateral position of the sample was controlled with a translation piezo stage (U-780, PI). The spinning disk is equipped with a multiline dichroic mirror (Di01-T405/488/568/647-13×15x0.5, Semrock) and an emission filter wheel with filters adapted to the studied emission: band pass 540/80 for EGFP/sfGFP and AF488 (FF01-540/80-25, Semrock), band pass 593/46 for sfCherry2 (FF01-593/46-25, Semrock), and long pass 655 for SiR-actin (Et655lp, Chroma). The laser excitation is provided by polarized continuous lasers (488-, 561- and 641-nm laser lines, Sapphire, Coherent) combined with a set of dichroic mirrors, each of the laser being used separately with a power of typically 0.5 mW at the entrance of the spinning disk. The laser beams are sent into an electro-optic modulator (EOM) (Pockels cell, No 28-NP; Quantum Technology) followed by a quarter wave plate (WPQ05M-488; Thorlabs) to create a linear rotating polarization. The voltages sent to the Pockels cell to provide known polarization rotations are determined in a preliminary calibration step, using a polarimeter based on the quarter wave plate method, as described in ^67^. As the whole optical path involves reflections on mirrors and transmission through a dichroic mirror, the polarization after the Pockels cell system is likely to be deformed. Polarization distortion compensation of the spinning disk dichroic mirror is provided by placing an identical dichroic mirror (Di01-T405/488/568/647-13×15x0.5, Semrock) in the path of the laser line just after the quarter wave plate, such that s and p polarization components are exchanged at the first and second dichroic transmissions. This configuration ensures minimization of the polarization ellipticity and diattenuation produced by the dichroic mirror. The remaining distorsions are characterized following the procedure of ^67^, using a polarimeter based on the quarter wave plate method. The beam is then expanded using a 10× telescope (BE10, Thorlabs) and sent directly to the microlens array of the CSU by reflection on a second entrance mirror. The microlens and pinhole arrays of the CSU disks rotate synchronously at a speed of 1,800 rpm, while the EMCCD and EOM are synchronized to ensure a fast stack recording for a given number of incident polarization ^67^. Exposure times are in the range of 0.1-0.5 s, and 18 polarization angles are typically measured per polarimetry stack, which leads to a few seconds per polarimetry stack.

##### Signal processing

Fluorescence is generated from the coupling of fluorophore dipoles with the incident linearly polarized electric field denoted *E*(*α*), whose orientation is an angle *α* with the horizontal axis *X* of the sample plane. Inside the confocal volume, each fluorescent molecule exhibits an absorption dipole vector *μ*_*abs*_ with an orientation (*θ*, *ϕ*) in the macroscopic sample frame. The recorded fluorescence intensity from a single molecule is proportional to the absorption probability *θ*_*abs*_(*θ*, *ϕ*) = |*μ*_*abs*_(*θ*, *ϕ*) · *E*(*α*)|^2^. The total intensity from an ensemble of molecules in the focal volume is therefore the sum of the intensities from all single molecules present in this volume, whose size is typically 300 nm laterally and 600 nm longitudinally. This results in an averaged intensity: *I*(*α*) = ∬|*μ*_*abs*_(*θ*, *ϕ*) · *E*(*α*)|^2^ sin *θ dθdϕ* ^34^. The intensity is thus maximized when the absorption dipoles of the molecules are aligned with the electric field. We assume that the orientations explored by molecular dipoles are constrained within an angular cone of total aperture angle *ψ*, oriented in the sample plane along the direction *ρ* relative to *X*, the horizontal axis of the sample plane. Physically, *ψ* is related to a « molecular order » quantity, which determines the degree of angular variations present within the focal spot at a given pixel position, averaged over time and space. Note that when fluorescent molecules are attached to actin with a degree of angular fluctuations due to their linker to actin, *ψ* encompasses three contributions : (1) the mean tilt angle ξ of the molecule with respect to the actin filament axis, (2) the angular fluctuations of the molecule due to its linker flexibility, and (3) the static organization of the actin filaments. The mean orientation *ρ*, on the other hand, determines the mean direction of the molecules. Therefore when the molecules are attached to actin in a constrained manner (i.e. angular fluctuations are not isotropic), in an assembly of aligned filaments, *ρ* can take two values : either *ρ* = 0° when the tilt angle of the molecules ξ is close to the filament axis with ξ < 45°, or *ρ* = 90° when the molecules are away from the filament axis with ξ > 45°. Thus, the angles *ρ* and *ψ* quantify the full information on the molecular organization at each pixel of an image. We note that the measurements performed in this work are limited to a projection of the fluorophores’ distribution in the sample plane, which is imposed by the manipulation of light polarization in this plane. This 2D projection leads to an overestimation of the order angle ψ when the cone distribution is tilted more than 45° out of plane ^34^.

The angles *ρ* and *ψ* are deduced from the measurement of the intensity modulation *I*(*α*), which takes the form ^34,67^: *I*(*α*) = *a*_0_ + *a*_2_(*ρ*, *ψ*) cos 2*α* + *a*_2_(*p*, *ψ*) sin 2*α*. The coefficients *a*_2_(*p*, *ψ*) and *a*_2_(*p*, *ψ*), which can be directly related to the parameters (*ρ*, *ψ*) (see below), are deduced from the decomposition *I*(*α*) into circular functions (cos 2*α*, sin 2*α*). In practice, when several input polarization angles *α*_*k*_ are used in a polarimetry stack (typically, for 18 polarization angles, *α*_*k*_ = 0, 10°, …, 170°), we use the operations *a*_2_ = 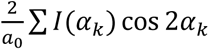 and 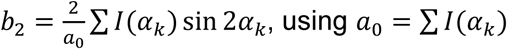.

To retrieve the angular parameters (*ρ*, *ψ*) from the measured quantities *a*_2_(*ρ*, *ψ*) and *a*_2_(*p*, *ψ*), the following method is used to account for polarization distorsions ^34^: The presence of polarization distorsions is modelled in the intensity equation *I*(*α*) = ∬|*μ*_*abs*_(*θ*, *ϕ*) · *E*(*α*)|^2^ sin *θ dθdϕ* by including a distorted *E*(*α*) = (cos *α*, *γ* sin *α e*^*iδ*^), with *γ* a diattenuation factor, which produces an energy loss between the s and p polarization components of an electric field, and *δ* a birefringence factor, which produces a phase difference between the s and p components. In this model, the polarization distorsions are supposed to originate from an equivalent phase plate whose axes coincide with the horizontal and vertical directions of the sample, which is reasonable considering that all reflections in the optical path involve s and p directions along these axes. Including these distorsions allows the construction of a map of the dependence of both *a*_2_and *b*_2_parameters, as fuctions of (*ρ*, *ψ*). Without any distorsions, these maps take the form of disks from which (*ρ*, *ψ*) can be unambiguously determined by the one-to-one relationship between (*ρ*, *ψ*) and (*a*_2_, *b*_2_) ^34^. In the presence of distorsions, the disks are deformed but the relation stays unambiguous, therefore it is possible to find (*ρ*, *ψ*) from the measurement of (*a*_2_, *b*_2_), using a minimization method in the (*ρ*, *ψ*) *vs* (*a*_2_, *b*_2_) lookup table for instance. Finally, the parameters (*ρ*, *ψ*) extracted from the (*ρ*, *ψ*) *vs* (*a*_2_, *b*_2_) disk analysis are represented in a single polarimetry image that combines molecular order and orientation, superimposed to the fluorescence intensity image built from the total intensity ∑ *I*(*α*_*k*_).

In experimental measurements, the *I*(*α*) modulation is affected by noise, which impacts the determination of the (*ρ*, *ψ*) parameters. The precision on the determination of (*ρ*, *ψ*) increases as the inverse square of the total intensity. It has been shown that above 5000 photons per pixel (which is typically the case for GFP imaging), the precision reaches about 1° for *ρ* and 3° for *ψ*, except at extreme high-order conditions (*ψ*∼0°) where the precision in *ψ* reaches 5° ^34^.

We note that the reasoning for the dependence of absorption probability on the fluorophore dipole orientation is similar to that for the dependence of emission probability: the polarized emission scheme exploited in fluorescence anisotropy and polarization emission analysis ^18,113^ is not exploited in this study, but could be similarly applied ^24,33,66,78–80^. Finally, the angle ψ used in this work can also be directly related to other quantities used to define molecular orientational organization, in particular the generic, distribution-independent order parameter used in aligned structures such as lipid membranes and liquid-crystalline polymers ^114^.

#### QUANTIFICATION AND STATISTICAL ANALYSIS

The quantification method for each experiment is described in the respective method details section. The statistical details of the experiments, including the exact value of n and what n represents, the definition of center, dispersion and precision measures (mean, median, SD, SEM) in the plots and graphs, the software and statistical test used to evaluate statistical significance of differences, and the definition of statistical significance are mentioned in the method details sections and respective Figure legends.

## KEY RESOURCES TABLE

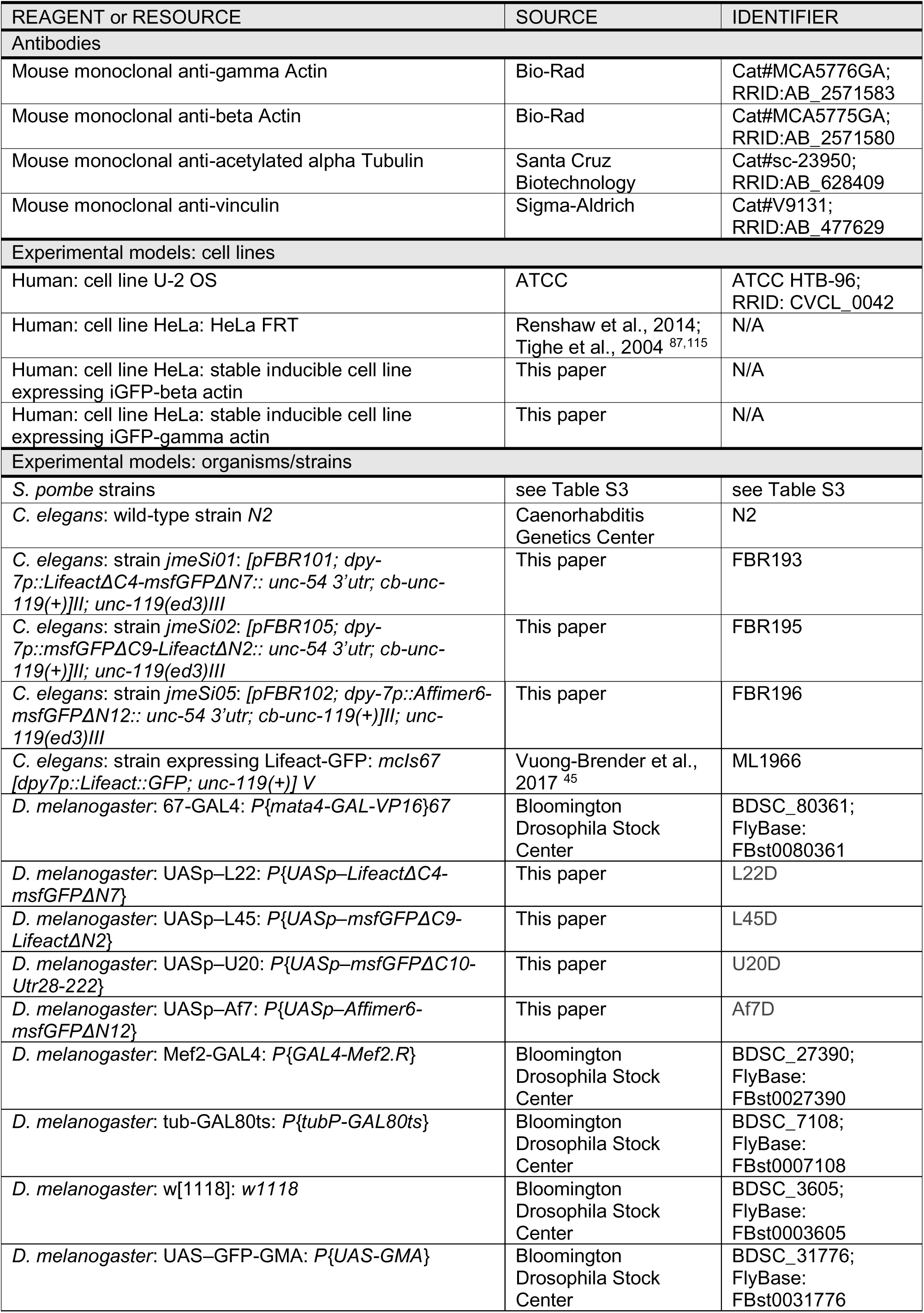
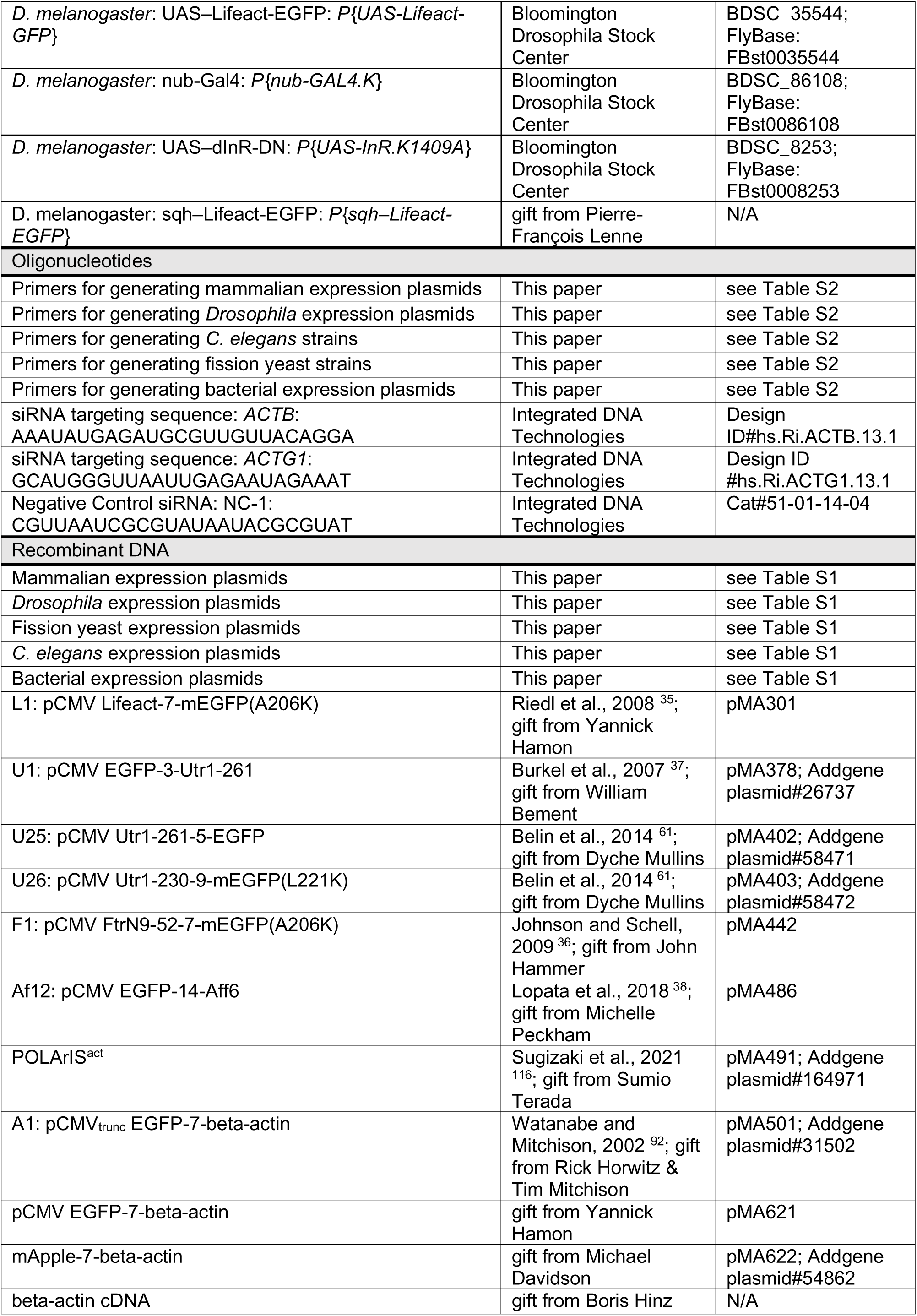
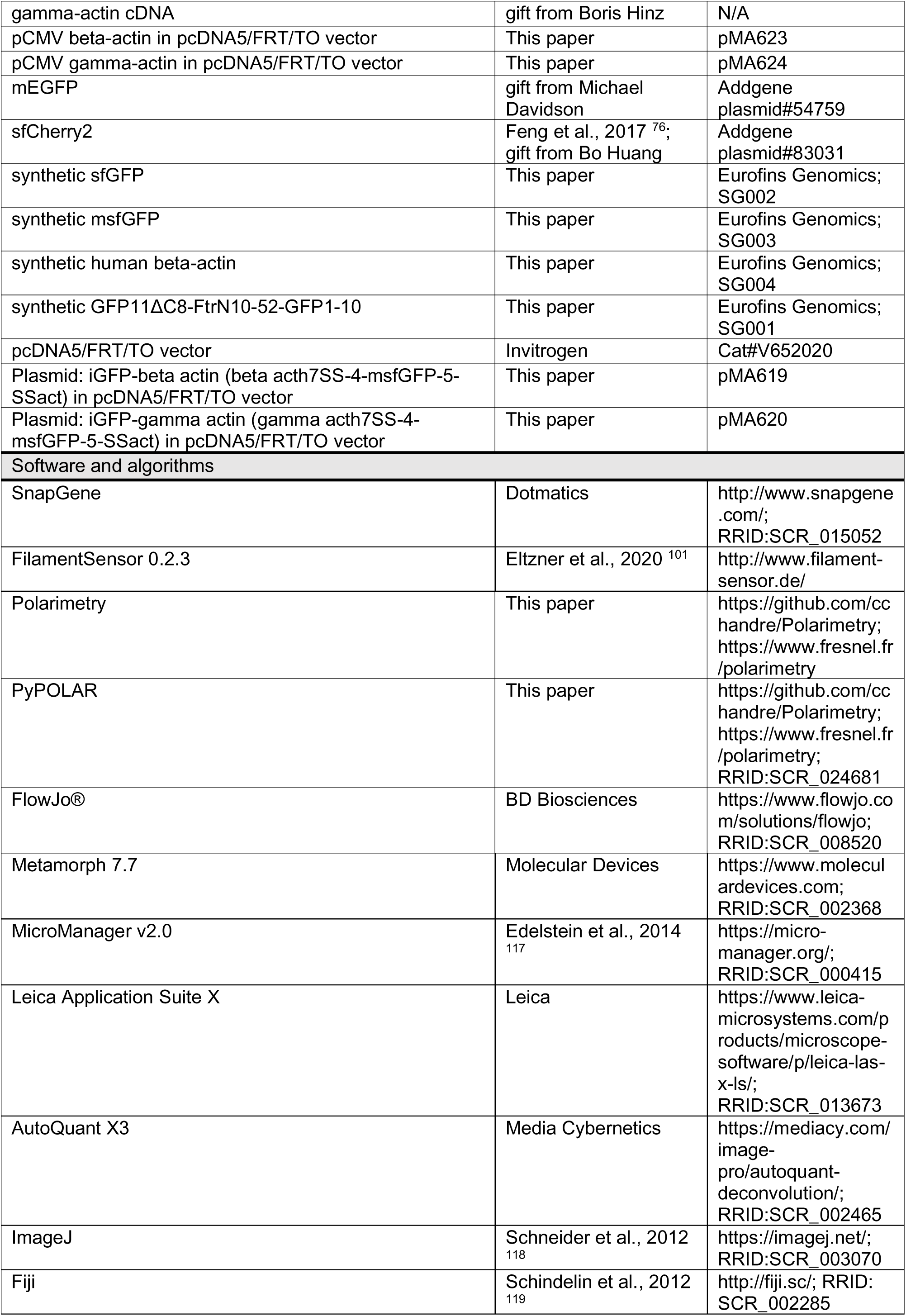
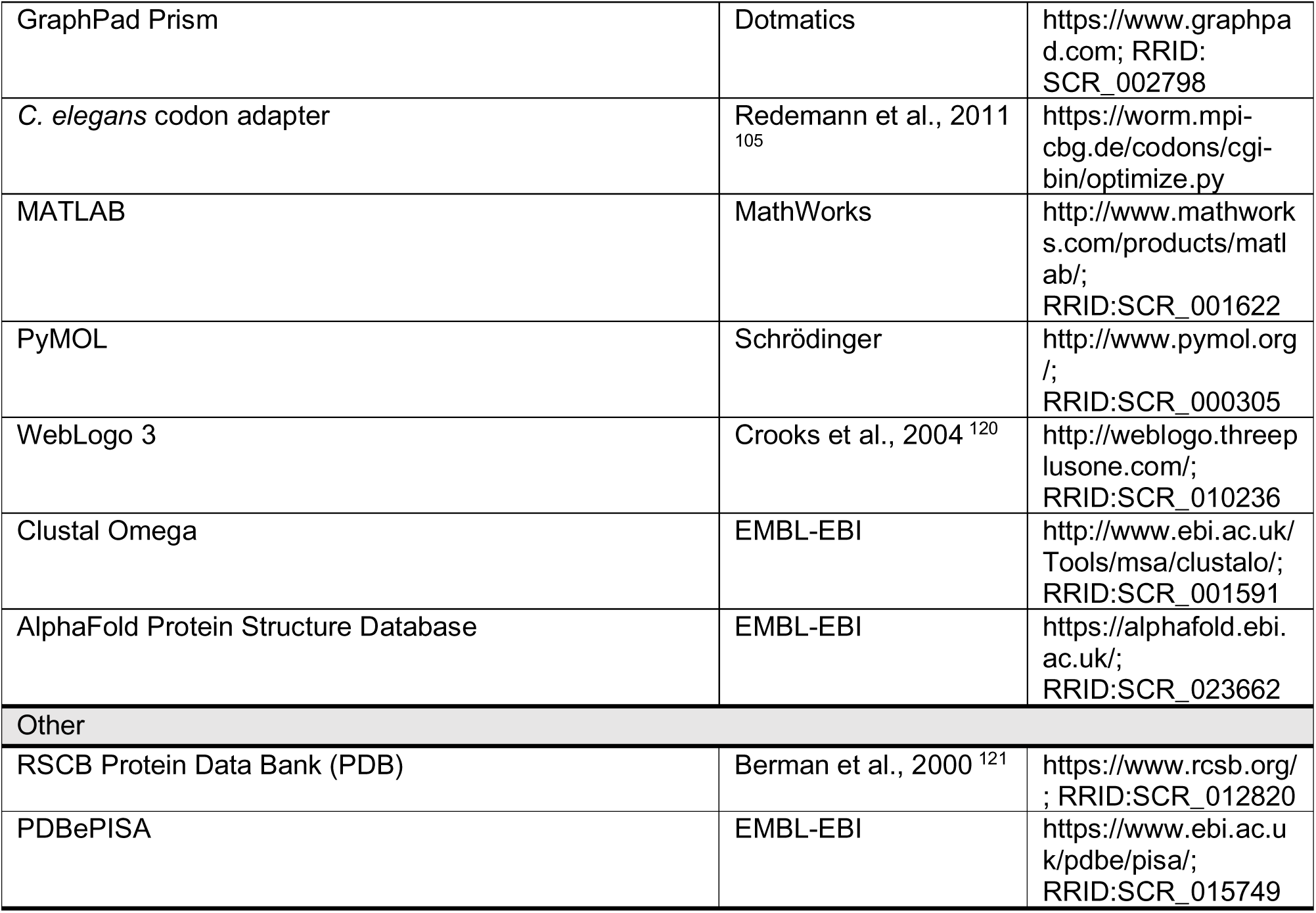

### Supplemental data

#### Design of GFP-based reporters with constrained GFP mobility

Tailoring the available F-actin localization reporters for organization measurements requires that GFP mobility is constrained. We reasoned that there are three main sources that contribute to the flexibility of the EGFP. The first source is the presence of an amino acid linker between the actin-binding moiety and the fused GFP. All the widely used GFP fusions tested (Figure 1E) have been generated by standard restriction-ligation cloning using the multiple cloning sites of the GFP-C1 and GFP-N1 vectors from Clontech ^1^ and thus inevitably introducing several amino acid residues in between the GFP terminus and the ABD. Although the presence of such a linker is reasonably assumed to be important for minimizing interference with protein folding and interactions with actin-binding proteins, we hypothesized that it also contributes to the rotational mobility of the fused EGFP. The second source is the flexibility of the terminus of the ABD to which GFP is fused. Thus, shortening or removing altogether the linker and/or flexible stretches from the terminus of the ABD could be promising approaches for constraining the GFP, assuming that protein folding, F-actin binding and interactions with actin-binding partners are not compromised.

The third source of flexibility are the N- and C-termini of EGFP themselves. The crystal structures of GFP ^2,3^, EGFP ^4^ and practically all fluorescent protein (FP) variants with the same termini show that the C-terminus after the end of the β11 strand comprises 11 residues that are unstructured and most of which are absent from the respective crystal structures due to their flexibility (Figure S1A,C). The N-terminal stretch preceding the β1 strand comprises 12 residues, with residues 5-9 forming a 3_10_ helix (in red in Figure S1A,C), while the first 4 amino acid residues are similarly unstructured and often not visible in the crystal structures. Shortening of the N-terminus by 4-6 residues has indeed been shown to constrain the mobility of GFPS65T and EGFP in C-terminal fusions to septins and nucleoporins ^5–7^. However, more extensive truncation of the termini has not been explored to date. This is probably due to early studies with wild-type GFP, GFPS65T or EGFP that showed that removing more than six N-terminal or more than nine C-terminal residues significantly impairs or abolishes GFP fluorescence altogether ^8–10^.

Prompted by the recent finding that better-folded variants of GFP tolerate more extensive terminal truncations ^11^, we made a side-by-side comparison of truncation mutants of EGFP and the exceptionally stable superfolder GFP (sfGFP) ^12^ to determine to what extent we can shorten their termini without compromising fluorescence. To this end, we expressed truncation mutants in U2OS cells and assessed their fluorescence both by fluorescence-activated cell sorting (FACS) and by spinning disk confocal fluorescence microscopy imaging. Both FACS and imaging confirmed the sensitivity of EGFP to terminal truncations, and showed, at the same time, that sfGFP missing either the entire N-(ΔN12) or C-terminus (ΔC11), or both N- and C-termini (ΔN12ΔC11), retains largely its fluorescence and is usable for fluorescence imaging (Figure S1C,D).

We hypothesized that engaging both GFP termini could also constrain GFP mobility. Intramolecular GFP fusions, with GFP placed within a protein structure, for example within a loop, or between a transmembrane and extracellular domain, could reduce its rotational mobility. Such constructs have notably been used to constrain GFP mobility in fusions with the integral membrane proteins integrin ^13^ and the major histocompatibility complex class I (MHC I) protein ^14^. Alternatively, circularly permuted

GFP (cpGFP) ^15–17^ with the original termini connected via an ABD (Figure 2A) could also act to constrain GFP mobility. Given that better-folded variants of GFP also behave much better in intramolecular fusions in terms of functionality ^18,19^ and that they can be beneficial for the folding and stability of circular permutants ^20^, we considered the use of sfGFP as the best choice for GFP fusion engineering. To suppress GFP dimerization-related artifacts, we further introduced the V206K mutation to generated monomeric sfGFP (msfGFP) ^21–23^ which we used for all subsequent screening (Figure S1B). (use part of the former in main text)

Besides the use of full-length msfGFP for fusions with ABDs, we employed two additional approaches for intramolecular G-actin fusions. Instead of the full-length GFP, we used the 16-residue GFP fragment, β11, that we complemented with co-expressed GFP1-10 in a bipartite split-GFP complementation assay-like manner ^24^ (Figure 6A). We finally employed the tetracysteine-biarsenical system, which uses among the smallest genetically encoded tags for fluorescent labeling and which resembles conceptually the bifunctional rhodamine approach (Figure 6A). To this end, we genetically fused a short peptide sequence containing 4 cysteines to G-actin. Upon addition of the membrane-permeable, nonfluorescent biarsenical dye FlAsH (fluorescein arsenical hairpin binder) and its specific binding to the tetracysteine motif, the dye becomes fluorescent *in situ* in live cells ^25,26^. Both the increased functionality due to the small size of the genetic tag ^27^ and the rigidity of the peptide-fluorophore complex suggested by nuclear magnetic resonance (NMR) ^28^ prompted us to generate such fusions for G-actin. Terminal fusions of G-actin with this approach incorporate into actin filaments in mammalian cells ^25^ though their functionality was not tested.

Taking together the above-mentioned points, we embarked on a screen of GFP fusions to the widely used ABDs (Figure 1E), namely Lifeact, Utrophin Utr1-261, F-tractin9-52, Affimer6, as well as to human non-muscle beta G-actin. To constrain GFP mobility, we generated terminal, intramolecular and circularly permuted GFP fusions without linker sequences, with shortened GFP termini and shortened termini of the ABD.

Our results from the screening of all fusions informed us on the following with respect to the mechanisms of FP immobilization. First, shortening of the N- and C-termini in terminal fusions was the most efficient way to constrain GFP mobility without compromising its fluorescence. Circularly permuted sfGFP fusions could constrain GFP mobility, but not nearly as efficiently as terminal fusions because cpGFP fusions tolerate poorly the shortening of the GFP termini or/and the shortening of the ABD: fluorescence or/and F-actin binding are rapidly compromised upon such shortening because of the flexibility required to connect the original termini of cpGFP. Second, removing only the linker between any full-length ABD and full-length GFP did not have any effect on GFP mobility: additional shortening of the ABD or/and the GFP was systematically required (Figure 2B-F). Third, the removal of at least seven to nine residues from the N-terminus of GFP or ten residues from its C-terminus, was necessary to start constraining its mobility. Strikingly, the removal of an additional single residue, either from the ABD or the GFP terminus, was often required to reduce ψ angle values by several tens of degrees (Figure 2C L42 vs. L45; Figure 2D F5 vs. F11; Figure 2E U13 vs. U20; Figure 2F Af6 vs. Af7). Importantly, shortening of the GFP termini could compromise F-actin binding if the fused ABD terminus was also shortened at the same time: two residues between the end of the GFP barrel and the ABD were typically needed so as not to compromise F-actin binding. Finally, FP immobilization ultimately depends on the flexibility of the terminus or insertion site (e.g., loop) of the actin-binding moiety to which the FP is fused: if the used terminus or insertion site is inherently flexible, even the shortest FP will remain mobile. The latter was the case, for example, for all C-terminal GFP fusions of Utr261 (Figure S3A,B).

#### Engineering actin filament organization reporters using Lifeact

The N-terminal 17 amino acids of the budding yeast actin-binding protein Abp140 are known as the actin-binding peptide “Lifeact” ^29^ and are used widely for labeling F-actin in live cells. The original reporter used a C-terminal EGFP ^29^ but N-terminal EGFP fusions have also been used ^30^. To constrain GFP mobility in fusions with Lifeact, we generated C-terminal fusions (Figure S2A, L1 to L37), N-terminal fusions (Figure S2A, L38 to L53), as well as circular permutants with Lifeact connecting the original N- and C-termini (Figure S2A, L54 to L72). To remove flexible stretches, we did not include linkers in the fusions, and we further shortened the N- or/and the C-termini of GFP. At the onset of this study, the crystal structure of Lifeact bound to F-actin (Figure S1F,G and ^31,32^ was not yet known. Circular dichroism and NMR spectroscopies had suggested that Lifeact forms a helix between residues 2 and 10 ^29^, though a helix from position 3 to 17, i.e. essentially encompassing the entire Lifeact sequence, was most commonly found with the secondary structure prediction server Robetta (predictions not shown). These findings, the fact that the very C-terminus of Lifeact is not conserved among budding yeast strains (Figure S1E and ^29^), and our quest for the minimal actin-binding stretch of Lifeact, prompted us to also shorten Lifeact on either or both its termini, aiming at removing flexible stretches that are not essential for actin binding.

We screened the generated fusions with respect to their fluorescent levels, their capacity to bind F-actin, their localization, and the extent to which GFP is constrained based on polarimetry measurements (Figure S2A,B). Extensive shortening of the msfGFP termini did not affect fluorescence of terminal fusions, in line with the results of our truncation screen. The fluorescence of cpGFP fusions, on the other hand, was expected to depend on the length and composition of the linker connecting the original termini. Earlier studies suggested that linkers comprising at least 20 residues are needed to connect the GFP barrel ends, i.e. in between the end of β11 and the beginning of β1 strands, to allow for stable cpGFP folding and fluorescence, and this using flexible glycine-rich linkers ^15–17^. Not surprisingly, fluorescence was severely compromised or absent for cpGFP fusions when using 11-16 residue-linkers.

The capacity of the fusions to bind F-actin, using SF labeling as a readout, depended, as expected, on the length of Lifeact. Shortening of Lifeact on either or both termini when fused to full-length msfGFP showed that (a) Val3 is essential for binding F-actin, (b) Gly2 is not needed *per se* for binding F-actin but that it contributes to the latter, and (c) the six C-terminal residues are also not essential for F-actin binding; these residues are also the least conserved ones among budding yeast strains (Figures S2A and S1E-J). These results are fully in line with the Lifeact-F-actin structure and the mutagenesis results in the respective reports ^31,32^ that show the essential character of the first 11 residues of Lifeact. Importantly, the proximity of either terminus of Lifeact to the terminus of GFP, notably when combining shortened Lifeact with shortened GFP termini, was critical for F-actin binding. This was particularly evident in cpGFP fusions with shortened GFP termini, whereby F-actin binding was compromised despite the presence of full-length Lifeact, suggesting that a minimum of flexibility is required on either side of the actin-binding moiety to allow for F-actin binding. This latter effect proved eventually to be the bottleneck for constraining efficiently GFP in cpGFP fusions given that even moderate shortening of the GFP termini compromised F-actin binding.

Under our low-level expression conditions (see methods for promoter details), all fusions labeled all types of SFs, notably dorsal, ventral, peripheral, perinuclear actin cap and arc SFs, including focal adhesions (FAs), as well as mitochondrial actin (Figure S1K). To exclude that the additional localization of fusions to arc nodes reflected GFP dimerization-related artifacts (Figure S1L), we compared fusions bearing one, two or all three GFP monomerizing mutations (Figure S1L); they all localized to arc nodes excluding such a scenario.

Terminal GFP fusions proved to be the most efficient way to constrain GFP mobility (Figures 2B,C and S2A,B). Polarimetry measurements showed that removing only the linker between full-length Lifeact and full-length GFP did not have any effect on GFP mobility (Figure S2A,B). The additional removal of seven residues from the N-terminus of GFP or ten residues from its C-terminus, was necessary to start constraining its mobility. Combining shorter GFP termini with shorter Lifeact termini also proved determinant, notably in N-terminal Lifeact fusions whereby the removal of a single residue, Gly2, from Lifeact, reduced ψ angle values by several tens of degrees (compare L42 with L45 in Figure S2A,B). Altogether, polarimetry measurements showed that we succeeded to immobilize GFP both in N- and C-terminal fusions with Lifeact. We chose to focus on fusions L22 (LifeactΔC4-msfGFPΔN7) and L45 (msfGFPΔC9-LifeactΔN2) as the best performing reporters for further functional characterization and F-actin organization measurements in live cells and tissues.

#### Engineering actin filament organization reporters using the Utrophin Calponin Homology Domain

The N-terminal 261 amino acids of human utrophin contain an F-actin-binding calponin homology domain known as Utr-CH or Utr261 ^33^, which is widely used for visualizing F-actin in live cells and tissues. The original GFP fusion is N-terminal to the Utr-CH domain ^33^ but C-terminal EGFP fusions have also been used successfully ^34^. To constrain GFP mobility in fusions with Utr261, we generated N-terminal fusions (Figure S3A, U1 to U24), C-terminal fusions (Figure S3A, U25 to U42), as well as circular permutants with Utr261 connecting the original N- and C-termini (Figure S3A, U43 to U63). We did not include linkers in the fusions, and we further shortened the N- or/and the C-termini of GFP. The structure of the utrophin calponin homology domain bound to F-actin ^31^ was not available at the time of the beginning of this study, but the biochemical and structural data concerning the N-terminal residues 28-261 of human utrophin ^35,36^ provided already key insights. Although the N-terminal 27 residues of utrophin maximize its affinity for F-actin, they are dispensable for F-actin binding ^35^. The susceptibility of these same residues to degradation further suggested that they are not part of a compact structure and might be inherently flexible ^35^. This observation aligns with their partial disorder in their complex with F-actin ^31^. A truncation mutant of Utr261, Utr230-EN ^37^, was also shown to bind cytoplasmic actin filaments. These findings prompted us to also shorten Utr261 on either or both termini.

We screened the generated utrophin fusions along the same lines as for the Lifeact fusions (Figures S3B-G). All constructs were fluorescent except the three cpGFP fusions with the shortest N- and C-termini of GFP, suggesting again that a minimum of flexibility is needed to allow connecting the original N- and C-termini of GFP while binding to F-actin. Shortening of Utr261 on either or both termini in terminal fusions with full-length msfGFP confirmed that the N-terminal 27 residues of Utr261 are indeed dispensable for binding F-actin, and further showed that the C-terminal 31 or 39 residues are also not required for binding F-actin (Figures S3A-G). Importantly, the proximity of residues 29-32 in the N-terminal helix of Utr261 to the C-terminus of GFP, notably when combining shortened Utr261 with shortened GFP termini, was critical for F-actin binding (Figures S2A and S2E), in line with the structure of the UtrCH-F-actin complex ^31^. The localization of all fusions was similar to the one of Lifeact fusions (Figure S3C-F).

Terminal GFP fusions proved to be the most efficient way to constrain GFP mobility (Figures 2E and S3A,B). As expected from the results of the Lifeact fusions, removing only the linker between full-length Utr261 and full-length GFP did not have any effect on GFP mobility. The additional removal of nine residues from the N-terminus of GFP and of the N-terminal 27 residues of Utr261 were necessary to start constraining its mobility. It is noteworthy that the additional removal of a tenth residue from the GFP C-terminus was sufficient to reduce ψ angle values by several tens of degrees (compare U13 with U20 in Figure 2E). Interestingly, all C-terminal GFP fusions, including combinations of the shortest C-terminus of Utr261 and the shortest N-terminus of GFP, and the recently reported construct UG7 ^38^, were flexible, reflecting most likely inherent flexibility in the very C-terminus of Utr261. We decided to focus on fusion U20 (msfGFPΔC10-Utr28-222) as the best performing reporter for further functional characterization and F-actin organization measurements in live cells and tissues.

#### Engineering actin filament organization reporters using F-tractin

The N-terminal residues 9-52 of the rat enzyme inositol triphosphate 3-kinase A (ITPKA), also known as F-tractin-P (P for prototype), were shown to contain an F-actin-binding domain, a C-terminal GFP fusion of which is widely used as a reporter of F-actin localization in live cells ^39,40^ (Figure 2D and S4A). A slightly shorter peptide, N9-40, has been shown to retain F-actin binding, was given the name F-tractin ^40^, and is used interchangeably with F-tractin-P for visualizing F-actin in live cells (John Hammer, National Institutes of Health, Bethesda, MD, USA, personal correspondence). To constrain GFP mobility in fusions with F-tractin and F-tractin-P, we generated C-terminal fusions (Figure S4C, F1 to F22), N-terminal fusions (Figure S4C, F23 to F27), as well as circular permutants with the F-tractin peptide connecting the original N- and C-termini (Figure S4C, F28 to F33). We did not include linkers in the fusions, and we further shortened the N- or/and the C-termini of GFP. The structure of F-tractin or F-tractin-P, alone or in complex with F-actin, has not been solved to date. Secondary structure prediction of F-tractin-P using the program JPred3 in the original article suggested that F-actin binding resides in a putative α-helix comprising residues ∼30-40 ^39^. Secondary structure prediction using multiple programs, including JPred4, PHD, Phyre2, RaptorX and AlphaFold, suggests that the glycine- and proline-rich N-terminal ∼30 residues are unstructured, with residues ∼30-50 predicted to form a helix (Figure S4B). These predictions prompted us to also shorten F-tractin on either of its termini.

We screened the generated F-tractin fusions similarly to the Lifeact and utrophin fusions (Figures S4C-E). Unlike the previous screens, all constructs were now fluorescent, including cpGFP fusions with the shortest N- and C-termini of GFP, reflecting most likely the fact that the N-terminus of F-tractin is unstructured and does not bind F-actin thus imposing less constraints for connecting the original termini of cpGFP. Shortening of F-tractin and combinations thereof with shortened GFP termini showed (a) that the N-terminal residues 9-14 are dispensable for F-actin binding and (b) that residues 37-40 are critical for F-actin binding (Figure S4C and S4E). All fusions localized similarly to Lifeact and utrophin fusions (Figure S4E).

C-terminal GFP fusions of F-tractin proved to be the most efficient way to constrain GFP mobility (Figures 2D and S4C,D). The additional removal of seven residues from the N-terminus of GFP was necessary to constrain its mobility. In line with our results from the Lifeact and utrophin screens, it was very striking that the additional removal of a single residue from the C-terminus of F-tractin was sufficient to reduce ψ angle values by several tens of degrees (compare F5 with F11 in Figure 2D). Interestingly, all C-terminal GFP fusions of F-tractin-P, including combinations with the shortest N-termini of GFP, were flexible; we observed the same behavior for all N-terminal fusions of F-tractin, including combinations with the shortest C-termini of GFP. We interpret both observations as reflecting the inherent flexibility of the respective F-tractin termini. The fusion F11 (F-tractinN9-39-msfGFPΔN7) turned out to be the best performing reporter for F-actin organization measurements in live cells.

#### Engineering actin filament organization reporters using Affimer6

Affimers, originally named Adhirons ^41^, are synthetic, non-antibody-based protein binders that can be engineered to bind specific proteins of interest with high affinity and specificity. Among the recently developed Affimers is the F-actin-binding Affimer, Affimer6 ^42^, an N-terminal GFP fusion of which can be used to monitor F-actin localization in live cells. To constrain GFP mobility in fusions with Affimer6, we generated C-terminal fusions (Figure S4F, Af1 to Af11), N-terminal fusions (Figure S4F, Af12 to Af16), as well as circular permutants with the Affimer scaffold connecting the original N- and C-termini (Figure S4F, Af17 to Af25). We did not include linkers in the fusions, and we further shortened the N- or/and the C-termini of GFP. The structure of the Affimer scaffold revealed a very compact fold, with hardly any flexible unstructured residues at its termini ^41^ (Figure S4I). Thus, we only attempted to shorten its C-terminus to remove potentially flexible residues that could contribute to GFP mobility in C-terminal GFP fusions. F-actin binding is not expected to be affected since the actin-binding loops are far from the C-terminus ^41^.

We screened the generated Affimer6 fusions in a similar manner to the other ABD fusions (Figure S4F-I). All constructs were fluorescent apart from the three cpGFP fusions with the shortest N- and C-termini of GFP: the highly compact structure of the Affimer scaffold does most likely not provide the flexibility needed to connect the original N- and C-termini of GFP in these cpGFP fusions. F-actin binding was compromised only for shortened Affimer C-termini combined with highly shortened GFP termini; it is possible that the GFP in these fusions adopts a position that interferes with the actin-binding loops. All Affimer6 fusions localized in an indistinguishable manner from the other ABD fusions (Figure S4H).

Fusing full-length Affimer6 C-terminally to the shortest N-terminus of GFP proved to be the most efficient way to constrain GFP mobility (Figures 2F and S4F,G). It was again remarkable that the presence of an additional single residue at the N-terminus of GFP was sufficient to increase ψ angle values by several tens of degrees (compare Af6 with Af7 in Figure S4F,G). N-terminal GFP fusions of Affimer6 with C-terminally truncated GFP also constrained GFP mobility, but much less efficiently, as was the recently reported Affimer6-based construct POLArISact ^43^. We decided to use fusion Af7 (Affimer6-msfGFPΔN12) as the best performing reporter for further functional characterization and F-actin organization measurements in live cells and tissues.

#### Engineering red FP-based actin filament organization reporters

The fact that robustly folding GFPs tolerate much more extensive terminal truncations without losing fluorescence (Figure S1C,D) prompted us to undertake a similar approach for red FPs. Thus, we compared side by side the sensitivity of the widely used red FP, mApple ^22,44^, and of superfolder Cherry 2 (sfCherry2) ^45^ to terminal truncations. Both FACS and imaging corroborated our results with green FPs: whereas mApple is sensitive to N-terminal truncations, sfCherry2 tolerates missing its entire N- (ΔN12) or C-terminus (ΔC4) (Figure S7A,B), making the latter the best choice for constraining mobility in its fusions to actin-binding domains.

We decided to generate selectively sfCherry2-based terminal fusions for Lifeact and Affimer6. The length and composition of the termini of sfCherry2 are not the same as for GFP necessitating a minimum of screening, but the results from our previous screens helped narrow down our efforts to a limited set of constructs. F-actin binding was, as expected, impaired by the proximity of shortened FP termini to shortened Lifeact. Indeed, the best performing constructs were Af30 (Affimer6-sfCherry2ΔN12) and L81 (sfCherry2ΔC4-Lifeact) combining full-length Affimer6 and Lifeact with the most extensively truncated sfCherry2 termini (Figure S7E,F).

#### Engineering actin filament organization reporters using G-actin

N-terminal GFP fusions of G-actin, with a flexible linker in between the GFP and the G-actin, are widely used for monitoring actin localization in live cells and tissues. Early studies using such fusions to *Dictyostelium discoideum* actin ^46^, to human non-muscle beta G-actin ^47^ and to the *Drosophila* non-muscle G-actin Act5C ^48^ showed that such fusions are able to copolymerize with G-actin and recapitulate endogenous actin distribution as assessed by phalloidin stainings; the choice of an appropriate promoter to keep GFP-actin levels low was shown to be critical for minimal perturbation ^47^. An important finding was that C-terminal tagging of G-actin, even with tags as small as a hexahistidine tag or a dodecapeptide, impairs its incorporation into actin filaments ^49,50^. *Drosophila* expressing such C-terminally tagged G-actin have flight muscle with virtually no detectable sarcomeric organization and are flightless, but N-terminal fusions with the same tags restore sarcomeric organization and flight capacity ^50^. Thus, to constrain GFP mobility in fusions with human non-muscle beta G-actin, we generated exclusively N-terminal fusions. We did not include linkers in these fusions, and we additionally shortened the C-termini of GFP (Figure S7G, A1 to A5). We also generated an N-terminal fusion with a tetracysteine peptide (Figure S7G, A6).

As an alternative to N-terminal fusions and to maximize our chances to constrain GFP mobility, we also considered generating intramolecular GFP fusions. ^51^ succeeded in generating a fully functional intramolecular mCherry fusion of the bacterial actin homolog MreB by inserting mCherry right before helix 7 of MreB ^52^. Very interestingly, a second study by ^49^ generated an intramolecular fusion of fission yeast actin by inserting a dodecapeptide-based tetracysteine tag into Ser233-Ser234 of the loop following helix 7 (h7, hereafter), which was not functional but incorporated into actin patches upon FlAsH labeling. Motivated by these studies, we chose to engineer intramolecular fusions by inserting either full-length GFP (Figure S7G, A7 to A23), the GFP strand β11 (Figure S7G, A24 to A37), or a tetracysteine peptide (Figure S7G,

A38 to A47), either before or after h7 (Figure 6B). We used the exact same insertion site after h7 as in the study by ^49^. Fluorescence imaging showed that only fusions into the loop following h7 incorporated into actin filaments (A8, A25, A40 and A41 in Figure S7G). We thus focused on this insertion site for subsequent screening.

Motivated by the functionality of the intramolecular human beta- and gamma-actin GFP fusions using the same insertion site as for construct A8 (see main text and Figure 6D-R), and in order to constrain fluorophore mobility, we generated constructs that did not include linkers and where the N- and C-termini of GFP were shortened (Figure S7G). Three out of four serines within our insertion site are highly conserved across actin sequences (Figure 6B,C): we also generated constructs with differences in the exact number and position of these serines to establish possible effects on fluorophore mobility.

We screened the generated G-actin fusions with regard to their fluorescence, their capacity to integrate into F-actin, their localization and the extent to which GFP or FlAsH is constrained based on polarimetry measurements (Figures 6S and S7G-I). All terminal and intramolecular fusions with GFP and tetracysteine peptides were fluorescent. However, the only intramolecular β11-based fusion that was fluorescent was the one that included linkers (fusion A25 in Figure S7G): we reasoned that the absence of linkers and further shortening in the subsequent β11-based fusions did not provide the flexibility needed for the complementation of β11 with GFP1-10 ^24^. The absence of linkers and the additional shortening of the GFP C-terminus in terminal fusions did not compromise incorporation into actin filaments, but seemed to enrich less these fusions in myosin-II containing SFs (Figure S7G,I). Myosin-II interacts with the actin N-terminus providing a possible explanation for the latter observation. The absence of linkers in intramolecular GFP fusions did not compromise copolymerizing with actin, either. F-actin binding upon additional shortening of both GFP N- and C-termini in intramolecular fusions depended on the extent of shortening, as well as the exact number and position of the serine residues encompassing the insertion site. As expected, the use of a truncated CMV promoter for low-level expression was critical, with the widely used full-strength CMV promoter leading systematically to aggregation (Figure S7I). All G-actin fusions localized similarly to the other ABD-GFP fusions (Figure S7I). Interestingly, intramolecular tetracysteine peptide fusions labeled additionally nuclear F-actin (A41 in Figure S7I), showing nuclear F-actin bundles morphologically very similar to ones detected with the use of a nuclear actin chromobody ^53^ and an actin-NLS-FLAG construct ^54,55^. The small size of these peptides compared to the size of GFP could possibly explain this difference: GFP fusions were typically excluded from the nucleus, consistent with such an explanation.

Terminal and intramolecular GFP fusions without linkers and with extensively shortened GFP termini were most efficient to constrain GFP mobility (Figures S7G,H). The fusions A4 (msfGFPΔC10-actin) and A18 (actin-h7-msfGFPΔN7ΔC11-SSSSactin) are the best performing ones for F-actin organization measurements in live cells.

